# Guanine-nucleotide Exchange Modulator, GIV/Girdin, Serves as a Tunable Valve for Growth Factor-Stimulated Cyclic AMP Signals

**DOI:** 10.1101/149781

**Authors:** Michael Getz, Lee Swanson, Debashish Sahoo, Pradipta Ghosh, Padmini Rangamani

## Abstract

Cellular levels of the versatile second messenger, cyclic-(c)AMP are regulated by the antagonistic actions of the canonical G protein→adenylyl cyclase pathway that is initiated by G-protein-coupled receptors (GPCRs) and by phosphodiesterases (PDEs); dysregulated cAMP signaling drives many diseases, including cancers. Recently, an alternative paradigm for cAMP signaling has emerged, in which growth factor-receptor tyrosine kinases (RTKs; *e.g*., EGFR) access and modulate G proteins via cytosolic guanine-nucleotide exchange modulator (GEM), GIV/Girdin; dysregulation of this pathway is frequently encountered in cancers. Here we present a comprehensive network-based compartmental model for the paradigm of GEM-dependent signaling that reveals unforeseen crosstalk and network dynamics between upstream events and the various feedback-loops that fine-tune the GEM action of GIV, and captures the experimentally determined dynamics of cAMP. The model also reveals that GIV acts a tunable control-valve within the RTK→cAMP pathway; hence, it modulates cAMP via mechanisms distinct from the two most-often targeted classes of cAMP modulators, GPCRs and PDEs.

## 1 Introduction

Cells constantly sense cues from their external environments and relay them to the interior; sensing and relaying signals from cell-surface receptors involves second messengers such as cyclic nucleotides [1, 2]. Of the various cyclic nucleotides, the first to be identified was cyclic adenosine 3,5-monophosphate (cAMP), a universal second messenger that is used by diverse forms of life, from the unicellular bacteria, to fungi, protozoans and mammals. cAMP relays signals triggered by hormones, ion channels, and neurotransmitters [3], and also binds and regulates other cAMP-binding proteins such as PKA and Epac1 [4].

Intracellular levels of cAMP are regulated by the antagonistic action of two classes of enzymes: adenylyl cyclases (ACs) and cyclic nucleotide phosphodiesterases (PDEs). ACs are membrane-bound enzymes that utilize ATP to generate cAMP; the latter transmits signals from cell-surface receptors to second messengers. PDEs, on the other hand, are soluble enzymes, and catalyze the degradation of the phosphodiester bond resulting in the conversion of cAMP to AMP. PDEs are activated by protein kinase A (PKA), a downstream effector of cAMP, resulting in a negative feedback loop between cAMP and PDEs [5-8]. Thus, the level of cAMP in cells is a fine balance between synthesis by ACs, degradation by PDEs, and feedback through the PKA-PDE loop [3]. Both ACs and PDEs are also subject to positive and negative regulation by numerous other signaling pathways [9-11], which coordinately maintain cAMP levels in normal cells within a finite physiologic range. Dysregulated circuits that give rise to too much or too little cAMP can be unhealthy, and many disease states in humans are characterized by signaling programs driven by abnormal levels of cellular cAMP [see legend, **Figure 1A**]. For example, in the context of cancers, multiple studies across different cancers (*e.g*., breast [12], melanoma [13], pancreas [14], *etc*.) agree that high levels of cAMP are generally protective, whereas low cAMP levels fuel cancer progression [reviewed in [15]]. High cAMP inhibits several harmful phenotypes of tumor cells such as proliferation, invasion, stemness, and chemoresistance, while enhancing differentiation and apoptosis [see legend, **Figure 1A**]. Therapies that target the canonical GPCR/G-protein-cAMP signaling pathway have been successfully translated to the clinics, and they account for 40% of currently marketed drugs that can treat a wide range of ailments [16], from hypertension to glaucoma. However, such strategies have largely failed to impact cancer care or outcome. Thus, how tumor cells avoid high levels of cAMP appears to be incompletely understood, and therapeutic strategies to elevate cAMP remain unrealized.

**Figure 1:**
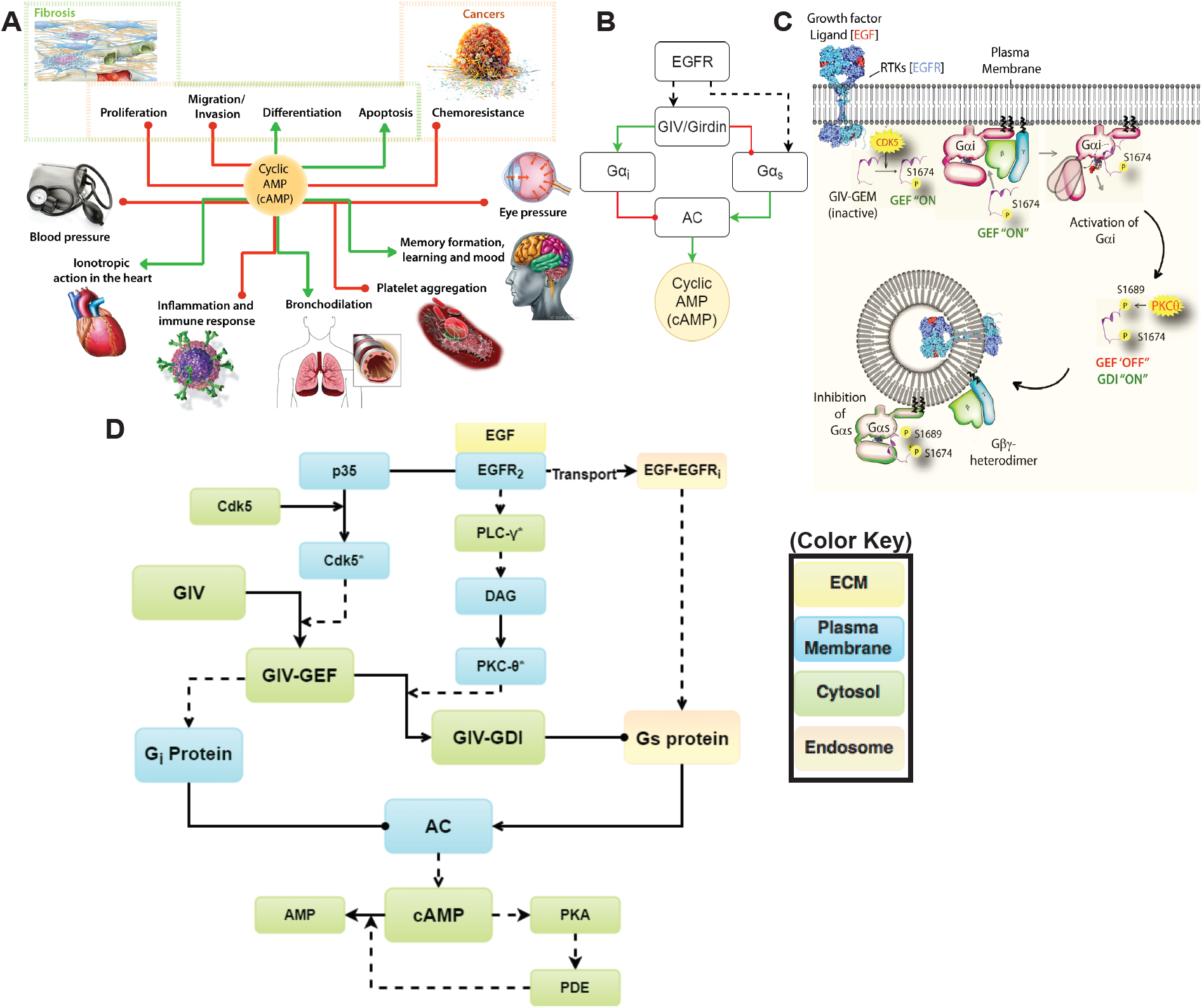
An emerging paradigm for modulation of cellular cAMP by growth factors. (**A**) Schematic summarizing the role of cyclic AMP (cAMP) in diverse biological processes. In cancers (top right), cAMP is largely protective as it inhibits proliferation, invasion, and chemoresistance, and promotes apoptosis and differentiation of tumor cells. Similarly, in the context of organ fibrosis, cAMP is a potent anti-fibrotic agent because it inhibits proliferation and migration and triggers apoptosis and return to quiescence for myofibroblasts, the major cell type implicated in fibrogenic disorders. Red lines indicate suppression and green lines indicate promotion. (**B**) A simplified circuit diagram of the non-canonical G protein→cAMP→PKA axis that is initiated by EGFR via GIV-GEM’s action on G*α_i_* (inhibits AC) and G*α_s_* (activates AC). Red lines indicate suppression and green lines indicate promotion. Interrupted black arrow = Activation of G*α_s_* occurs downstream of EGFR via unknown mechanisms. (**C**) A schematic showing the spatial features of G*α_i_* and G*α_s_* modulation downstream of EGFR, based on previously published work [17, 22, 28]. When the ligand binds the receptor, GIV is recruited to EGFR, and GIV-dependent signaling is initiated at the PM through the activation of CDK5. A single phosphorylation at S1674 activates GIV’s GEF function, which allows GIV-GEF to activate G*α_i_* in the vicinity of ligand-activated EGFR. GIV’s GEF activity towards G*α_i_* is subsequently terminated, and its GDI activity towards G*α_i_* is turned ‘ON’ by the phosphorylation at S1689 by PKC-*θ*. PKC-*θ* is activated by active EGFR through the PLC-*γ*→DAG→PKC-*θ* axis (not shown). Dually phosphorylated GIV (S1674 and S1689) binds and inhibits G*α_s_* signaling on endosomes. (**D**) A reaction network model showing the different signaling nodes and connections from EGFR to the cAMP→PKA signaling axis. Solid lines indicate a binding interaction; interrupted lines indicate enzymatic reaction. The color key (right, boxed) denotes the different compartments in which the components reside.

Recently, the regulation of cAMP by non-canonical G protein signaling that is initiated by growth factors [17-20] has emerged as a new signaling paradigm. Growth factor signaling is a major form of signal transduction in eukaryotes, and dysregulated growth factor signaling (*e.g*., copy number variations or activating mutations in RTKs, increased growth factor production/concentration) is also often encountered in advanced tumors and is frequently targeted with varying degrees of success [21]. A body of work published by us and others have revealed that RTKs bind and activate trimeric G proteins via a family of proteins called Guanine-nucleotide Exchange Modulators (GEMs; [17]). While the members of this family share very little sequence homology, and act within diverse signaling cascades, what unites them is the ability to couple activation of these cascades to G protein signaling via an evolutionarily conserved motif of approximately 30 amino acids that directly binds and modulates G*α_i_* and G*α_s_* proteins. Most importantly, GIV-GEM serves as a GEF for G*α_i_* and as a GDI for G*α_s_* [22], and despite this apparent paradox, both forms of modulation lead to suppression of cellular cyclic AMP [19]. By demonstrating how GIV, a prototypical member of a family of cytosolic guanine nucleotide exchange modulators (GEMs; [17, 22]), uses a SH2-like module [23] to directly bind cytoplasmic tails of ligand-activated RTKs such as EGFR [24] we provided a definitive structural basis for several decades of observations made by researchers that G-proteins can be coupled to and modulated by growth factors (reviewed in [25]). A series of studies that have followed since have revealed that growth factor-triggered non-canonical G protein→cAMP signaling through GIV has unique spatiotemporal properties and prolonged dynamics that are distinct from canonical GPCR-dependent signaling [reviewed in [18]]. More importantly, by straddling two major eukaryotic signaling hubs [RTKs and G proteins] that are most frequently targeted for their therapeutic significance, GIV has its own growing list of pathophysiologic importance as a therapeutic target in a variety of disease states, most prominently in cancers. High levels of GIV expression fuel multiple ominous properties of cancer cells, *e.g*., invasiveness, chemoresistance, stemness, survival, *etc*., and is associated with poorer outcome in multiple cancers, and inhibition of GIV’s G protein modulatory function has emerged as a plausible strategy to combat aggressive traits of cancers [26]; (reviewed in [17, 20]).

Despite these insights, the paradigm of RTK-dependent cAMP signaling remains nascent with many unknowns. Although the pathway may appear to be a linear connection between input (the growth factor RTKs) and output (G-proteins) elements, experimental data shows that non-canonical G protein→cAMP signaling via GIV-GEM integrates multiple input and output signals, with multiple feedback loops, and yet remains spatiotemporally segregated. It is initiated and terminated by specific phosphoevents, at specific times, on distinct membranes. However, signaling through this pathway can be disrupted by disassembling any one of the key signaling interfaces (reviewed in [20]). Therefore, the behavior of such complex systems is hard to grasp by intuition.

Here, we use systems biology approaches, namely mathematical and computational modeling, as the primary tools to generate a comprehensive model that can provide insights into some of these issues. This model, the first of its kind, not only connects two of the most widely studied eukaryotic signaling hubs [RTKs and G proteins], but also reveals surprising insights into the workings of GIV-GEM and provides a mechanistic and predictive framework for experimental design and clinical outcome.

## Results and discussion

### Construction and experimental validation of a compartmental model for non-canonical G-protein signaling triggered by growth factors

The emerging paradigm of non-canonical modulation of Gi/s proteins by growth factor RTKs is comprised of several temporally and spatially separated components (**Figure 1B, C**); each component is analyzed as distinct modules within a larger network model (**Figure 1D**). The model consists of four modules - **Module 1** focuses on the dynamics of EGFR (**Table S2**); **Module 2** represents the dynamics of the formation of the EGFR·GIV·G*α_i_* complex (**Table S3**; within this complex GIV-GEM serves as a GEF for G*α_i_*); **Module 3** represents the dynamics of the formation of the G*α_s_*·GIV-GDI complex (**Table S4**; within this complex GIV-GEM serves as a GDI for G*α_s_*); and **Module 4** represents the dynamics of cAMP formation (**Table S5**,**S6**).

Within each module, the biochemical reaction network includes several known interactions. For example, binding of EGF to EGFR at the plasma membrane (PM) initiates a cascade of events, which includes receptor dimerization, cross-phosphorylation of the cytoplasmic tails, recruitment of signaling adaptors, and phosphoactivation of a plethora of enzymes to relay downstream signaling. Of relevance to this paradigm, GIV, a multi-domain signal transducer contains a SH2-like domain in its C-terminus, which enables its recruitment to autophosphorylated cytoplasmic tails of EGFR [23]. We used this modular network to investigate the role played by GIV in regulating the dynamics of EGFR, EGFR·GIV·G*α_i_* complex, G*α_s_·*GIV-GDI complex, and cAMP.

### EGFR dynamics at the plasma membrane and the endosomal membrane

**Module 1** of the reaction network models the dynamics of EGFR at the PM and the endosome (**Figure 2A**, **Table S2**). At the PM, EGFR is activated by ligand binding, receptor dimerization, and cross-phosphorylation; activated EGFR is internalized to the early endosome through endocytosis, from where it can be either recycled or degraded [27]. Active PM EGFR also forms a complex with GIV-GEM, and via GIV with G*α_i_*, leading to the activation of G*α_i_* [22]. On the other hand, while it remains unclear when and where EGF/EGFR activates G*α_s_*, it is known that a pool of GIV that is on endosomes containing internalized EGFR binds and inactivates G*α_s_* on the endosomal membrane. Once inactivated, G*α_s_*-GDP enhances the degradation rate of internalized, endosomal EGFR, thereby limiting the pool of receptors available for recycling to the PM and serves to attenuate growth factor signaling [28].

**Figure 2:**
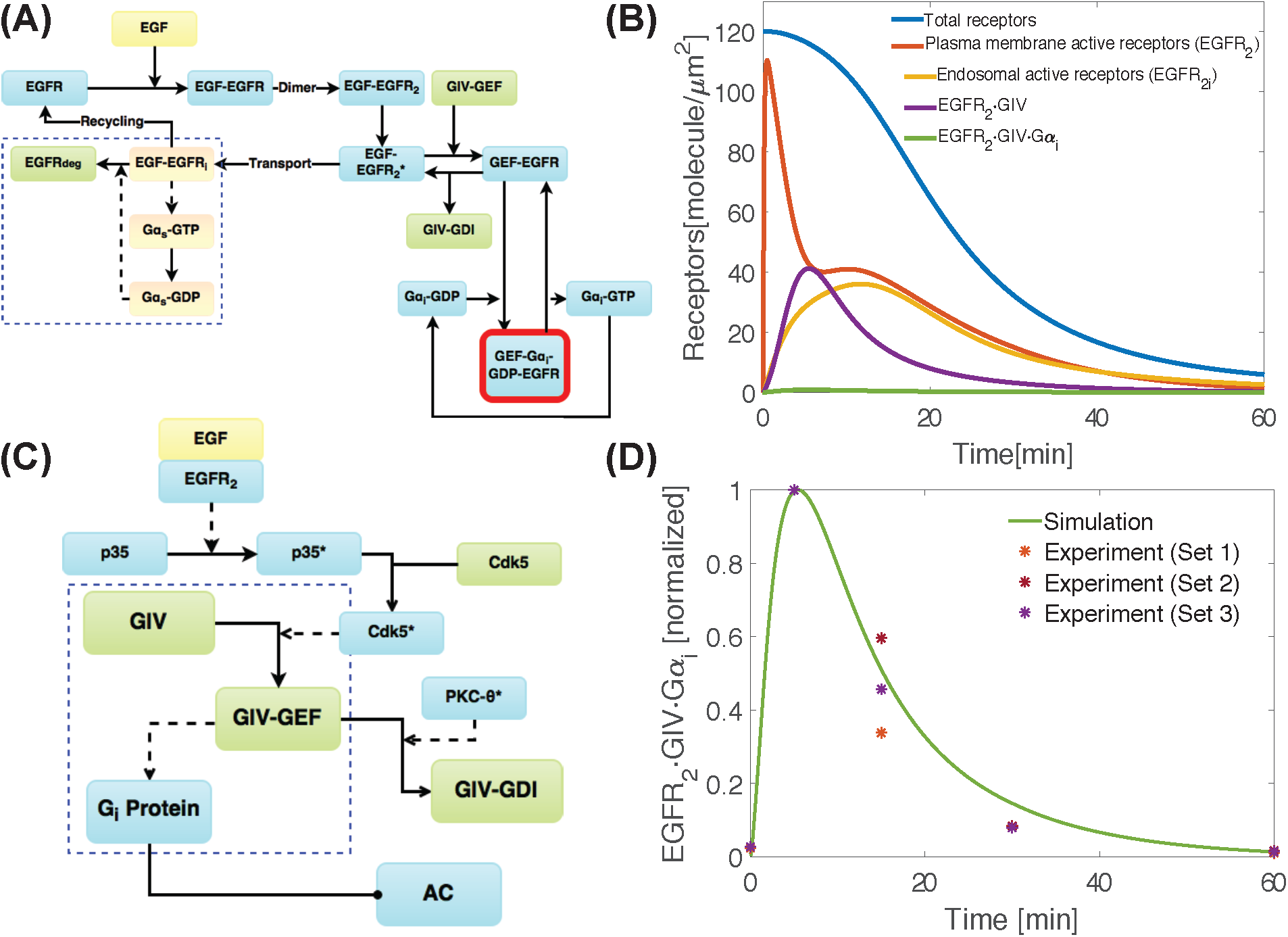
Early time scale events within the EGF/EGFR signaling cascade that are initiated at the plasma membrane drive G*α_i_* dynamics. (**A**) Network module 1 showing receptor interactions with feedback from G*α_s_*-GDP whose presence on endosomes accelerates receptor degradation due to rapid endosomal maturation [28]. (**B**) Graphs display the dynamics of different pools of EGFR over 1 hour as determined by simulations. EGFR dimer dynamics at the PM (red line), at the endosome (yellow line), bound to GIV-GEF (purple line), and in the EGFR·GIV·G*α_i_* complex (green line) are shown. The total number of dimerized receptors (blue line) decreases over time due to receptor degradation. (**C**) Module 2 showing the formation of EGFR·GIV·G*α_i_* ternary complex, a pre-requisite event necessary for activation of G*α_i_* during EGF signaling. (**D**) Simulations of dynamics of the formation of the EGFR·GIV·G*α_i_* complex based on network module in (**C**). The membrane density of the EGFR·GIV·G*α_i_* complex was normalized to its peak value and compared against experimental data (*) in which protein-protein interaction assays were performed using lysates of cells responding to EGF [Figure 1D and S1 of [22]]. The effect of different kinetic parameters on EGFR kinetics and the formation of the EGFR·GIV·G*α_i_* complex are shown in **Figure S1** and **S2** respectively.

Simulations from the model show that EGFR dynamics is governed by multiple time-scales when ligand stimulation triggers the redistribution of receptors from the PM to different pools (**Figure 2B**). The PM-pool of active receptors increases rapidly upon ligand stimulation (**Figure 2B**, red line) and subsequently recruits GIV, forming GIV-GEF-EGFR complexes (**Figure 2B**, purple line). The endosomal pool of active receptors increases at a slower time scale (**Figure 2B**, yellow line) than the PM-pool of active receptors. Recycling of the endosomal pool of receptors to the PM leads to a small second burst in the PM pool of receptors around 10 min (**Figure 2B**, yellow line). These findings are in agreement with Schoeberl et al. [27], indicating that our model accurately captures the EGFR dynamics. The total number of active receptors decreases over time because of G*α_s_*-GDP-dependent receptor degradation (**Figure 2B**, blue line). The pool of receptors in the GIV-GEF·EGFR complex subsequently interact with G*α_i_* at the PM to form the EGFR·GIV·G*α_i_* complex. The effect of kinetic parameters of EGFR dynamics is shown in **Figure S1** and we find that the balance of PM-pool versus internalized pool of EGFR is closely regulated by both the internalization rate and the G*α_s_*-GDP dependent receptor degradation rate [28].

### Dynamics of G*α_i_* signaling: activation kinetics are shaped by both upstream EGFR dynamics and downstream PLC-*γ*→DAG→PKC-*θ* signaling events

We next asked how EGFR dynamics affect the dynamics of G*α_i_* signaling at the PM. Activation of EGFR at the PM triggers a series of downstream events, including the activation of CDK5 at the PM by its cofactor, p35 [29]. CDK5 phosphorylates GIV at Ser(S)1675 and enhances GIV’s ability to bind G*α_i_*, *i.e*., CDK5 turns inactive GIV to into active GIV-GEF [30]. This allows GIV to couple G*α_i_* to EGFR by assembling ternary EGFR·GIV·G*α_i_* complexes at the PM [31] and activate G*α_i_* in the vicinity of ligand-activated EGFR (**Module 2** in the model, **Figure 2C**, **Table S2**, **S3**). EGFR also triggers the activation of the PLC-*γ*-DAG-PKC-*θ* pathway [32]; PKC-*θ* phosphorylates GIV at S1689 and terminates GIV GEF activity towards G*α_i_* [33]. Such sequential phosphorylation has another function - it converts GIV that is a GEF for G*α_i_* (GIV-GEF) into GIV that now serves as a GDI for G*α_s_*(GIV-GDI); GIV-GDI binds and inhibits GDP exchange on G*α_s_* [22].

We asked, how do the CDK5 and the PLC-*γ* pathways regulate dynamics of the EGFR·GIV·G*α_i_* complex formation, which is the key precursor event essential for transactivation of G*α_i_* by EGF/EGFR [23, 31]. Because the actual concentration of this complex in cells is not known, and is likely to vary from cell to cell, we analyzed peak times and normalized density of the EGFR·GIV·G*α_i_* complex formation. The temporal dynamics of these normalized densities of the EGFR·GIV·G*α_i_* complexes generated from simulations were in good agreement with experimental measurements, as determined by GST pulldown assays [22, 30] carried out in HeLa cells responding to EGF (**Figure 2D**).

Sensitivity analyses showed that despite the substantial number of model parameters (Tables **S10** and **S12**), the formation of the EGFR·GIV·G*α_i_* complex is sensitive only to a few kinetic parameters and initial conditions over time (**Tables S10** and **S12**, **Figure S2**). For example, a ten-fold variation of the forward rate for the binding of GIV-GEF to the activated receptor (*k_f_* in reaction 15, **Table S3**) affected the peak values of the complex formation (**Figure S2A**) but not the temporal features of the EGFR·GIV·G*α_i_* complex formation (**Figure S2B**). Similarly, the activation of the GIV-GEF function by CDK5 (reaction 14, **Table S3**) affected the density of the complex (**Figure S2C**) but not the temporal dynamics (**Figure S2D**).

The dynamics of the EGFR·GIV·G*α_i_* complexes, however, were sensitive to the initial concentrations of G*α_i_* (expected), GIV (expected), Cdk5 (expected), PIP_2_ (unexpected) and PLC-*γ* (unexpected) (**Table S10**). The sensitivity of EGFR·GIV·G*α_i_* complex formation to PIP_2_ and PLC-*γ* likely stems from network cross-talk, because the PLC-*γ*→DAG→PKC-*θ* pathway terminates GIV-GEF, triggering the dissociation of GIV and G*α_i_*, which triggers the disassembly of the EGFR·GIV·G*α_i_* complexes (**Figures 1C, 1D, 2A**). Changes in PLC-*γ* impacted both the density and temporal dynamics of the EGFR·GIV·G*α_i_* complexes. As expected, when the PLC-*γ*→DAG→PKC-*θ* pathway is inhibited, the lifetime of GIV-GEF is prolonged and *vice versa* (**Figure S2E**). This effect is evident when comparing the normalized densities against experiments (**Figure S2F**).

We conclude that early activation of GIV-GEF, and the observed dynamics of the assembly of EGFR·GIV·G*α_i_* complexes are not only dependent on the upstream kinetics of EGFR activation, but also on the downstream conversion of GIV-GEF to GIV-GDI, mediated by the PLC-*γ*→DAG→PKC-*θ* pathway. Findings also indicate that the connections within the network effectively capture the dynamics of transactivation of G*α_i_* by EGFR via GIV-GEF.

### Dynamics of G*α_s_* activation is most compatible with delayed activation triggered by internalized EGFR and inactivation by GIV-GEM on endosomes

Although GIV-GDI inhibits the activity of G*α_s_*-GTP [22], the exact mechanism of G*α_s_* activation by EGFR is currently unknown. Prior studies have shown that G*α_s_* is located on early, sorting and recycling endosomes [34] and that upon EGF stimulation, its activation/inactivation on endosomes regulates endosome maturation and EGFR degradation [35]; in cells without G*α_s_*, or in those expressing a constitutively active mutant G*α_s_*, internalized EGFR stays longer in endosomes, thereby, prolonging signaling from that compartment [28]. We asked when and where G*α_s_* is activated. Because compartmentalized EGFR signaling (PM versus endosomes) occurs on different time scales (**Figure 2B**), and G*α_i_* and G*α_s_* have different timescales of activation [5 min and 15 min respectively] [22], we reasoned that computationally predicted dynamics of all three possible scenarios of compartmentalized G*α_s_* activation *i.e*. [1) exclusively at the PM (**Figure 3A**, blue boxes); 2) exclusively at the endosomes (**Figure 3A**, yellow boxes); and 3) both at the PM and then on the endosomes (**Figure 3A**, both)], can provide insights into which option might be in accordance with the actual observed time scales for the same.

**Figure 3:**
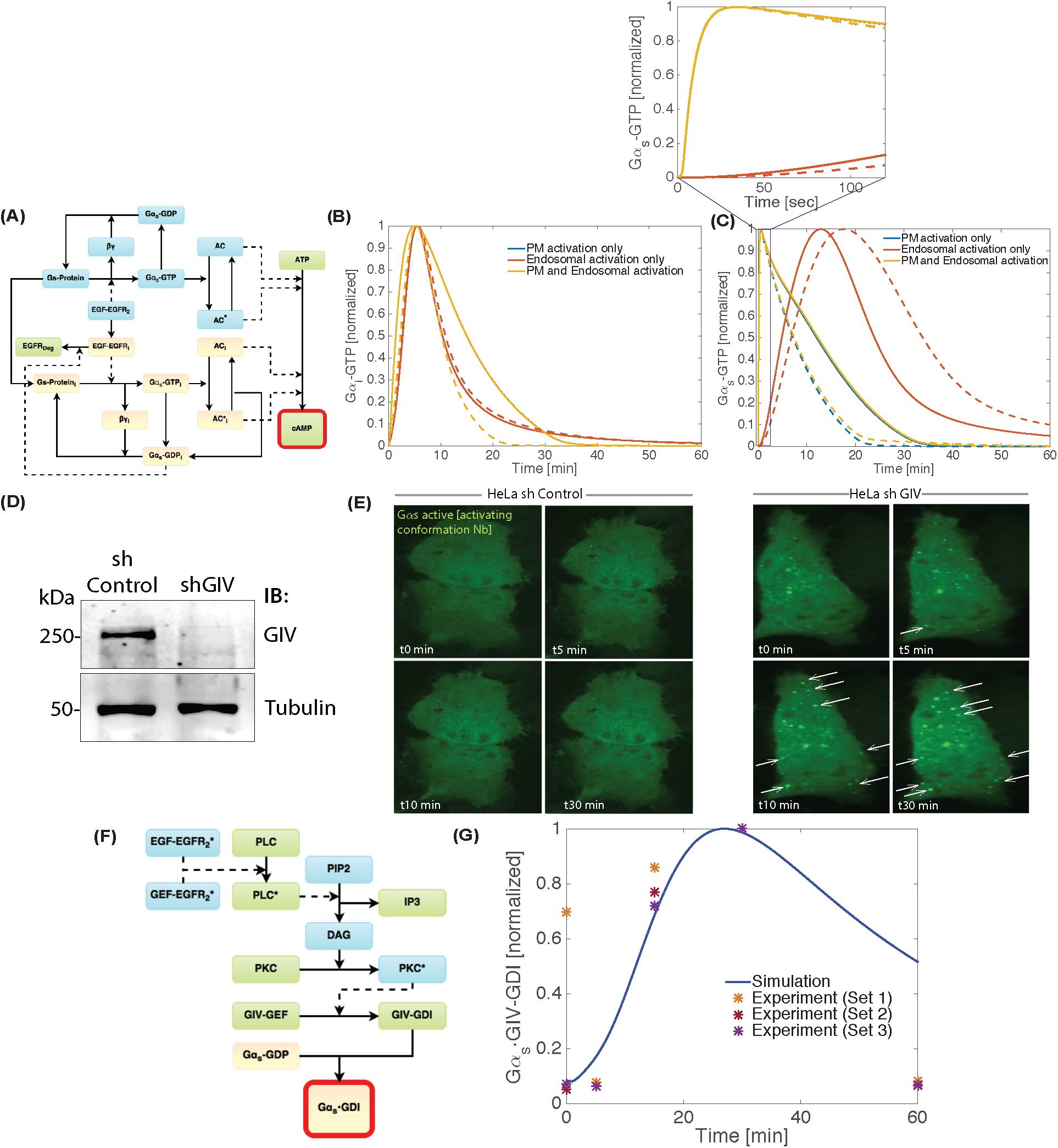
Late time scale events within the EGF/EGFR signaling cascade that are initiated on the endosome membrane drive G*α_s_* dynamics. (**A**) Network module showing the interactions of G*α_s_* and G*α_i_* with GIV at the PM and the endosome membrane. (**B**) Simulations conducted for the module shown in **A** shows that G*α_i_*-GTP dynamics at the PM are unaffected by the compartment in which G*α_s_*-GTP is activated. (**C**) Simulations conducted for the module shown in **A** comparing the dynamics of G*α_s_* activation in response to growth factor stimulation in 3 compartmental settings [see color key] and in the presence [solid lines] or absence [interrupted] of GIV. Activation of G*α_s_* at the PM alone is predicted to have a rapid activation and inactivation kinetics (see (**C**, zoom) for a 2 min zoom), while G*α_s_* activation on the endosome membranes is predicted to confer prolonged dynamics over longer time scales. In all cases, the presence/absence of GIV only impacts the prolonged phase, predicting higher G*α_s_* activation without GIV. (**D**) Equal aliquots of lysates of Hela cells used in (**E**) were analyzed for GIV depletion by immunoblotting(IB). Band densitometry confirmed > 95% depletion of GIV. (**E**) Freeze-frame images from live cell movies showing the dynamics of G*α_s_* activation in response to EGF, as determined by a biosensor that binds and helps detect the nucleotide-free intermediate during G*α_s_* activation [22]. Control (shControl) and GIV-depleted (shGIV) Hela cells expressing GFP-tagged anti-G*α_s_*·GTP conformational biosensor, nanobody Nb37-GFP were serum starved overnight and stimulated with 50 nM EGF and analyzed by live cell imaging using a Leica scanning disk microscope for 20 min. Freeze frames from representative cells are shown. In the presence of GIV (shControl) little or no G*α_s_* activity was seen after EGF stimulation; however, in GIV-depleted cells, G*α_s_* activity was seen on vesicular structures, likely to be endosomes (arrowheads; see Supplementary Movies 1-2). Bright puncta = active G*α_s_* on endocytic vesicles and/or endosomes. Bar = 10 *μM*. (**F**) Network module showing the interactions leading to the formation of G*α_s_*·GIV-GDI complex. (**G**) Dynamics of the formation of the G*α_s_*·GIV-GDI complex, a prerequisite event for inhibition of G*α_s_* by GIV, were simulated based on the network diagram shown in (**F**). The membrane density of this complex was normalized to its peak value and compared against experimental data (*) which is protein-protein interaction assays performed using lysates of cells responding to EGF [Figure 1D and S1 of [22]]. A good qualitative agreement was observed until 30 min, but not at 60 min, indicating that certain downstream regulators of G*α_s_* may be missing from the model (*e.g*. phosphatases) The effect of different kinetic parameters on the formation of the G*α_s_*·GIV-GDI complex are shown in **Figure S3**.

In the first scenario, where ligand-activated EGFR triggers G*α_s_* activation exclusively at the PM, activation is predicted to be rapid with peak concentration at 35 sec, similar to the case of *β*2-adrenergic receptors peak activity at 15 sec [36] (also see **Figure 7A**); this kinetic pattern mimics the dynamics of rapid EGFR activation at the PM [compare blue line in the zoomed in region of **Figure 3C** (new) with red line in **Figure 2B**]. In the second scenario, where ligand-activated EGFR triggers G*α_s_* activation exclusively on endosomal membranes, the time of peak activity is around 15 min (**Figure 3C**, red line), in accordance with the time scales of G*α_s_* activation and cAMP production [22]. Finally, if we consider a scenario where ligand-activated EGFR triggers G*α_s_* activation both at the PM and on endosomes, we observe a first peak of rapid activation at around 35 sec, followed by a second burst at around 15 min. In all three scenarios, activation of G*α_s_* [concentration of G*α_s_*-GTP] was higher in the absence of GIV’s GDI activity (i.e., when the concentration of GIV is set to zero; **Figure 3C**). Based on the dynamics of EGFR at the PM [rapid, almost instantaneous] and on the endosome [approximately 10 min] (**Figure 2B**) and similar timescales for G*α_s_* activation observed from the different modes of G*α_s_* activation (**Figure 3C**), we predict that G*α_s_* is likely activated on endosomes. It is noteworthy that the spatiotemporal features of such non-canonical G*α_s_* signaling does not affect the kinetics of G*α_i_* activation, either in the presence (**Figure 3B**, solid line) or absence of GIV (**Figure 3B**, interrupted line), indicating that EGFR transmodulates G*α_i_* and G*α_s_* independently.

To validate model predictions, we used a G*α_s_* conformational biosensor, nanobody Nb37-GFP that binds and helps detect the nucleotide-free intermediate during G*α_s_* activation [22]. Prior studies have extensively validated this tool and demonstrated its ability to detect G*α_s_* activation in real time, both at the PM (seen as a burst of signal in the cell periphery) and within early and recycling endoscomes (seen as dynamic punctate vesicles inside cells [37-40]. In control cells, no significant G*α_s_* activity was detected, neither at the PM, nor on endosomes, neither before, nor after ligand stimulation, indicating that G*α_s_* is either not activated after EGF stimulation or that its activity is efficiently suppressed by some modulator, presumably GIV, for sustained periods of time. In GIV-depleted cells [80-85% depletion of endogenous GIV by shRNA sequence targeting the 3’ UTR [22] **Figure 3D**], G*α_s_* activity was easily detected roughly 15 min after ligand stimulation and exclusively on vesicular structures, likely to be endosomes (**Figure 3E**); no such signal was noted at the PM, which is where canonical activation of G*α_s_* by GPCR is initiated [22]. These results obtained in live cells using conformation sensitive antibodies reveal a much delayed and compartmentalized pattern of non-canonical cyclical activation/inactivation of G*α_s_*downstream of EGF; findings are also consistent with our *in vitro* enzymology assays published previously [22] in that GIV’s GDI function normally inhibits G*α_s_* activity (hence not much fluorescence in control cells, but increased signals in GIV-depleted cells). As for what activates G*α_s_* downstream of EGF/EGFR, few studies have shown that EGFR binds G*α_s_* [41, 42] through its juxtamembrane region [43], and that this interaction triggers phosphoactivation of G*α_s_* [41]. Such transactivation of G*α_s_* by EGFR in cardiomyocytes is accompanied by augmented AC activation, elevation of cAMP, increased heart rate and contractility [41, 44]. Our model neither proves nor disproves this model for direct transactivation of G*α_s_* by EGFR, but reveals that activation of G*α_s_* is delayed and pinpoints endosomes as the site of such activation.

Finally, we evaluated the dynamics of formation of the G*α_s_*·GIV-GDI complex [**Module 2**; **Figure 3F**], the precursor event that is essential for transinhibition of G*α_s_* by EGF/EGFR [22]. Our model for the dynamics of assembly of G*α_s_*·GIV-GDI complexes (**Table S2**, **S4**, **S5**) included the kinetics of receptor internalization, G*α_s_* activation by internalized receptors, conversion of GIV-GEF to GIV-GDI by the PLC-*γ*→DAG→PKC-*θ* pathway, and the G*α_s_*-GDP-dependent degradation of endosomal EGFR (**Figure 2A**). Simulations from this model showed a good qualitative agreement between normalized G*α_s_*·GIV-GDI complex formation between model and cell-based experiments [22], particularly until 50 min after EGF stimulation (**Figure 3G**).

The role of kinetic parameters and initial conditions affecting the formation of the G*α_s_*·GIV-GDI complex were explored in detail (**Figure S3**) and we found that the dynamics of G*α_s_*·GIV-GDI complex formation is more sensitive to internalization and degradation of EGFR than to any other kinetic parameters. However, our model was unable to capture the precipitous reduction in the normalized concentrations of G*α_s_*·GIV-GDI complexes at 60 min. We speculate that the discrepancy between model and experiment may stem from the fact that the model is fine-tuned to compute the G*α_s_*·GIV-GDI complexes that are located exclusively on the endosomes, whereas the experiment assessed G*α_s_*·GIV-GDI complexes in whole cells (not restricted to the endosomes) by proximity ligand assays (PLA) on endogenous proteins or by GST pulldown assays using cell lysates as source of G*α_s_*. Experimentally, it is not yet possible to assess specifically the number of cytosolic versus endosomal vs other membrane-localized G*α_s_*·GIV-GDI complex numbers in living cells responding to EGF. Because G*α_s_* on endosomes escapes redistribution after ligand stimulation, it has a prolonged half-life to enable sustained signaling from that location [45]. It is possible that the endosomal pool of GIV-GDI has a similarly prolonged half-life, which could explain the unexpectedly high number of complexes predicted at 60 min.

Alternatively, the discrepancy may simply reflect an incompleteness in network modeling. For example, one plausible group of unknown proteins that are missing in our model are downstream phosphatases that presumably act on GIV-GDI on endosomes, and are responsible for the decline in the number of G*α_s_*·GIV-GDI complexes at later time points.

### Biological predictions from the model: compartmentalized modulation of G*α_i_* and G*α_s_* governs EGF-triggered cAMP dynamics

Because EGF/EGFR triggers activation of G*α_i_* at the PM first, followed by activation of G*α_s_* on the endosomes later, production of cAMP must be a balance between the antagonistic actions of these two G proteins on membrane-bound ACs (**Figure 4A**, **Table S5**). Because EGFR triggers G*α_i_* activation predominantly at the PM, we assumed that the PM-pool of G*α_i_* inhibits the PM AC→cAMP pathway at the PM. Similarly, because G*α_s_* is activated predominantly on endosomes and endosomal ACs [eACs] can be stimulated at that location to synthesize cAMP locally [46], we assumed that the endosomal-pool of G*α_s_* likely stimulates the eAC→cAMP pathway (**Table S5**). To capture the dynamics of cAMP in our model network, we included such compartmentalized G protein-AC interactions (**Figure 4A**).

**Figure 4:**
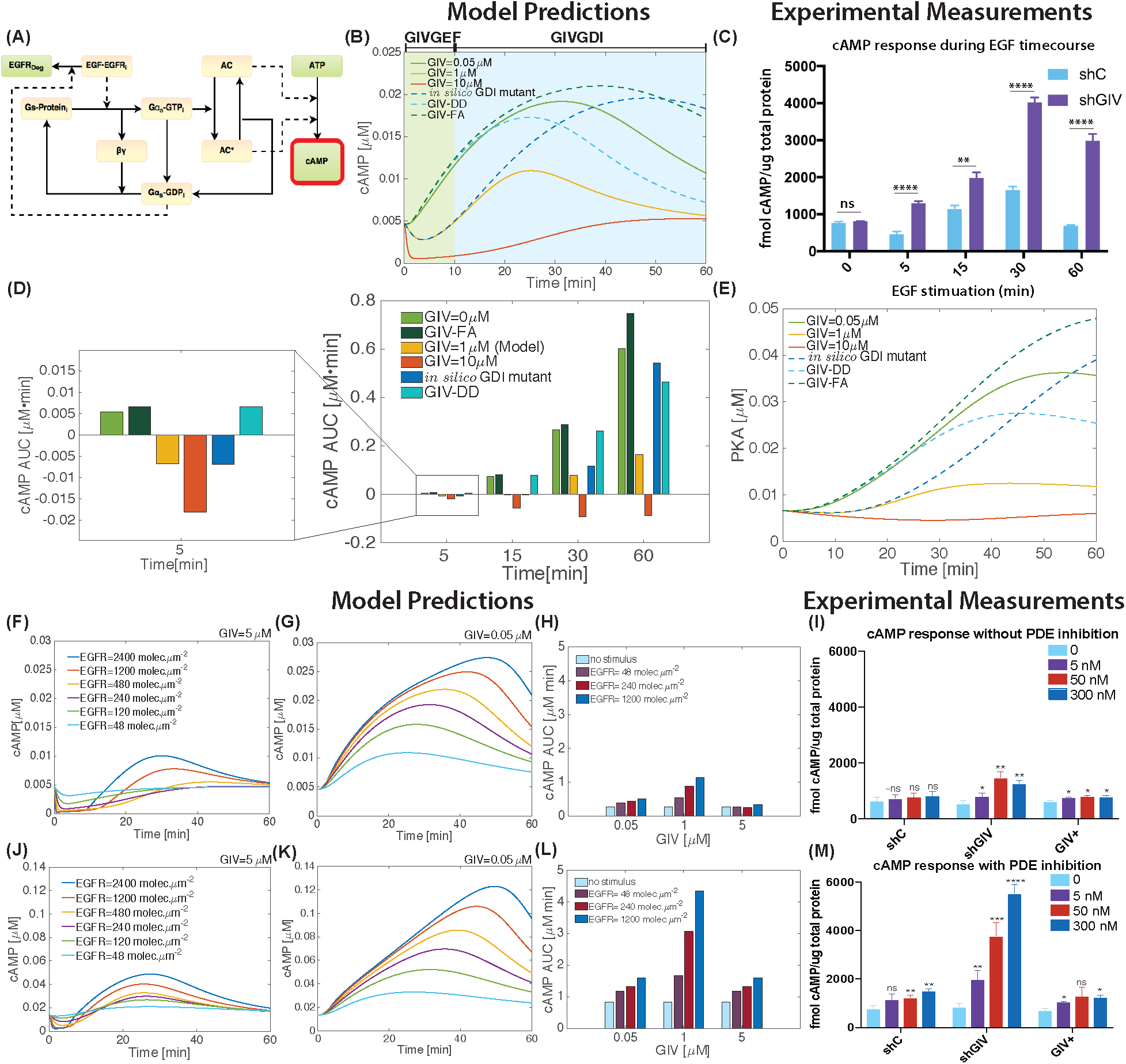
Dynamics of growth factor triggered cAMP signaling via GIV. (**A**) Network module of cAMP synthesis mediated by AC. (**B**) Simulations of cAMP dynamics in response to EGF based on module in **A** identify the two distinct regimes of GIV GEM’s effect on cAMP dynamics. The early 0-5 min phase (green region) is characterized by a dip in cAMP concentration; GIV’s GEF function on G*α_s_* dominates in this phase. This is followed by a delayed approximately 10-60 min phase (blue region), which is characterized by an increase in the concentration of cAMP; GIV’s GDI function on G*α_s_* dominates this phase. Simulations predict that cAMP dynamics will decrease with increasing GIV expression [compare the yellow line (control GIV in the model) with the red (high GIV) and the green (low GIV) lines]. In high GIV states (red line), the transition from GIV-GEF dominant (early, approximately 0-5 min) to GIV-GDI dominant (late, approximately 10-60 min) regimes is evident as the line transitions from negative to positive. Simulations for cAMP dynamics are also displayed for 3 other conditions: 1) GIV in the absence of its GEF effect on G*α_i_* (the GIV-DD mutant), in the absence of both its GEF and GDI effects (GIV-FA mutant), and in the absence of its GDI effect (in the presence of an in silico GDI-deficient mutant). The GIV-DD mutant (dashed cyan line) doesn’t show the initial decrease in cAMP concentration; the *in silico* GDI mutant (dot dashed blue line) shows an initial decrease in cAMP concentration but then a prolonged increase in cAMP; the GIV-FA (black dashed lines) shows cAMP dynamics same as that of low GIV (green line), i.e., no initial decrease and a prolonged increase in cAMP. (**C**) control or GIV-depleated (shGIV) HeLA cells were serum starved (0.2% FBD, 16h) prior to stimulation with 50nM EGF for the indicated time points. cAMP produced in response to EGF was measured by radioimmunoassay (RIA) as detailed in ‘Materials and method’. Bar graphs compare the cAMP levels in shC vs shGIV cells at each time point. Error bars indicated mean S.D of three independent experiments. ns= not significant; ^**^p=0.01,^****^p=0.0001. An overlay of the both the sh and shGIV cAMP experimental and computational data can be found in **Figure S5**.(**D**) The area under the curve (AUC) for cAMP dynamics was calculated for different time points after EGF stimulation. The transition from GIV-GEF dominant (early, 0-5 min) to GIV-GDI dominant (late, 10-60 min) regimes is evident in the transition from AUC going from negative to positive. The AUC remains negative at all times for GIV=10 *μΜ*, indicating that cAMP levels do not increase when GIV is high. The magnified image shows the AUC at 5 min. (**E**) PKA response upon EGFR stimulation at different GIV concentrations show that they follow a similar GIV-dependent trend to cAMP (see panel **C**). (**F**-**G**) Simulations comparing the impact of variable input signals [via EGF/EGFR] on cAMP dynamics in low GIV (**F**) and high GIV (**G**) states. When the concentration of GIV is set to 0.05μ M (**F**), increasing EGFR copy number results in increasing cAMP. When GIV concentration is set at high levels (5 *μΜ*; (**G**)), increasing EGFR copy number has little or no impact on cAMP concentrations. (**H**) AUCs calculated from **F** and **G** are displayed. cAMP AUC over time is pronounced for no-GIV state (0 *μΜ*; green bars); there is a very small change in the AUC over time for high-GIV state (5 *μΜ*; red bars). (**I**) Control (shC) or GIV-depleted (shGIV) HeLa cells or GIV-depleted cells rescued with shRNA-resistant GIV-WT (GIV+) were stimulated with EGF and assessed for cAMP levels exactly as in 4C with two exceptions. Measurements were carried out only at 60 min, and 3 different concentrations of EGF were tested as described. Bar graphs compare the cAMP levels in response to varying EGF concentrations. Error bars indicate mean ± S.D. of three independent experiments. ns= not significant *^*^p*=0.05; *^**^p*=0.01; ^***^*p*=0.001. (**J**-**K**) Same as (**F**-**G**) with reduced concentration of PDE to mimic inhibition. Simulations show that when PDE is inhibited, the impact of variable input signals [via EGF/EGFR] on cAMP dynamics in low GIV (**J**) and high GIV (**K**) states does not change even though the actual concentration of cAMP is increased. (**L**) AUC calculations show the higher cAMP concentrations when PDE is inhibited. cAMP AUC over time is pronounced for no-GIV state (0 *μΜ*; green bars); there is a very small change in the AUC over time for high-GIV state (5 *μΜ*; red bars). (**M**) Same as in **I**, with one additional step of pre-treatment of cells with 200 *μΜ* IBMX (20 min) prior to EGF stimulation. Error bars indicate mean ± S.D. of three independent experiments. *^*^p*=0.05; ^**^*p*=0.01; ^***^*p*=0.001; *^****^p*=0.0001. See **Figure S6** for the effect of GIV-GEM on cAMP peak time, **Figure S8** for the effect of receptor dynamics on cAMP, **Figure S9** for additional experimental data, and **Figure S10** for the effect of CDK5 and PKC-*θ* phosphorylation of PDE.

Our model predicted that the early inhibition of cAMP is due to the G*α_i_*-mediated inhibition of AC (the green regime) (**Figure 4B**); cAMP production is increased later due to the activation of G*α_s_* on the endosome (the blue regime) (**Figure 4B**). These dynamics are consistent with previously published GIV-dependent cAMP dynamics, measured by FRET [22]. While activation of GIV-GEF occurs earlier [within 5 min] at the PM, conversion of GIV-GEF to GIV-GDI occurs later [15-30 min] when EGFR is already compartmentalized in endosomes (**Figure 1B**); such temporally separated compartmentalized modulation of two G*_α_*-proteins with opposing effects on AC ensures suppression of cAMP at both early and later times during EGF signaling [22]. Because GIV modulates both G*α_i_* and G*α_s_* in different compartments and at different time scales, the model predicts that increasing GIV concentration should dampen overall cAMP response to EGF, and that decreasing GIV concentration should do the opposite (**Figure 4B**, compare GIV = 0.05 *μM* with GIV = 10 *μM*). These predictions were tested by measuring cAMP in control (shControl) and GIV-depleted (shGIV) cells at various time points after EGF stimulation (**Figure 4C**) by a radioimmunoassay (RIA). We found that compared to control cells, cellular levels of cAMP were always higher, both at early and late time points after EGF stimulation. In fact, when superimposed, the model and experiment showed good agreement throughout 60 min (**Figure S5**). As expected, sensitivity analyses confirmed that cAMP is sensitive to the initial concentrations of PDE, AC, G*α_i_*, and G*α_s_* and the reaction rates associated with AC, PKA, and PDE (**Table S11**, **S14**).

To dissect what might be the relative contributions of the two G protein modulatory functions of GIV [GEF versus GDI] on cAMP production, we investigated cAMP dynamics in three conditions (**Figure 4B**) – 1) GEF-deficient but GDI-proficient (mimicked experimentally by the GIV-S1764D/S1689D mutant, GIV-DD [22], 2) both GEF- and GDI-deficient (mimicked experimentally by GIV-F1685A mutant, GIV-FA [22, 47], and 3) GEF-proficient but GDI-deficient (an *in silico* mutant because there is no known mutant yet that can mimic this situation in experiments). In the first scenario, where GIV’s GEF function is selectively lost, but GDI function is preserved, increase in cAMP concentration occurred early (**Figure 4B**, dashed cyan line) as observed previously in cells expressing the GIV-DD mutant [22]. In the second scenario, where both GEF and GDI functions were lost, increase in cAMP concentration occurred early and such elevation was sustained (**Figure 4B**, dashed dark green line), as observed previously in cells expressing the GIV-FA mutant [22]; this mirrored the profile observed in GIV-depleted cells (**Figure 4B**, solid green line). Finally, in the third scenario, selective blocking of GIV’s GDI function using an *in silico* mutant resulted in an early decrease followed by a prolonged increase in cAMP concentration (**Figure 4B**, dot-dashed blue line).

While the dynamics of cAMP production provide insight into how different conditions lead to changes in concentration, the <u>a</u>rea <u>u</u>nder the <u>c</u>urve [AUC] for cAMP concentration provides information critical for decision-making, buffers from time scale variations, and averages the effect of fluctuations in concentrations [48]. AUCs for cAMP at different time points were calculated to investigate how the cumulative cAMP signal varies under different GIV conditions (**Figure 4D**). For the control bars (in orange), we observe that at the 5 min time point, the AUC is negative. This represents the initial decrease in cAMP concentration. The AUC becomes positive and increases by 15 min, signifying a net accumulation of cAMP. The AUCs look similar in the GIV-FA mutant [defective in both GDI and GEF functions] as well as in the absence of GIV, i.e., it increases progressively through 60 min (**Figure 4D**, compare the light green and dark green bars). If GIV levels are increased (10 *μM*, red bars) the AUCs remain negative throughout, showing the sustained nature of the dampening effect of GIV on cAMP. This dampening effect on cAMP is achieved primarily via activation of G*α_i_* in the short term [GEF regime] and via inhibition of G*α_s_* in the long term [GDI regime].

We next investigated the effect of cAMP dynamics on PKA. Activation of PKA by cAMP was modeled as a Hill function [49] and the fitting was compared against the experimental data as shown in **Figure S7**. The temporal dynamics of cAMP are reflected in the downstream dynamics of PKA and PDE (**Figure 4E**). The delay in PKA activation at short time scales [5 min] corresponds to the GIV-GEF mediated decrease in cAMP for control conditions (**Figure 4E**). As cAMP increases, PKA activity increases. In cases where there is no GIV-GEF mediated decrease in cAMP at the short time scale, there is no delay in PKA activation, resulting in an immediate and increased accumulation of PKA.

We also investigated how GIV concentration affects cAMP dynamics [output signal] with varying EGF/EGFR numbers [input signals]. When GIV concentrations were set to 0.05 *μM* in the model (to simulate cells that don’t have GIV), the network showed sensitivity, in that, increased input signals triggered increased output signals (**Figure 4F**); this effect was even more pronounced in the absence of PDE (**Figure 4J**). Such sensitivity was replaced by robustness when GIV concentrations were set to high levels (GIV = 5 *μM* **Figure 4G**), *i.e*., increased input signals failed to initiate proportional output signals; this effect was virtually unchanged and robustness was preserved despite the absence of PDE (**Figure 4K**). These effects are also evident by studying the AUC (**Figure 4H**, **L**).

To test these predictions, we measured cAMP by RIA in control and GIV-depleted HeLa cells as in **Figure 4C**, except in this instance we measured only at 60 min, but with varying doses of EGF [experimental equivalent of variable input in simulations]. To recapitulate simulations in the presence or absence of PDE, assays were carried out in parallel in the presence or absence of 3-isobutyl-1-methylxanthine (IBMX), an inhibitor of PDE (**Figure 4I**, **M**). In the presence of GIV, cAMP production is robustly suppressed in response to increasing EGF ligand (**Figure 4I**). In the absence of GIV, cAMP production is sensitive to increased EGF, an effect that is further accentuated when PDEs are inhibited with IBMX (**Figure 4M**). Taken together, these results indicate that GIV primarily serves as a dampener of cellular cAMP that is triggered downstream of EGF; unlike PDE, which reduces cellular cAMP by degrading it, GIV does so by fine-tuning its production by G proteins and membrane ACs.

## Clinical predictions from the model – from math to man

### Concurrent upregulation of both GIV and EGFR maximally reduces cAMP and carries poor prognosis in colorectal tumors

We next asked how cAMP levels that are triggered by growth factors and modulated via GIV can be related to tumor aggressiveness and clinical outcome. First, we conducted simulations for a wide range of GIV (0.05-5 *μΜ*) and EGFR (120 to 2400 EGFR *molecules/μm*^2^) concentrations to identify how crosstalk between these two variable components regulates cAMP levels (**Figure 5A-D**). Within each category of EGFR concentration [low or high], cAMP levels are the highest when GIV levels are lowest, and *vice versa*. In the setting of high EGFR expression (**Figure 5B**), the impact of changing GIV was the highest, *i.e*., the range of cAMP response was the widest. By contrast, in the setting of low EGFR expression (**Figure 5A**) the impact of changing GIV on the cAMP levels was minimal. Because the EGFR copy number and GIV copy number variables in our model are a surrogate for EGFR or GIV signaling states, respectively, which cumulatively represents all perturbations that can impact the functions of both EGFR and GIV [i.e., activating mutations, phosphomodifications, gene copy number variation, or simply protein overexpression in tumors], we conclude that the G*α_i_/*G*α_s_*/cAMP modulatory function of GIV exerts its maximal impact in the setting of high EGFR signaling states. These findings indicate that the two modules [EGFR and GIV-dependent G*α_s_*/G*α_i_*/cAMP signaling] are intertwined via a functional crosstalk.

**Figure 5:**
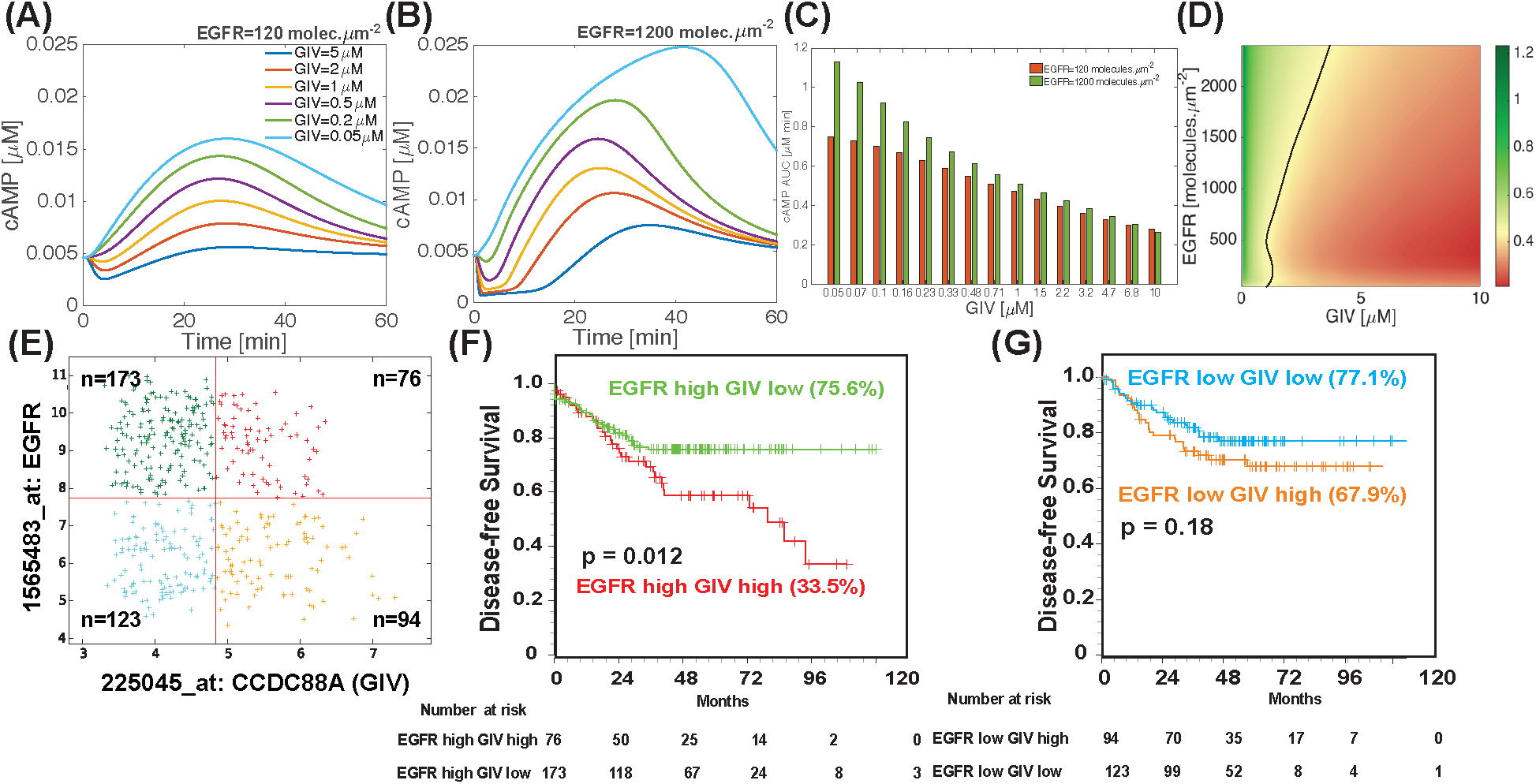
The impact of levels of expression of GIV and EGFR on cAMP dynamics; comparison of model predictions and clinical outcome [disease-free survival] in patients with colorectal cancers. (**A**-**B**) Simulations comparing the impact of variable GIV expression on cAMP dynamics in low EGFR (**A**) and high EGFR (**B**) states. When the concentration of EGFR is set at low levels (120 molecules*μm*^-2^; **A**), changing GIV copy number has very little impact on cAMP. When EGFR concentration is set at high levels (2400 molecules*μm*^-2^; **B**), changing GIV copy number has a larger impact on cAMP concentrations. Effect of additional combinations of GIV and EGFR concentrations on cAMP dynamics are shown in **Figure S11**. (**C**) Findings in **A**-**B** are displayed as bar graphs of area under the curve (AUC) for cAMP concentration, as integrated over 1 h. Red bars = low EGFR state; green bars = high EGFR state. (**D**) A heat map shows the area under the curve for cAMP concentration over 1 h for different concentrations of GIV [X-axis] and EGFR receptor [Y-axis]. This is a linear plot, with both the GIV axis and the EGFR axis increasing linearly. The black line, over the yellow region, corresponds to the control condition in the simulation. For any given level of EGFR expression, increasing GIV expression decreases the AUC for cAMP. For any given level of GIV expression, increasing EGFR expression increases the AUC for cAMP, but this effect is true only for lower concentrations of GIV. (**E**-**G**) GIV expression status in colon cancers has an impact on disease-free survival (DFS) when the level of expression of EGFR is high. Hegemon software was used to graph individual arrays according to the expression levels of EGFR and GIV (CCDC88A) in a data set containing 466 patients with colon cancer (see Methods; **E**). Survival analysis using Kaplan-Meier curves showed that among patients with high EGFR, concurrent expression of GIV at high levels carried significantly worse prognosis than those with low GIV (**F**). Survival analysis among patients with low EGFR showed that levels of expression of GIV did not have a significant impact on DFS (**G**).

In order to quantify the extent of this crosstalk, we conducted simulations for a wider range of EGFR [48-2400 *molecules/μm*^2^] and GIV concentrations [0.05-10 *μΜ*] and calculated the AUC for the cAMP dynamics (**Figure 5C, D**). Varying GIV concentrations resulted in cAMP changes within a narrow range in low EGFR state, but showed larger variance in the high EGFR state (**Figure 5C**). Because cAMP serves as a potent antagonist for almost all harmful properties in cancer cells (see **Figure 1A**), we hypothesized that chronic ‘high cAMP-state’ or ‘low GIV’ signature in cancers could serve as a good prognosticator and conversely, ‘low cAMP-state’ or ‘high GIV’ signature could serve as poor prognosticator. We posited that there could be regimes of operation in the EGFR-GIV space that could be exploited for prognostication in cancers. To further dissect this space, we plotted the variations in AUC for EGFR and GIV variations (**Figure 5D**). The value of AUC corresponding to the control (GIV *1μM* and EGFR 120 *molecules/μm*^2^, **Table S9**) is approximately 0.5 *μM·min* and is denoted by the yellow color and marked as a black solid line for different EGFR and GIV concentrations in the heat map; elevated cAMP level is denoted by green and reduced cAMP by red. We observed that increasing EGFR increased cAMP AUC for low GIV concentrations. But as GIV concentrations increased, even at high EGFR, the cAMP AUC decreased, indicating that the impact of increasing GIV on cAMP AUC was higher than the impact of increasing EGFR. Therefore, GIV levels, rather than EGFR copy number alone, can be thought of as a key determinant and a driver for chronically suppressed cAMP levels.

Having identified that GIV levels rather than EGFR levels alone are key to signaling in cells, we investigated if the EGFR-GIV axis played a role in clinical outcome as well. To determine the impact of crosstalk between EGFR and GIV on clinical outcome, we compared the mRNA expression levels to disease-free survival (DFS) in a data set of 466 patients with colorectal cancers (see Methods). Patients were stratified into negative (low) and positive (high) subgroups with regard to GIV (CCDC88A) and EGFR gene-expression levels with the use of the StepMiner algorithm, implemented within the Hegemon software (hierarchical exploration of gene-expression microarrays online; [50]) (**Figure 5E**). Kaplan-Meier analyses of DFS over time showed that among patients with high EGFR, expression of GIV at high levels [which favors low cAMP levels] carried a significantly poorer prognosis compared to those with low GIV [which favors increased cAMP levels] (**Figure 5F**), indicating that a low cAMP-signature state indeed carries poor prognosis. Among patients with low EGFR, expression of GIV at high or low levels did not impact survival (Figure 5G). Among patients with low EGFR, expression of GIV at high or low levels did not impact survival (**Figure 5G**). That the impact of EGFR-GIV interplay on patient survival are significant in a rigorous Kaplan-Meier analysis of a sufficiently large cohort of patients, despite numerous independent variables indicates that the interplay between EGFR and the G protein modulator, GIV is an important determinant of cancer progression. Findings are in agreement with the profiles of cAMP modulation by GIV downstream of EGFR, and the well-defined anti-cancer effect of cAMP.

Conversely, among patients with high levels of GIV, survival was significantly different between those with high versus low EGFR; no such trend was noted among patients with low GIV (**Figure 5F**). Thus, the high GIV/high EGFR signature carried a poorer prognosis compared to all other patients. More importantly, patients with tumors expressing high EGFR did as well as those expressing low EGFR provided the levels of GIV in those tumors was low. Consistent with the fact that cAMP is a potent anti-tumor second messenger, these findings reveal that 1) high levels of EGFR signaling does not, by itself, fuel aggressive traits or carry a poor prognosis, but does so when GIV levels are concurrently elevated; 2) in tumors with low GIV, the high EGFR signaling state may be critical for maintaining high cAMP levels and therefore, critical for dampening several aggressive tumor traits [**Figure 1A**].

The crosstalk between EGFR and GIV the we define here, and its impact on clinical outcome provide a plausible explanation for some long-standing conundrums in the field of oncology. Deregulated growth factor signaling (*e.g*., copy number variations or activating mutations in EGFR, increased growth factor production/concentration) is often encountered and targeted for therapy in advanced cancers [21]. Although activating EGFR mutations, copy number variations, and levels of EGFR protein expression seem to be closely related to each other [51], the prognostic impact of EGFR expression in cancers has been ambiguous [52]. In some cancers, high EGFR copy numbers are associated with poor outcome [53, 54]; in others, high EGFR expression unexpectedly favors better overall and progression-free survival [55-57]. Thus, due to reasons that are unclear, not all tumors with high EGF/EGFR signaling have an aggressive clinical course. Dysregulated GIV expression, on the other hand, is consistently associated with poorer outcome across a variety of cancers [19]. Our findings that GIV levels in tumors with high EGFR are a key determinant of the levels of the anti-tumor second messenger cAMP, have provided a potential molecular basis for why elevated EGFR signaling in some tumors can be a beneficial in some, but a driver of metastatic progression in others. Because cAMP levels in tumor cells and GIV levels have been previously implicated in anti-apoptotic signaling [58] and the development of chemoresistance [59], it is possible that the GIV-EGFR crosstalk we define here also determines how well patients may respond to anti-EGFR therapies, and who may be at highest risk for developing drug resistance. Whether such is the case, remains to be evaluated.

### Concurrent downregulation of both GIV and PDE activity maximally increases cAMP and carries a good prognosis in colorectal tumors

Another factor that plays an important role in regulating cAMP levels is PDE. We next conducted simulations for different GIV (0.05 to 1 *μM*) and PDE (0.04 to 2 *μM*) concentrations to identify how the crosstalk between these two variable components regulates cAMP levels. Within each category of PDE concentration [low vs high PDE states; (**Figure 6A-B**)], cAMP levels are the highest when GIV levels are lowest, and *vice versa*. In the setting of low PDE activity (**Figure 6B**), the impact of changing GIV was the highest, *i.e*., the range of cAMP response was the widest. By contrast, in the setting of high PDE activity (**Figure 6A**) the impact of changing GIV on the cAMP levels was minimal. These effects can be seen also when comparing the AUCs for the low vs high PDE states, calculated over 1 hr (**Figure 6C**). While there is no significant change in the AUC with increasing GIV in a high-PDE state (red bars), increase in GIV leads to a decrease in cAMP in PDE state (green bars). That is, for a given GIV concentration, the effect of PDE is always stronger. Furthermore, a heat map of cAMP AUCs (**Figure 6D**) shows the interplay between PDE and GIV concentrations over a wide range. For low PDE concentration, increasing GIV decreases cAMP AUC, but the cAMP AUC is well above the yellow value (marked as control). However, increase in PDE concentration leads to a dramatic decline in cAMP AUC even when GIV levels are low; this condition is likely to result in futile cycling [high cAMP production due to low GIV and high cAMP clearance due to high PDE signaling]. Together, these findings indicate that the effect of GIV concentration on cAMP levels in cells is discernible only when PDE activity is low. Because high PDE state virtually abolishes all effects of GIV-dependent inhibition of cAMP production, we also conclude that in this GIV-PDE crosstalk, PDE is a dominant node and GIV is the subordinate node. Because the ‘PDE copy number’ variable in our model is a surrogate for PDE activity/signaling, which cumulatively represents all perturbations that can impact the functions of PDEs in cells [i.e., phosphomodifications, mislocalization, or simply protein overexpression / hyperactivation in tumors; [60]], we conclude that the G*α_i_*/G*α_s_* modulatory function of GIV exerts its maximal impact on whether cAMP is high or low in the setting of low PDE signaling states.

**Figure 6:**
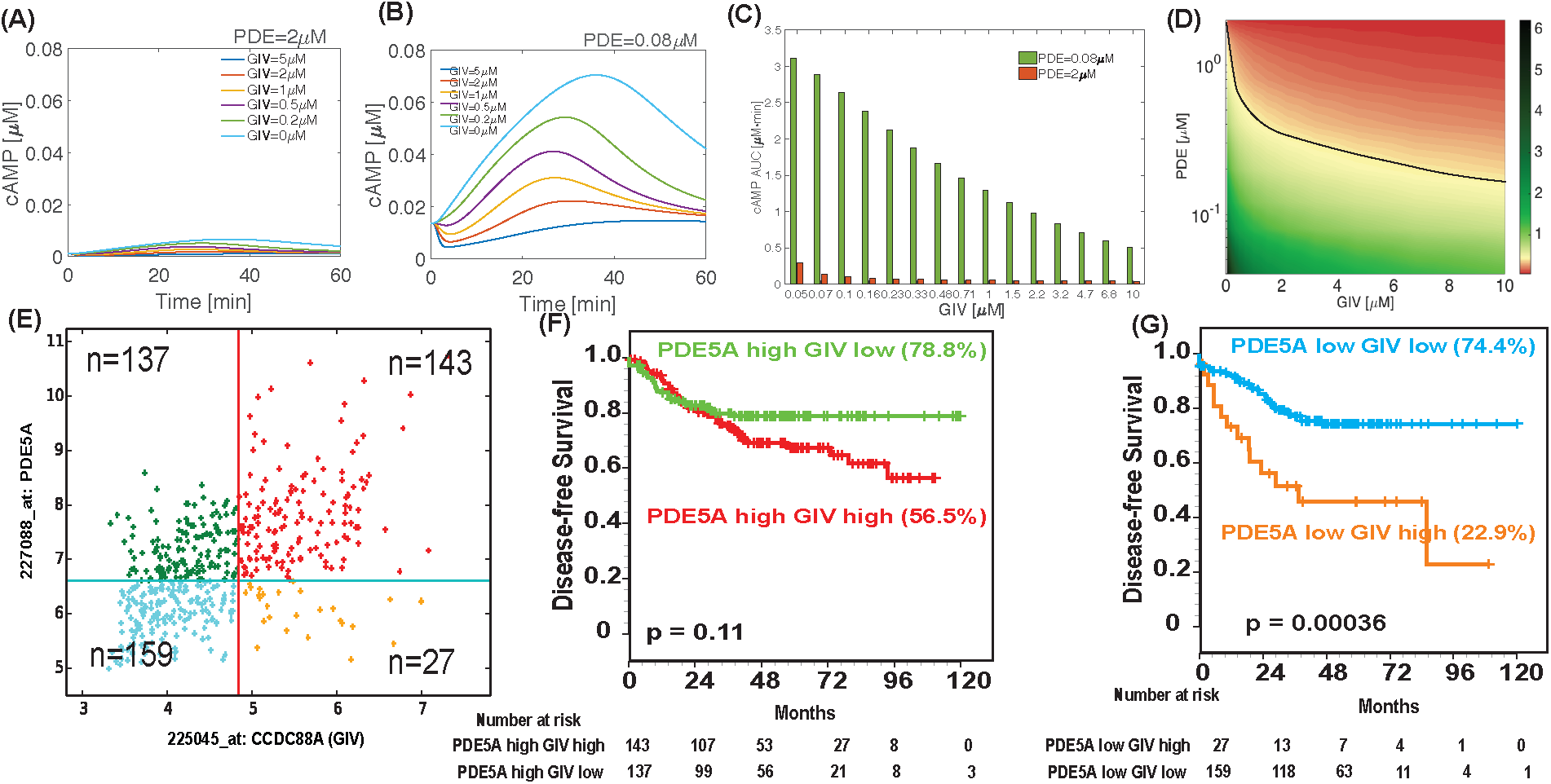
The impact of levels of expression of GIV and PDE on cAMP dynamics; comparison of model predictions and clinical outcome [disease-free survival] in patients with colorectal cancers. (**A**-**B**) Simulations comparing the impact of variable GIV expression on cAMP dynamics in high PDE (**A**) and low PDE (**B**) states. When the concentration of PDE is set at high levels (2 *μM*; **A**), changing GIV copy number has very little impact on cAMP. When PDE concentration is set at low levels (0.08 *μM*; **B**), changing GIV copy number has a larger impact on cAMP concentrations. Effect of additional combinations of GIV and PDE concentrations on cAMP dynamics are shown in **Figure S12**. (**C**) Findings in **A**-**B** are displayed as area under the curve (AUC) for cAMP concentration, as integrated over 1 h. Red bars = high PDE state; green bars = low PDE state. (**D**) A heat map shows the area under the curve for cAMP concentration over 1 h for different concentrations of GIV [X-axis] and activity levels of PDE [Y-axis]. This is a semi-log plot, with the PDE axis on the log scale and the GIV axis on the linear scale. The black line, over the yellow region, corresponds to the control condition in the simulation. As anticipated, the maximum amount of cAMP AUC is seen for low PDE and low GIV concentrations. For any given level of PDE activity, increasing GIV expression decreases the AUC for cAMP, but this effect is seen exclusively at low PDE activity states. For any given level of GIV expression, increasing PDE activity levels decreases the AUC for cAMP across all concentrations of GIV. (**E**-**G**) GIV expression status in colon cancers has an impact on disease-free survival (DFS) only when the level of expression of PDE5A are low. Hegemon software was used to graph individual arrays according to the expression levels of PDE5A and GIV (CCDC88A) in a data set containing 466 patients with colon cancer (see Methods; **E**). Survival analysis using Kaplan-Meier curves showed that among patients with high PDE5A, high vs low GIV expression did not carry any statistically significant difference in DFS **(F)**. Survival analysis among patients with low PDE5A showed that patients whose tumors had high levels of expression of GIV had a significantly shorter DFS than those with tumors expressing low levels of GIV **(G)**. See also **Figure S13** for patient survival curves for other PDE isoforms and GIV on DFS.

**Figure 7:**
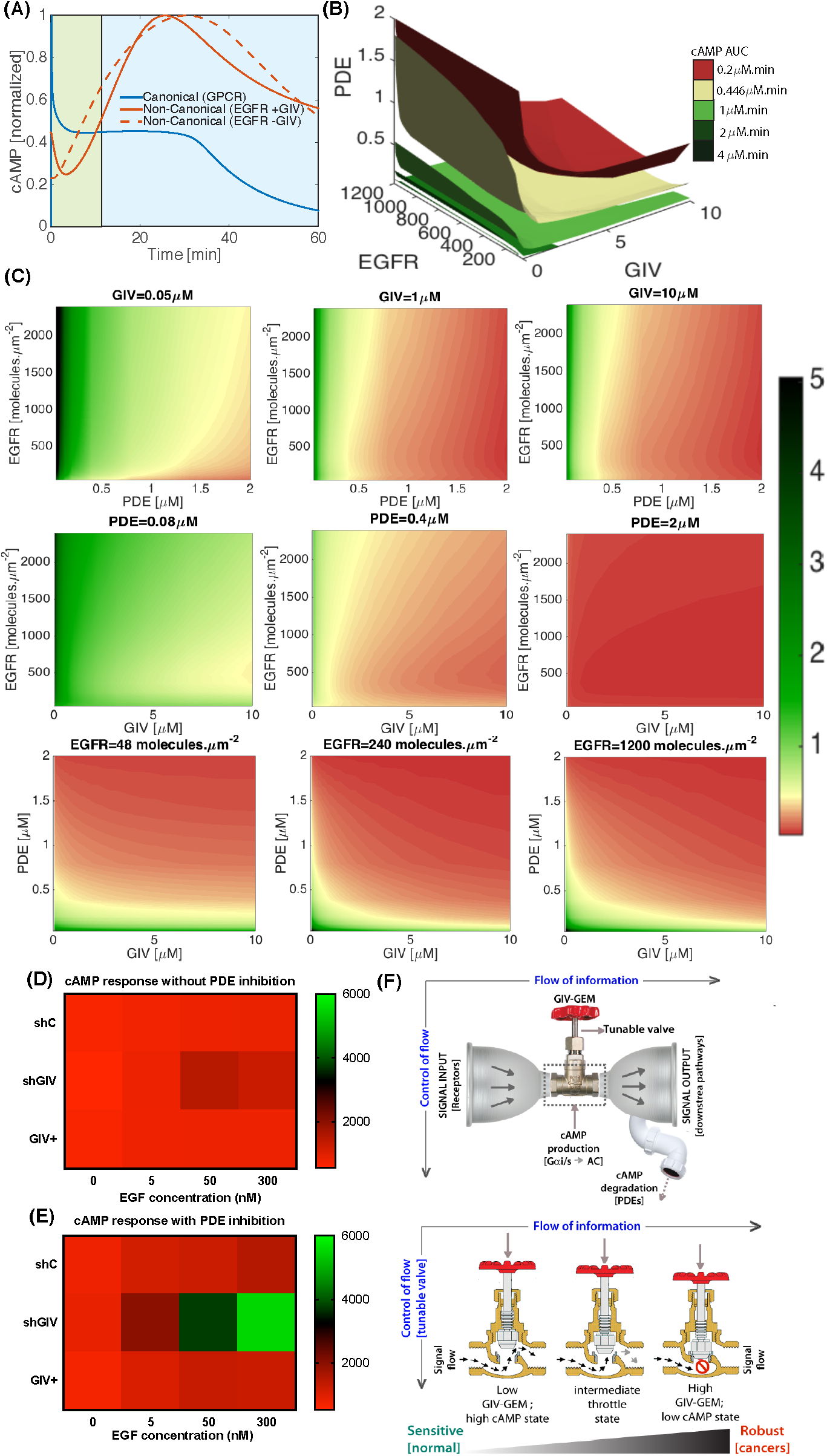
Compartmental modulation of G proteins by GIV allows it to serve as a tunable valve for growth factor stimulated cAMP production; low-GIV state imparts sensitiveness, whereas high-GIV state imparts robustness. (**A**) Simulations of cAMP dynamics that is initiated by the canonical GPCR-stimulated pathway (*β*2-adrenergic receptor stimulating G*α_s_*; blue line) and the non-canonical RTK-stimulated pathway that is modulated by GIV-GEM (red lines; solid = with GIV; interrupted = without GIV). In both cases, cAMP values (y axis) were normalized to the max value during a 6G min simulation. Canonical signaling is finite with a predominant PM phase, the non-canonical pathway features prolonged time scales due to a predominant endosomal phase. The interrupted line at approximately 5 min indicates the time period when ligand activated EGFR is typically rapidly endocytosed, marking a watershed between end of PM and beginning of endosomal phase of signaling. (**B**) A 4-D map showing the relationships between EGFR (input signal), GIV (control valve), and PDE (degradation sink) on cAMP dynamics (output signal). The different planes on this map correspond to the same value of cAMP AUC (see color key on right). The control value is shown in yellow (G.4436 *μM·min*). (**C**) Heatmaps showing AUCs for different planes from the 4-D map in **B**. (**C**, Top) Constant GIV planes: As GIV concentration increases, the cAMP AUC decreases for different values of EGFR and PDE; when GIV is low, cAMP AUC is sensitive to the amounts of EGFR and PDE (left panel). When GIV concentration increases [left to right], cAMP AUC loses sensitivity to EGFR input (right panel). (**C**, Middle) Constant PDE planes: When PDE is low, cAMP AUC is sensitive to the amounts of values of EGFR and GIV (left panel); when PDE activity increases [left to right], cAMP AUC loses sensitivity until no effect of EGFR or GIV is seen at highest concentrations (right panel). (**C**, Bottom) Constant EGFR planes: When EGFR is low, cAMP AUC is low; since PDE and GIV both suppress cAMP production (left panel). As EGFR concentration increases, the regions with high cAMP AUC increases showing a clear effect of increasing input (middle and left panels). (**D**, **E**) Heat maps generated using GraphPad PRISM for cAMP measurements performed in control (shC), GIV-depleted (shGIV) and GIV-depleted cells rescued with GIV-WT (GIV+) HeLa cells responding to varying EGF concentrations in **Figure 4I** (**F**; without PDE inhibition) and **4M** (**G**; with PDE inhibition). (**F**) Schematic summarizing the unique impacts of GIV-GEM on the EGFR→cAMP pathway, as revealed by systems biology. Top: Within the ‘bow-tie’ macroarchitecture of layered signal flow in any circuit, incoming signals from RTKs like EGFR and possibly other other RTKs known to signal via GIV[signal input; left] are integrated by core proteins like GIV-GEM [center] which modulate second messengers like cAMP, which subsequently impacts multiple target proteins such as kinases, phosphatases, and transcription factors [output signals; right]. Prior systems biology work had concluded that cellular concentrations of cAMP is a key determinant of robustness at the core of information (signal) flow [74-76]. While cAMP production is tuned up or down by variable levels of GIV and its compartmentalized action on G*α_i_*/G*α_s_* and ACs within the RTK→cAMP pathway, cAMP degradation by PDEs serves as a dominant sink [drain pipe]. Bottom: The inner workings within the knot of the bow-tie (rectangle inset) is depicted where vertical flow of ‘control’, up or down-regulation of GIV-GEM in cells serves as a tunable control valve, allowing cells to control cAMP production in cells responding to growth factors. When GIV-GEM expression is low [as seen in the normal epithelium on the left], increasing input signals can trigger some of the highest levels of cellular cAMP, thereby conferring sensitivity. Increasing GIV-GEM expression throttles the cAMP response [middle], such that, when GIV-GEM is expressed highly [as seen across all cancers; see supplementary figures **Figure S14**, **S15**], cAMP levels remain low, regardless of the amount of input signals, thereby conferring robustness [right].

As before, since GIV appeared to a control axis in determining the effect of PDE in cAMP in cells, we asked if the GIV-PDE axis played an important role in clinical data as well. To determine the impact of crosstalk between various PDE isoforms and GIV on clinical outcome, we carried out using the StepMiner algorithm, implemented within the Hegemon software on the same set of 466 patients with colorectal cancers as before, except patients were now stratified into low and high subgroups with regard to GIV (CCDC88A) and PDE gene-expression (**Figure 6E**). Among the 11 known PDE isoforms, we evaluated those that have previously been linked to colon cancer progression [PDE5A (**Figure 6E-G**), 4A and 10A (**Figure S12**)]. Kaplan-Meier analyses of DFS over time showed that although expression of GIV at high levels was associated with disease progression and poorer survival in both low and high PDE groups, the risk of progression was not statistically significant in the high-PDE state (**Figure 6F**) but highly significant in the low-PDE state (**Figure 6G**). Thus, the low GIV/low PDE signature carried a better prognosis compared to all other patients. Consistent with the fact that cAMP is a potent anti-tumor second messenger, these findings reveal that - 1) high levels of PDE signaling may not be a bad thing, especially when GIV levels are low; 2) in tumors with low PDE signaling, the low GIV signaling state may serve as a key synergy for driving up cAMP levels and therefore, critical for dampening several aggressive tumor traits (**Figure 1A**).

In the context of PDE, it has been demonstrated that overexpression of PDE isoforms in various cancers leads to impaired cAMP and/or cGMP generation [61]. PDE inhibitors in tumour models *in vitro* and *in vivo* have been shown to induce apoptosis and cell cycle arrest in a broad spectrum of tumour cells [62]. Despite the vast amount of preclinical evidence, there have been no PDE inhibitors that have successfully translated to the cancer clinics. For example, based on the role of cAMP in apoptosis and drug resistance, our model predicts that those with low GIV/high EGFR [high cAMP state] are likely to respond well to anti-EGFR therapy inducing tumor cell apoptosis, whereas those with high GIV/high EGFR [low cAMP state] may be at highest risk for developing drug resistance. Similarly, our finding that low PDE levels in the setting of high GIV carries a poor prognosis predicts that the benefits of PDE inhibitors may be limited to patients who have low GIV expression in their tumors. Whether such predictions hold true, remains to be investigated.

## Conclusions

Systems biology aims to understand and control the properties of biological networks; experimental data collected using top-down approaches are used to construct *in silico* bottom-up models, with the ultimate goal of generating experimentally testable predictions. In this work, we used a systems biology approach to construct the first-ever compartmental network model of growth-factor triggered cAMP signaling, and identified two key features of non-canonical G protein signaling via GIV-GEM. First, we identified that compartmentalized RTK signaling at the PM and on the endosomes directly imparts a *delayed* and *prolonged* cAMP dynamics lasting over an hour, which is distinct from the canonical GPCR/G protein pathway; GPCRs initiate more rapid and finite cAMP dynamics in the order of msec to min (**Figure 7A**) [63]. In the case of GPCRs, the PM-based signals are believed to be the dominant component of the overall cAMP dynamics with signal attenuation during endocytosis (**Figure 7A**) [37]. By contrast, in the case of RTK-mediated cAMP dynamics via GIV-GEM, the post-endocytic (i.e., endosomal signaling) component constitutes a dominant component of the overall cAMP dynamics that are triggered by RTKs (**Figure 7A**). What may be the impact of these distinct temporal features on RTK signaling? It is noteworthy that RTK-triggered cAMP dynamics that are modulated by GIV-GEM spans 5 min to > 60 min, which coincides with other RTK signaling, trafficking events and transcriptional response, i.e., the major temporal domain of RTK activity, the so-called “window of activity” [64]. We acknowledge that more work is needed to obtain better temporal and compartmental resolution of endosomal complexes including the G*α_s_·*GIV-GDI complexes. The 5 min to 1 h time scale encompasses the time of peak mRNA expression of many immediate-early genes (which peak at 20 min) and delayed-early genes (which peak between 40 min and 2 hours); these transcriptional targets not only generate feedback within the RTK-signaling cascade, but also set up crosstalk with other signaling pathways [64, 65]. In fact, GIV-GEM has indeed been found to modulate myriad downstream signaling pathways from the activity of small GTPases, kinases and phosphatases, to transcription factors [reviewed in [18]]; how GIV-GEM has such a widespread and broad impact had remained a mystery. It is possible that such broad impact could stem from GIV’s ability to modulate the cellular levels of the versatile second messenger cAMP in a sustained manner throughout the window of RTK activity although other mechanisms might also be at play. Although our model and simulations were carried out using the prototype RTK, EGFR, fundamentals identified here are likely to be relevant also in the case of others RTKs, *e.g*., VEGFR, IGF-1R, InsulinR, PDGFR, etc. Each of these RTKs are known to engage GIV and rely on its G protein modulatory function [reviewed in [18]] and can signal both at the PM and within endosomes [66-70]. Future work will explore the impact of such multi-receptor integration by GIV-GEM.

Second, our network model has helped us identify key design principles of the action of GIV-GEM within the EGF/EGFR signaling circuit by enabling construction of a map to identify the relationship between the key components– input[EGFR]→valve[GIV] →output[cAMP]→sink[PDE] (**Figure 7B-C**), validate the impact of such relationship with experimental assessment of cellular cAMP (**Figure 7D-E**) and interpret the role of each component in the context of network architecture ((**Figure 7F**). That there is a complex, non-linear and non-intuitive crosstalk between EGFR, GIV, and PDE in regulating cAMP levels is evident from the fact that the isoplanes, which capture the same cAMP AUC are not flat but are bent surfaces in this space. It appears that variation of cellular concentration of functionally active GIV-GEM molecules serves as the most tunable component that regulates the flow of signal from EGF/EGFR [input] to cAMP [output] (**Figure 7B-E**). At low concentrations of GIV, such as those found in normal tissues [**Figure S14**, **S15**], cAMP levels are sensitive to increased signal input via EGF/EGFR, i.e., higher input elicits higher output. Such sensitivity is virtually abolished and replaced by robustness at higher GIV concentrations found in a variety of cancers [**Figure 7 B**, **C**; **S14**, **S15**], i.e., higher input fails to elicit higher output and instead, cAMP levels stay at low and relatively constant. Regulation of cellular cAMP concentration by the EGFR-GIV interplay appears to be dependent on PDE concentration (**Figure 7D**); when PDE is high, there is virtually little or no effect of changes in EGFR or GIV on cAMP concentrations. The PDE-GIV relationship for cAMP production responds to different EGFR inputs proportionally, with increasing EGFR copy numbers resulting in higher cAMP AUC (**Figure 7E**). This 3-way interplay between EGFR, GIV and PDE is obvious also in experimental data derived from HeLa cells (**Figure 4, I, M**). Heat maps derived from that data (**Figure 7D-E**) show that the EGFR→cAMP pathway is most sensitive [i.e., higher input (EGF) signal, produces higher output (cAMP)] when the activities of both GIV and PDE are at their lowest (shGIV, with PDE inhibition; **Figure 7E**). Conversely, the EGFR→cAMP pathway is most robust [i.e., output (cAMP) is maintained at low levels despite higher input (EGF) signal] when both GIV and PDE are high (shC, without PDE inhibition; **Figure 7D**). When GIV is low and PDE is high (shGIV, without PDE inhibition; **Figure 7E**), cAMP levels do not go up, likely because increased production is balanced by increased degradation. Why would a cell waste energy (ATP) in such a ‘futile cycle’? This situation is reminiscent of the maintenance of steady-state cGMP levels in the sub-*μM* range in thalamic neurons by concomitant guanylyl cyclase and PDE2 activities [71] and cAMP levels in pyramidal cortical neurons by concomitant AC and PDE4 activities [72]. Prior studies have suggested that such tonic cAMP production and PKA activity enable signal integration and crosstalk with other cascades [73]; unlike an on/off system gated exclusively by G*α_s_* proteins, tonic activity allows both up- and downregulation by activation of G*α_i_* or inhibition of G*α_s_* (via GIV-GEM) and by PDEs. Our findings suggest that such up/down tunability is best achieved by changing the cellular concentrations of GIV. Because the flow of information in layers within signal transduction circuits in general [74-76], and more specifically for RTKs like EGFR [64, 77] is believed to conform to bow-tie microarchitecture, and cAMP is considered as one of the universal carrier molecules at the knot of such bow-ties which determines robustness [76], we conclude that GIV-GEM operates at the knot of the bow-tie as a tunable valve for controlling robustness within the circuit (see legend, **Figure 7F**). Because layering of control of [information] flow is believed to conform to an hourglass architecture [75], in which diverse functions and diverse components are intertwined via universal carriers, GIV’s ability to control the universal carrier, cAMP could explain why GIV has been found to be important for diverse cellular functions and impact diverse components [18]. In an hourglass architecture, the lower and higher layers tend to see frequent evolutionary changes, while the carriers at the waist of the hourglass appear to be constant/invariant and sometimes, virtually ‘ossified’. Of relevance to our model, the importance of cAMP appears to be indeed ossified from unicellular organism to human alike, and GIV-GEM is expressed in ubiquitously in all tissues from fish to man and GIV-like GEMs have so far been identified as early as in C. *elegans* [78].

Third, our work also provides valuable clues into the impact of increased robustness at high-GIV states in cancers. Robustness in signaling is an organizing principle in biology, not only for the maintenance of homeostasis but also in the development and progression of chronic debilitating diseases like cancers; it is widely accepted that tumor cells hijack such robustness to gain growth and survival advantage during the development of cancer [64, 79, 80]. Consistently, we found that GIV mRNA levels and DNA copy numbers are invariably higher across multiple cancers when compared to their respective normal tissue of origin (**Figure S14**, **S15**). Because GIV has been found to regulate several harmful properties of tumor cells across a variety of cancers (multiple studies, reviewed in [20]), it is possible that the high-GIV driven robustness maintains cAMP at low constant levels despite increasing input signals as a tumor evolves when targeted by biologicals or chemotherapy agents. Such a phenomenon could be a part of a higher order organizing principle in most aggressive cancers, and therefore, justify GIV as a potential target for network-based anti-cancer therapy.

Although our model captures experimentally observed time courses and generates testable hypotheses, it has 3 major limitations. First, the compartmental well-mixed model we used, does not account for the spatial location and geometries of the different compartments and cell shape, many of which can affect the dynamics of cell signaling [81, 82]. Second, our model focuses exclusively on cAMP as output signal and does not account for other EGF/EGFR-driven signaling pathways that are known to regulate cellular responses as these likely would require their own study, such as the Ras-Raf-MEK-ERK pathway which is known to modulate cAMP and be modulated by GIV through affecting adaptor protein recruitment [24]. Third, our model focuses exclusively on EGFR and does not account for the diverse classes of receptors [multiple RTKs, GPCRs, integrins, etc.] that also use GIV to access and modulate G proteins. Despite these restrictions, we can identify some fundamental features of growth factor-triggered cAMP signaling for the first time using systems biology, including the role of compartmentalization, cross-talk between EGFR and GIV, GIV-dependent robustness within the RTK-cAMP signaling axis, and cross-talk between PDE and GIV in controlling cAMP concentration.

We conclude that GIV utilizes compartmental segregation to modulate the dynamics of RTK-G protein-cAMP signaling and confers robustness to these dynamics by functioning as a tunable control valve. Future systems efforts will build on this model to unravel further exciting features of GIV as a critical hub for signaling regulation at the knot of a bowtie [76] and elucidate the hidden complexity that arises from network architecture in non-canonical G protein signaling.

## Methods

### Modular construction of the reaction network

A biochemical network model was constructed to capture the main events in the signal transduction cascade from EGF to cAMP through GIV (**Figure 1D**). We constructed the compartmental computational model in a modular manner, where each module represents key events within the network. The model was trained using key data sets published over the past decade on GIV-GEM, most notably, those that defined the spatiotemporal kinetics of EGFR·GIV, EGFR·GIV·G*α_i_* interactions [24, 26, 31], dynamics of phosphoregulation of GIV-GEM [30, 33], and most importantly, the dual modulation of G*α_i_*_/_*_s_* by GIV-GEM that is brought about by temporally and spatially separated phosphorylation events [22]. Last, but not least, we also used published role of G*α_s_* in the feedback regulation of endocytic downregulation of EGF/EGFR signaling [28]. We note here that while there are many more biochemical components involved in signaling from EGF to cAMP, our choice of components was based on experimentally measured temporal dynamics of GIV-GEF and GIV-GDI functions. The modules are as follows.

**Module 1** consists of EGFR activation through EGF and internalization dynamics leading to the formation of EGFR·GIV·G*α_i_* trimeric complex. This module includes the phenomenon that endosomal maturation and EGFR degradation in lysosomes requires the presence of inactive G*α_s_* [GDP-bound state] [28]. The presence of G*α_s_* in the inactive state promotes maturation of endosomes, shuts down the mitogenic MAPK-ERK1/2 signals from endosomes and suppresses cell proliferation [28]. In the absence of G*α_s_* or in cells expressing a constitutively active mutant G*α_s_*, EGFR stays longer in endosomes, MAPK - ERK1/2 signals are enhanced and cells proliferate [28] (**Figure 2A**).

**Module 2** contains EGFR-mediated activation of PLC-*γ* and downstream activation of PKC-*θ*; the latter phosphorylates GIV at S1689 and terminates its ability to activate G*α_i_* [33]. This phosphoevent does not impact GIV’s ability to inhibit G*α_s_* [22]. Consequently, when it comes to G protein modulatory functions of GIV, phosphorylation by PKC-*θ* converts GIV-GEF into GIV-GDI (**Figure 3F**).

**Module 3** contains the dynamics of the endosomal EGFR and how it activates G*α_s_*, and subsequently adenylyl cyclase (AC) leading to the synthesis of cAMP (**Figure 3A**).

**Module 4** contains the dynamics of the AC and how it synthesizes cAMP, leading to downstream effectors and controllers (**Figure 3A**) we currently only consider PDE feedback for cAMP reduction. Overall, the model contains 54 reactions. The complete set of reactions for each of the modules, their parameters and interactions, and the list of assumptions underlying network construction are provided as online supplementary materials (Tables **S2** - **S6**).

We assumed that the signaling components were present in large-enough quantities, and different concentrations of each component were computed to explore how varying expression levels in different tissues/cell types impact the signaling pathway. Such assumption allowed us to generate a deterministic dynamical model. The model contains six different compartments: (i)PM, (ii) extracellular space, (iii) cytosol, (iv) endosomes, and (v) endosomal membranes. It was assumed that each compartment is well-mixed and fluxes were used to depict transport across the different compartments so that the dynamic changes in the concentrations of the different components can be tracked. Each interaction was modeled as a chemical reaction either using mass-action kinetics for binding-unbinding reactions, and Michaelis-Menten kinetics for enzyme-catalyzed reactions, as is standard for models such as this [83, 84].

The network of interactions was constructed using the Virtual Cell modeling platform (http://www.nrcam.uchc.edu). We chose this platform because it is a user-friendly computational cell biology software, which allows us to generate the system of differential equations based on the input reactions and has been used successfully to model signaling networks of various sizes with a high degree of numerical accuracy [85-88]. Also, the Virtual Cell platform has built-in capabilities to conduct dynamic sensitivity analysis, which is an important aspect of dynamic systems modeling. As we discuss in later sections, we use this capability to identify sources of system robustness and sloppiness.

### Characteristics of the signaling cascade

In order to characterize the dynamics of the different protein activities, we use the area under the curve for the concentration versus time curve [48]. The area under the curve gives the total signal activated over the time of observation and for the *i^th^* species is given by *AUC_i_* in Eq. **1**. This gives a measure of the total signal for different conditions.

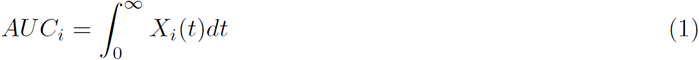

### Comparison with experimental data

The experimental data was extracted from Figure 1D of [22] using ImageJ for GEF-G*α_i_*-EGFR complex and G*α_s_*-GDI complex. The data was normalized such that the maximum value was 1. Parameter fitting using COPASI [89] was used to then match the normalized experimental data against the model output. Goodness of fit between experimental values and model output was determined using a root mean squared error (RMSE).

### Dynamic parametric sensitivity analysis

Since a continuing challenge in building computational models of signaling networks is the choice of kinetic parameters, we conducted a dynamic parametric sensitivity analysis. This sensitivity analysis of the model was performed with the goal of identifying the set of parameters and initial concentrations that the model response is most sensitive to. The log sensitivity coefficient of the concentration of the *i^th^* species *C_i_*, with respect to parameter *k_j_* is given by [90, 91]

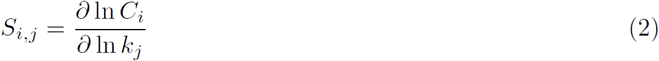

Since we are studying a dynamical system and not steady state behavior, we used the Virtual cell software to calculate the change in log sensitivity over time 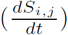. The resulting time course gives us information about the time dependence of parametric sensitivity coefficients for the system. The variable of interest, *C_i_* is said to be robust with respect to a parameter *k_j_* if the log sensitivity is of the order 1 [90]. We refer the reader to [90, 91] for a complete introduction to dynamical sensitivity analysis. We conducted dynamic sensitivity analysis for all the kinetic parameters for the reactions and initial concentrations of the different species in the model.

## Measurement of cAMP

HeLa cells were serum starved (0.2 % FBS, 16 h) and incubated with isobutylmethylxanthine (IBMX, 200 *μM*, 20 min) followed by EGF. Stimulation was carried out either using fixed EGF concentrations followed by assessment of cAMP at various time points (as in **Figure 4C**) or using varying EGF concentrations followed by an assessment of cAMP at 60 min (as in **Figure 4I,M**). Reactions were terminated by aspiration of media and addition of 150 *μl* of ice-cold TCA 7.5% (w/v). cAMP content in TCA extracts was determined by radioimmunoassay (RIA) and normalized to protein [(determined using a dye binding protein assay (Bio-Rad)] [18, 92]. Data is expressed as fmol cAMP / *μg* total protein.

### Stratification of colon cancer patients in distinct gene-expression subgroups and comparative analysis of their survival outcomes

The association between the levels of GIV (CCDC88A) and either EGFR or PDE mRNA expression and patient survival was tested in cohort of 466 patients where each tumor had been annotated with the disease-free survival (DFS) information of the corresponding patient. This cohort included gene expression data from four publicly available NCBI-GEO data-series (GSE14333, GSE17538, GSE31595, GSE37892) [93-96], and contained information on 466 unique primary colon carcinoma samples, collected from patients at various clinical stages (AJCC Stage I-IV/Duke’s Stage A-D) by five independent institutions: 1) the H. Lee Moffit Cancer Center in Tampa, Florida, USA (n = 164); 2) the Vanderbilt Medical Center in Nashville, Tennessee, USA (n = 55); 3) the Royal Melbourne Hospital in Melbourne, Australia (n = 80); 4) the Institut PaoliCalmette in Marseille, France (n = 130); 5) the Roskilde Hospital in Copenhagen, Denmark (n = 37). To avoid redundancies (i.e. identical samples replicated two or more times across multiple NCBI-GEO datasets) all 466 samples contained in this subset were cross-checked to exclude the presence of duplicates. A complete list of all GSMIDs of the experiments contained within the NCBI-GEO discovery dataset has been published previously [50]. To investigate the relationship between the mRNA expression levels of selected genes (i.e. CCDCDDC, Wnt5a, EGFR and FZD7) and the clinical outcomes of the 466 colon cancer patients represented within the NCBI-GEO discovery dataset, we applied the Hegemon software tool [50]. The Hegemon software is an upgrade of the BooleanNet software [97], where individual gene-expression arrays, after having been plotted on a two-axis chart based on the expression levels of any two given genes, can be stratified using the StepMiner algorithm and automatically compared for survival outcomes using Kaplan-Meier curves and logrank tests. Since all 466 samples contained in the dataset had been analyzed using the Affymetrix HG-U133 Plus 2.0 platform (GPL570), the threshold gene-expression levels for GIV/CCDC88A, PDE and EGFR were calculated using the StepMiner algorithm based on the expression distribution of the 25, 955 experiments performed on the Affymetrix HG-U133 Plus 2.0 platform. We stratified the patient population of the NCBI-GEO discovery dataset in different gene-expression subgroups, based on either the mRNA expression levels of GIV/CCDC88A alone (i.e. CCDC88A neg vs. pos), PDE alone (i.e., PDE neg vs. pos), EGFR alone (i.e. EGFR neg vs. pos), or a combination of GIV and either EGFR or PDE. Once grouped based on their gene-expression levels, patient subsets were compared for survival outcomes using both Kaplan-Meier survival curves and multivariate analysis based on the Cox proportional hazards method.

## Acknowledgments

This work was supported by ARO W911NF-16-1-0411, AFOSR FA9550-15-1-0124, and NSF PHY-1505017 grants to P.R. and NIH grants CA100768, CA160911 and DK099226 (to P.G). M.G. was supported by the UCSD Frontiers of Innovation Scholars Program (FISP) G3020, and by a grant from the National Institutes of Health, USA (NIH grant T32EB009380). L.S. was supported by T32DK0070202, Chancellor’s Research Excellence Scholarships (CRES) for Graduate Students (UCSD), and a graduate research fellowship from the microbial sciences initiative (MSI, also at UCSD). The *Virtual Cell* suite (http://vcell.org) is supported by National Institute for General Medical Sciences, NIH (Grant Number P41 GM103313).

## Supplementary Material

### Supplementary Movie Legends

**Supplementary Movie 1:** Activation of G*α_s_* in response to EGF, as determined by nanobody Nb37-GFP in control HeLa cells [shControl]. The movie shows EGF-dependent activation of G*α_s_* as detected by live-cell imaging using the G*α_s_* conformational biosensor nanobody Nb37-GFP that binds and helps detect the nucleotide-free intermediate during G*α_s_* activation [1]. In control HeLa cells responding to EGF little or no G*α_s_* activity was seen. Quantification of these findings have been published in [2] (Magnification, 63 x).

**Supplementary Movie 2:** Activation of G*α_s_* in response to EGF, as determined by nanobody NB37-GFP in GIV-depleted HeLa cells (shGIV). The movie show EGF-dependent activation of G*α_s_* as detected by live-cell imaging using the G*α_s_* conformational biosensor nanobody Nb37-GFP that binds and helps detect the nucleotide-free intermediate during G*α_s_* activation [1]. Compared with controls (Movie S1), in GIV-depleted cells a significant increase in G*α_s_* activity was seen on vesicular structures that are likely to be endosomes. Quantification of these findings have been published in [2] (Magnification, 63x).

### 2 Introduction to modeling chemical reactions

#### 2.1 Mass-action kinetics

We generated an ordinary differential equation (ODE) for every species using mass-action kinetics for each binding reaction. The law of mass action states that the rate of a chemical reaction is proportional to the product of the concentration of the reactants raised to the power of their stoichiometric coefficient. For example, consider the one-reaction system:

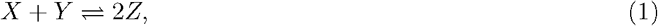
where the forward and backward rates are *k*_1_ and *k*_2_. The differential equations describing the dynamics of species *X, Y*, and *Z* under mass-action kinetics are:

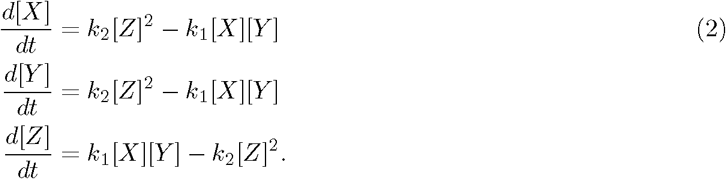

Mass action kinetics rely on the assumption that the rate constant, *k*, is constant over time. However, within a restricted space such as a membrane, the rate constant may change over time due to restricted diffusion and mass action kinetics may not be accurate [3]. We assumed that most binding interactions occur rapidly enough such that k remains constant.

#### 2.2 Michaelis-Menten kinetics

We used Michaelis-Menten kinetics to model enzyme-catalyzed reactions. When a reaction is catalyzed by an enzyme with kinetic properties *k_cat_* and *K_M_*,

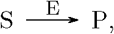
then the reaction rate is given by

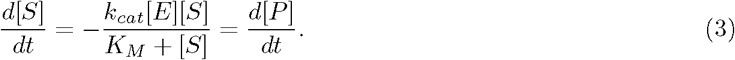

For Michaelis-Menten kinetics to apply, the concentrations of the reactants and products must be in large enough quantities, and one of the following conditions must apply:

1. The concentration of substrate is very much larger than the concentration of products: [S]≫[P].
2. The energy released in the reaction is very large: ∆*G* ≪ 0.

#### 2.3 Transport between compartments

Flux between different cellular compartments was modeled as a reaction rate that captures the rate of species transport per unit time. For the transport of species A between compartments c and d, we used rate equations of the form

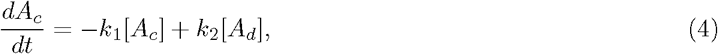
where *k*_1_ and *k*_2_ are the transport in and out of compartment c respectively. When utilizing flux for compartmental transport it is important to note the interaction is only valid when the compartments are large and the corresponding surface area conversion factors are accounted for. Reactions that used fluxes to model different cellular components as follows: phosphorylated EGF-EGFR_2_ internalization [R4], EGFR internalization [R8], G*α_i_* transport to an endosome [R19], G*α_i_*-GDP transport to PM [R21], AC internalization [R32], and G*α_s_* internalization [R37].

### 3 Model development for growth-factor based cAMP signaling

#### 3.1 Compartment sizes

We conducted simulations using the following compartments for a computational HeLa cell: cytoplasm, plasma membrane, endosome, endosomal membrane, and a nucleoplasm. We assumed that the cell was spherical shaped and used a cytosolic volume of 2000 *μm*^3^ [4]. We assumed the endosomes to be fixed in size during the time course of signaling, with a diameter of 87 nm [5], slightly smaller than the size of a large endosome (100nm). Villasenor *et al*. [6] reported that about 50 endosomes are created after 30 min of 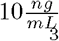 EGF stimulation in HeLa cells. Using this value, we calculated the total endosomal volume to be 0.138 *μm*^3^ and surface area of 5 *μm*^2^ using 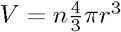 and membrane area by A = *n*4*πr*^2^, where n is the final number of endosomes. The different compartment sizes are shown in **Table S1**.

**Table S1:** Sizes of different compartments used within the model

| Compartment | size | Notes and References |
| --- | --- | --- |
| EC | $5000 \mu m^3$ | - |
| PM | $1256 \mu m^2$ | [4] |
| Cytosol | $2000 \mu m^3$ | [4] |
| Endosome | $0.138 \mu m^3$ | [6], Est. |
| Endosomal membrane | $5 \mu m^2$ | [6], Est. |
| Nucleosol | $200 \mu m^3$ | Assumed 10% of Cytosol |

#### 3.2 Model Kinetics

We conducted simulations for 60 min based on the time course of RTK→cAMP signaling [2]. We did not account for the regeneration of ATP and PIP2, and assumed that these values are constant and high. We did not include mitogen-activated protein kinase (MAPK) or calcium pathways in this model.

#### 3.3 Module 1: EGF Receptor Module

The receptor module captures the key events of

1. Ligand binding and dimerization.
2. Receptor activation and internalization.
3. Receptor endosomal recycling.
4. Receptor G*α_s_*-GDP dependent degradation.

This module is based on Shoerberl *et al*. [5] for EGFR activation, internalization, and recycling. Binding of EGFR to scaffolding proteins was not included in this model. G*α_s_*·GIV-GDI dependent degradation of EGFR was modeled based on [7]. Although the exact mechanism of G*α_s_*-GDP based degradation is unknown, we used a constitutive model to capture the effect of degradation of EGFR by G*α_s_* using a G*α_s_*-GDP independent basal rate and a G*α_s_*-GDP dependent catalytic rate.

##### 3.3.1 Kinetic parameters

The kinetic binding parameters were chosen based on the values given in Schoeberl *et al*. [5], Berkers *et al*. [8], and French *et al*. [9]. The rate of EGF binding to EGFR at the plasma membrane was set through values reported in Berkers *et al*. [8]. The binding rate of endosomal EGF to EGFR was set by the relative ratio of pH=6 and pH=7.4 as shown in French *et al*. [9]. Degradation of EGFR was set through the global parameters *k_b_*, basal degradation, and *k_c_*, G*α_s_*-GDP-dependent degradation. The internalization and degradation rates were modified to fit experimental data for receptor internalization as shown in [6, 10] and for experimental data shown in **Figure 2D**, **3G**, **S4**,**S5**.

#### 3.4 Module 2: Transactivation of Ga_i_ by EGFR via GIV-GEF interactions

The GIV-GEF module captures the key events of

1. Activation of GIV through CDK5.
2. GIV·EGFR binding and amplification of receptor signaling events.
3. Formation of EGFR·GIV·G*α_i_* complex and activation of G*α_i_*.

The activating step for GIV-GEF is CDK5-mediated phosphorylation of S1674 on GIV [2]. GIV-GEF is then later turned “off” by PKC-*θ*, which is activated downstream of PLC-*γ*. Once activated, GIV-GEF binds to EGFR and G*α_i_*-GDP to assemble the EGFR·GIV·G*α_i_* complex [11]. Previously published pathways, kinetics and dynamics of CDK5 activation were used to build the model [12-14] (see ‘Kinetic parameters’ section below). We did not track the dynamics of *βγs* in this model.

##### 3.4.1 Kinetic parameters

Kinetic parameters were determined through a combination values from the literature and experimental data fitting. Activation of p35 by the receptor (**Table S3** reaction 12), was determined using rates from Bhalla *et al*. [15] and by fitting simulations to new (cAMP timecourse) and previously published experimental data (GIV-GEM IB), **Figure 2D**, **3G**, **S4**, **S5** [2]. The rate of p35 degradation was determined based on the known half-life of 20 to 30 min [16] then refit to the GIV-GEF curve, **Figure S4**. Maximum binding of CDK5 to p35 (**Table S3** reaction 13), was set to 80% based on published experimental data [12]. Binding of CDK5 to active p35 was assumed to be very rapid. The rate of GIV-GEF activation by CDK5 (**Table S3** reaction 14), was fit to immunoblotting data [2] (**Figure 2D**, **3G**, **S4**, **S5**). EGFR_2_·GIV and EGFR·GIV·G*α_i_* formation rates were determined by fitting of experimental data using COPASI [17] and using the experimentally determined dissociation constant (*K_d_*) of EGFR·GIV·G*α_i_* formation [11, 18].

#### 3.5 Module 3: Transinhibition of G*α_s_* by EGFR via GIV-GDI

The GIV-GDI module captures the key events of

1. PLC-*γ* activation and PIP_2_ hydrolysis.
2. Enhanced PLC-*γ* activation through EGFR_2_·GIV.
3. DAG dependent PKC-*θ* activity.
4. Termination of GIV-GEF [for G*α_i_*], and its conversion to GIV-GDI through PKC-*θ* and reduction of GEF activity.

The action of PKC-*θ* on GIV was based on prior work [2], which showed that targeted phosphorylation on site S1689 terminates GIV’s GEF function, only allowing GDI function to be active. For the purposes of our model, it was assumed that the PLC-*γ*→PKC-*θ* axis acts after CDK5, as shown previously [2]. In doing so, the PLC-*γ*→PKC-*θ* axis phosphorylates GIV-GEF that is activated by CDK5, but not inactive GIV [2].

In the model, activation of PKC-*θ* was achieved through the action of PLC-*γ*. PLC-*γ* activation was modeled to be a function of both EGFR and EGFR_2_·GIV; the latter assumption was made based on prior work [11], which showed that GIV enhances EGF triggered PLC-*γ* signaling. Once active, PLC-*γ* hydrolyzes PIP_2_, creating IP_3_ and DAG [19]; DAG then binds and activates PKC-*θ*, inducing the localization of the latter to the PM. PKC-*θ* then phosphorylates GIV-GEF in both the unbound and the receptor bound form. G*α_s_*·GIV-GDI complex formation and function was based on [2, 7] where it was shown to only act of the GDP form of G*α_s_*.

##### 3.5.1 Kinetic parameters

The initial choice of kinetic parameters were based on the previously published rates for activation of PLC-*γ*, PIP_2_, and IP_3_ degradation [15]. These parameters were then refined by fitting the dynamics of the G*α_s_*·GIV-GDI complex to immunobloting data [2] (**Figure 2D**, **3G**, **S4**, **S5**); the rate of GDI activation through PKC-*θ* and the rate of formation of the G*α_s_*·GIV-GDI complex were determined by fitting simulations to immunoblot data (**Figure 2D**, **3G**, **S4**, **S5**).

#### 3.6 Module 4: Reactions for the production and degradation of cAMP

The cAMP module captures the key events of

1. Inhibition of basal activity of the PM-pool of AC by G*α_i_*.
2. Activation of endosome-pool of AC by G*α_s_*.
3. Internalization of AC, G*α_s_*, G*α_i_*.
4. Production of cAMP, activation of PKA and PDE.

In the model, we assumed that AC activation through EGFR occurs only on the endosome [13, 20], because we assumed that EGFR is only able to activate G*_s_* proteins on the endosome. Binding of internalized G*α_s_*-GTP to AC activates and allows increased catalytic activity of AC [21]. The binding of G*α_i_* to AC was modeled to reflect the inhibition of all AC activity [22]. AC inhibition was allowed to occur on both membranes.

cAMP production by AC was modeled with Michaelis-Menten kinetics [21]. Because cellular ATP is in the millimolar range [23], a large excess compared to the concentrations of the signaling molecules, the concentration of ATP was assumed to be constant. Once four cAMP molecules bind to the four distinct binding sites on PKA, the quadruple occupancy leads to activation of the catalytic subunit, PKAc, which separates from regulatory subunit [24]. In our model, PKA activation was modeled using a Hill equation [25]. We assumed that PKAc had the same steady state concentration in the nucleosol as the cytosol. PDE activation through PKAc and enzymatic function was based on [21]; cAMP is degraded by PDE. We did not consider any AKAPs or AKIPs because they fall outside the scope of the current model; it possible that their inclusion may impact response strength and timescales. removed mentions of CREB.

##### 3.6.1 Kinetic parameters

Previously published activation kinetics of the AC→cAMP→PKA cascade [21] were modified to be closer to observed experimental values [26, 27]. Alousi *et al*. [26] estimated a ratio of G*α_s_* to AC of greater than 33.3, while Post [27] reported a value of 78.3 in isolated adult rat ventricular myocytes. We therefore chose a ratio of 50 to set our inital values. The binding rates of G*α_s_*-GTP and G*α_i_*-GTP to AC were based on values in [21]. Rates of inactivation of G*α_s_* and G*α_i_* bound to AC were determined by the GTP hydrolysis activity of AC [28]. Internalization rates of G*α_s_* and AC were defined as the same parameter, *k_int_*, determined through fitting to immunoblot data [2] (**Figure 2D**, **3G**, **S4**, **S5**). The kinetic rates governing PKA activity were determined by using steady state dose-response curves to fit a Hill equation (**Figure S7**) [25], dissociation of cAMP from the regulatory subunits [29], and reformation of the PKA holoenzyme.

#### 3.7 Role of additional interactions from CDK5 and PKC-*θ* to PDE

Previous studies have shown that both CDK5 [30] and PKC-*θ* [31] influence PDE activation. We modeled interactions of CDK5 and PKC-*θ* to activation of PDE, leading to further suppression of the cAMP signal; the additional interactions are shown in **Table S7**. The addition of these interactions did not alter the time course of cAMP production but reduced the amount of cAMP produced **Figure S10**.

##### 3.7.1 Kinetic parameters

We tested the role of additional PDE activation pathways through CDK5 and PKC-*θ* [30, 31]; We assumed these interactions would double the maximum PDE concentration based on previously published data [30, 31] that has shown that suppressing the affects of CDK5 and PKC-*θ* on PDE reduced PDE activity by 1.25 to 1.5-fold.

### 4 Model access in Virtual Cell program [VCell]

The simulations for the full network, shown in **Figure 1**, were carried in the Virtual Cell program (http://vcell.org/). The model is named ‘GIV-GEM Paradoxical Signaling’, with additional models ‘GIV-GEM Paradoxical Signaling GsPM’ (PM activities of Gs Protein) and ‘GIV-GEM Paradoxical Signaling CDKPKC’ (PKC and CDK PDE interactions), and is available under the publicly shared models with the username ‘mgetz’. The Virtual Cell is supported by NIH Grant Number P41 GM103313 from the National Institute for General Medical Sciences. Complete instructions on how to access publicly shared models can be found on the Virtual Cell homepage. A detailed protocol/user guide on how to develop models in Virtual Cell has been published elsewhere [32, 33].

#### 4.1 Parameter estimation with COPASI

Parameter estimation was carried out using COPASI [17] built into the VCell program [32]. Parameter estimation was conducted simultaneously on GIV-GEF, GIV-GDI, cAMP, EGFR·GIV·G*α_i_*, and G*α_s_*·GIV-GDI blot data from experiments. The particular estimation method used was evolutionary programming (EP) for 300 generations with a 25 population size using 1 random number generator. This was performed for two runs before results were reported. Evolutionary programming functions as follows: once presented with a optimization problem, i.e. minimizing error between the absolute values of experimental data and simulation data (E=|*y*_1_ − *y*_2_|), the algorithm develops a set of potential solutions by varying parameters chosen by the user. At the next “generation” each individual solution produces two “offspring” one of the same solution and one with slight random parameter variations. Therefore, at the end of this generation there are double the number of potential solutions. These are then reduced to the original number of solutions by comparing the error against the other solutions, keeping only the lowest errors and deleting all other solutions. For more information on this method see [34].

**Table S2:**
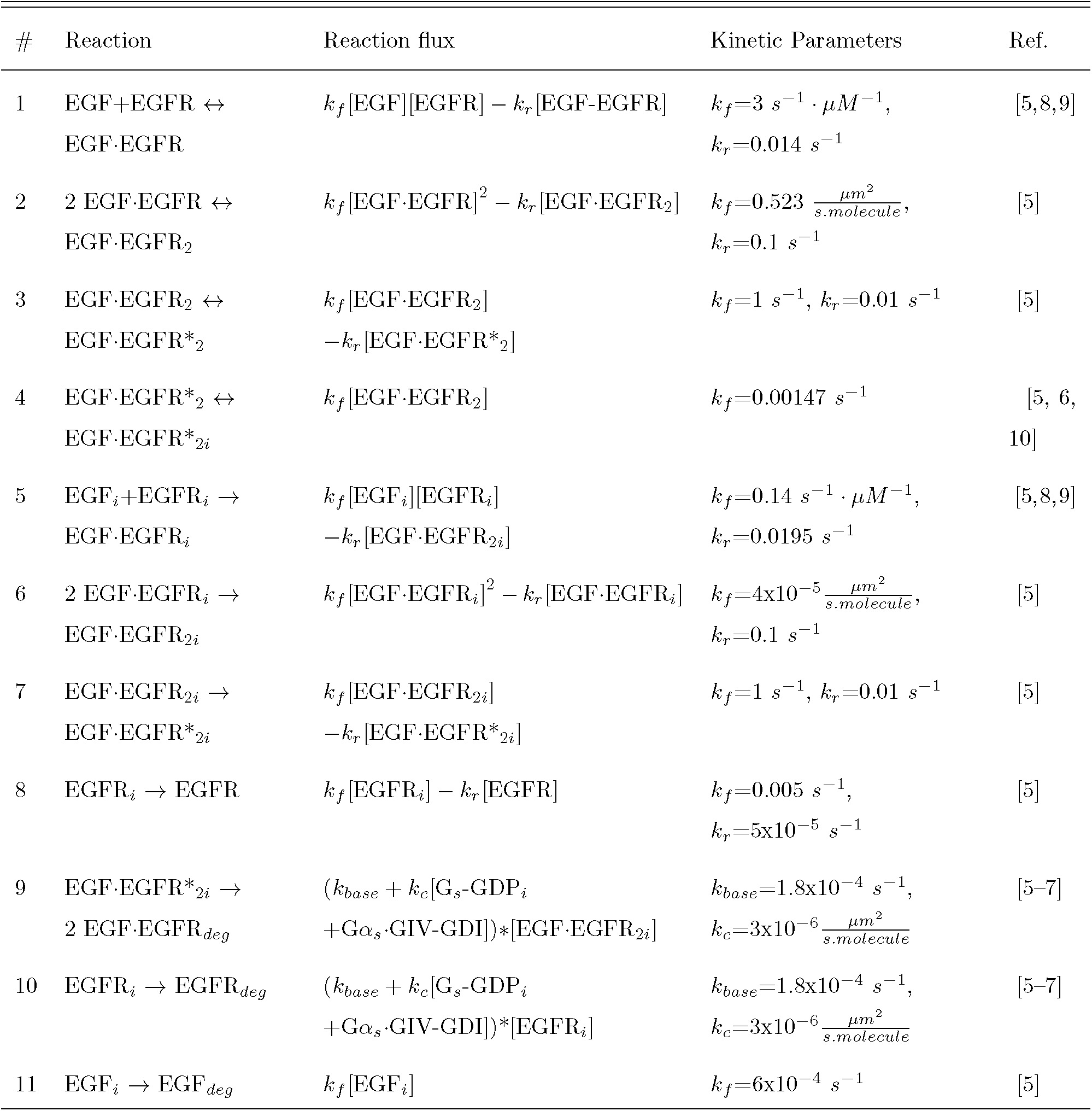
Reactions for **Module 1** (**Figure 2A**), outlining EGFR activation, internalization, and degradation.

| # | Reaction | Reaction flux | Kinetic Parameters | Ref. |
| --- | --- | --- | --- | --- |
| 1 | EGF+EGFR ↔<br>EGF·EGFR | $k_f[\text{EGF}][\text{EGFR}] - k_r[\text{EGF}\cdot\text{EGFR}]$ | $k_f=3 \text{ s}^{-1} \cdot \mu\text{M}^{-1},$<br>$k_r=0.014 \text{ s}^{-1}$ | [5,8,9] |
| 2 | 2 EGF·EGFR ↔<br>EGF·EGFR<sub>2</sub> | $k_f[\text{EGF}\cdot\text{EGFR}]^2 - k_r[\text{EGF}\cdot\text{EGFR}_2]$ | $k_f=0.523 \frac{\mu\text{m}^2}{\text{s}\cdot\text{molecule}},$<br>$k_r=0.1 \text{ s}^{-1}$ | [5] |
| 3 | EGF·EGFR<sub>2</sub> ↔<br>EGF·EGFR*<sub>2</sub> | $k_f[\text{EGF}\cdot\text{EGFR}_2]$<br>$-k_r[\text{EGF}\cdot\text{EGFR}^*_2]$ | $k_f=1 \text{ s}^{-1}, k_r=0.01 \text{ s}^{-1}$ | [5] |
| 4 | EGF·EGFR*<sub>2</sub> ↔<br>EGF·EGFR*<sub>2i</sub> | $k_f[\text{EGF}\cdot\text{EGFR}_2]$ | $k_f=0.00147 \text{ s}^{-1}$ | [5, 6,<br>10] |
| 5 | EGF<sub>i</sub>+EGFR<sub>i</sub> →<br>EGF·EGFR<sub>i</sub> | $k_f[\text{EGF}_i][\text{EGFR}_i]$<br>$-k_r[\text{EGF}\cdot\text{EGFR}_{2i}]$ | $k_f=0.14 \text{ s}^{-1} \cdot \mu\text{M}^{-1},$<br>$k_r=0.0195 \text{ s}^{-1}$ | [5,8,9] |
| 6 | 2 EGF·EGFR<sub>i</sub> →<br>EGF·EGFR<sub>2i</sub> | $k_f[\text{EGF}\cdot\text{EGFR}_i]^2 - k_r[\text{EGF}\cdot\text{EGFR}_i]$ | $k_f=4\text{x}10^{-5} \frac{\mu\text{m}^2}{\text{s}\cdot\text{molecule}},$<br>$k_r=0.1 \text{ s}^{-1}$ | [5] |
| 7 | EGF·EGFR<sub>2i</sub> →<br>EGF·EGFR*<sub>2i</sub> | $k_f[\text{EGF}\cdot\text{EGFR}_{2i}]$<br>$-k_r[\text{EGF}\cdot\text{EGFR}^*_{2i}]$ | $k_f=1 \text{ s}^{-1}, k_r=0.01 \text{ s}^{-1}$ | [5] |
| 8 | EGFR<sub>i</sub> → EGFR | $k_f[\text{EGFR}_i] - k_r[\text{EGFR}]$ | $k_f=0.005 \text{ s}^{-1},$<br>$k_r=5\text{x}10^{-5} \text{ s}^{-1}$ | [5] |
| 9 | EGF·EGFR*<sub>2i</sub> →<br>2 EGF·EGFR<sub>deg</sub> | $(k_{base} + k_c[\text{G}_s\text{-GDP}_i$<br>$+ \text{G}\alpha_s \cdot \text{GIV-GDI}]) * [\text{EGF}\cdot\text{EGFR}_{2i}]$ | $k_{base}=1.8\text{x}10^{-4} \text{ s}^{-1},$<br>$k_c=3\text{x}10^{-6} \frac{\mu\text{m}^2}{\text{s}\cdot\text{molecule}}$ | [5-7] |
| 10 | EGFR<sub>i</sub> → EGFR<sub>deg</sub> | $(k_{base} + k_c[\text{G}_s\text{-GDP}_i$<br>$+ \text{G}\alpha_s \cdot \text{GIV-GDI}]) * [\text{EGFR}_i]$ | $k_{base}=1.8\text{x}10^{-4} \text{ s}^{-1},$<br>$k_c=3\text{x}10^{-6} \frac{\mu\text{m}^2}{\text{s}\cdot\text{molecule}}$ | [5-7] |
| 11 | EGF<sub>i</sub> → EGF<sub>deg</sub> | $k_f[\text{EGF}_i]$ | $k_f=6\text{x}10^{-4} \text{ s}^{-1}$ | [5] |

**Table S3:** Reactions for **Module 2** (**Figure 2C**), outlining protein-protein interactions leading to the transactivation of G*α_i_* by EGFR via GIV-GEF.

| # | Reaction | Reaction flux | Kinetic Parameters | Ref. |
| --- | --- | --- | --- | --- |
| 12 | $p35 \rightarrow p35^*$ | $\frac{k_{cat}[EGF \cdot EGFR_2][p35]}{K_M + [p35]} - k_r[p35^*]$ | $K_M = 957 \frac{molecule}{\mu m^2}$,<br>$k_{cat} = 0.1 s^{-1}$, $k_r = 0.001 s^{-1}$ | [16, 35] |
| 13 | $p35^* + CDK5 \rightarrow CDK5^*$ | $k_f[p35^*][cdk5] - k_r[cdk5^*]$ | $k_f = 45 \mu M^{-1} \cdot s^{-1}$,<br>$k_r = 10 s^{-1}$ | [12, 13] |
| 14 | $GIV \rightarrow GIV-GEF$ | $\frac{k_{cat}[CDK5^*][GIV]}{K_M + [GIV]} - k_r[GIV-GEF]$ | $K_M = 6 \mu M$, $k_{cat} = 0.3 s^{-1}$,<br>$k_r = 0.0026 s^{-1}$ | Est.<sup>1</sup>,<br>[12] |
| 15 | $GIV-GEF + EGF \cdot EGFR^*_2 \leftrightarrow EGFR_2 \cdot GIV$ | $k_f[GIV-GEF][EGF \cdot EGFR^*_2] - k_r[EGFR_2 \cdot GIV]$ | $k_f = 0.021 \mu M^{-1} \cdot s^{-1}$<br>$k_r = 6.48 \times 10^{-4} s^{-1}$ | [11] |
| 16 | $G\alpha_i-GDP + EGFR_2 \cdot GIV \leftrightarrow EGFR_2 \cdot GIV \cdot G\alpha_i$ | $k_f[EGFR_2 \cdot GIV][G\alpha_i-GDP] - k_r[EGFR_2 \cdot GIV \cdot G\alpha_i]$ | $k_f = 0.005 \frac{\mu m^2}{s \cdot molecule}$,<br>$k_r = 1.142 s^{-1}$,<br>($K_d = 0.24 \mu M$) | [18] |
| 17 | $EGFR_2 \cdot GIV \cdot G\alpha_i \rightarrow G\alpha_i-GTP + EGFR_2 \cdot GIV$ | $k_f[EGFR_2 \cdot GIV \cdot G\alpha_i]$ | $k_f = 4.7 s^{-1}$ | [15] |
| 18 | $G\alpha_i-GTP \rightarrow G\alpha_i-GDP$ | $k_f[G\alpha_i-GTP]$ | $k_f = 0.139 s^{-1}$ | [21] |
| 19 | $G\alpha_i-GTP \rightarrow G\alpha_i-GTP_i$ | $k_f[G\alpha_i-GTP] - k_r[G\alpha_i-GTP_{int}]$ | $k_f = 0.05 s^{-1}$, $k_r = 0.001 s^{-1}$ | Est.<sup>2</sup> |
| 20 | $G\alpha_i-GTP_i \rightarrow G\alpha_i-GDP_i$ | $k_f[G\alpha_i-GTP_{int}]$ | $k_f = 0.139 s^{-1}$ | [21] |
| 21 | $G\alpha_i-GDP_i \rightarrow G\alpha_i-GDP$ | $k_f[G\alpha_i-GDP_i]$ | $k_f = 0.0011 s^{-1}$ | Est.<sup>2</sup> |

<sup>1</sup>Using experimental CDK5 activities [12], an initial guess was used for the rate, and then refined through fitting simulations to experimental data.
<sup>2</sup>Internalization rates were found by fitting simulations to experimental data.

**Table S4:** Reactions for **Module 3** (**Figure 3F**), outlining protein-protein interactions leading to the transibhibition of G*α_s_* by EGFR via GIV-GDI activation.

| # | Reaction | Reaction flux | Kinetic Parameters | Ref. |
| --- | --- | --- | --- | --- |
| 22 | $PLC-\gamma \rightarrow PLC-\gamma^*$ | $\frac{k_{cat}[EGF-EGFR_2][PLC-\gamma]}{K_M+[PLC-\gamma]}$<br>$-k_r[PLC-\gamma^*]$ | $k_{cat}=9.4 \times 10^{-4} \text{ s}^{-1}$, $K_M=1 \text{ } \mu M$,<br>$k_r=8.4 \times 10^{-5} \text{ s}^{-1}$ | [35],<br>Est.<sup>1</sup> |
| 23 | $PLC-\gamma \rightarrow PLC-\gamma^*$ | $\frac{k_{cat}[EGFR_2 \cdot GIV][PLC-\gamma]}{K_M+[PLC-\gamma]}$ | $k_{cat}=0.001 \text{ s}^{-1}$, $K_M=1 \text{ } \mu M$ | [11],<br>Est.<sup>2</sup> |
| 24 | $PIP2 \rightarrow IP3+DAG$ | $\frac{k_{cat}[PLC-\gamma^*][PIP2]}{K_M+[PIP2]}$ | $k_{cat}=14 \frac{\text{molecule}}{\mu m^2 \cdot \mu M \cdot s}$,<br>$K_M=5000 \frac{\text{molecule}}{\mu m^2}$ | [15] |
| 25 | $IP3 \rightarrow \text{Inositol}$ | $k_f[IP3]$ | $k_f=2.5 \text{ s}^{-1}$ | [15] |
| 26 | $PKC-\theta+DAG \rightarrow PKC-\theta^*$ | $k_f[PKC-\theta][DAG]$<br>$-k_r[PKC-\theta^*]$ | $k_f=0.1 \text{ } \mu M^{-1} \cdot \text{s}^{-1}$,<br>$k_r=1 \times 10^{-5} \text{ s}^{-1}$ | [36] |
| 27 | $DAG \rightarrow DAG_{deg}$ | $k_f[DAG]$ | $k_f=6.7 \times 10^{-4} \text{ s}^{-1}$ | [37] |
| 28 | $PKC-\theta^* \rightarrow PKC-\theta + DAG_{deg}$ | $k_f[PKC-\theta^*]$ | $k_f=0.028 \text{ s}^{-1}$ | Est.<sup>3</sup> |
| 29 | $EGFR_2 \cdot GIV \rightarrow GIV-GDI + EGF-EGFR^*_2$ | $\frac{k_{cat}[PKC-\theta][EGFR_2 \cdot GIV]}{K_M+EGFR_2 \cdot GIV}$ | $k_{cat}=1.528 \text{ s}^{-1}$,<br>$K_M=131 \frac{\text{molecule}}{\mu m^2}$ | [15] |
| 30 | $GIV-GEF \rightarrow GIV-GDI$ | $\frac{k_{cat}[PKC-\theta][GIV-GEF]}{K_M+[GIV-GEF]}$ | $k_{cat}=0.031 \text{ s}^{-1}$, $K_M=6 \text{ } \mu M$ | [15] |
| 31 | $GIV-GDI \rightarrow GIV$ | $k_f[GIV-GDI]$ | $k_f=2.3 \times 10^{-4} \text{ s}^{-1}$ | Est.<sup>1</sup> |

<sup>1</sup>Degradation rate set based on an assumed 20 to 30 min half life; then changed based on fitting
<sup>2</sup>Rate determined by immunoblot fitting
<sup>3</sup>Degradation rate of DAG assumed 10x faster when bound to PKC- $\theta$ , values were fit using a 10x faster rate as a starting point.

**Table S5:**
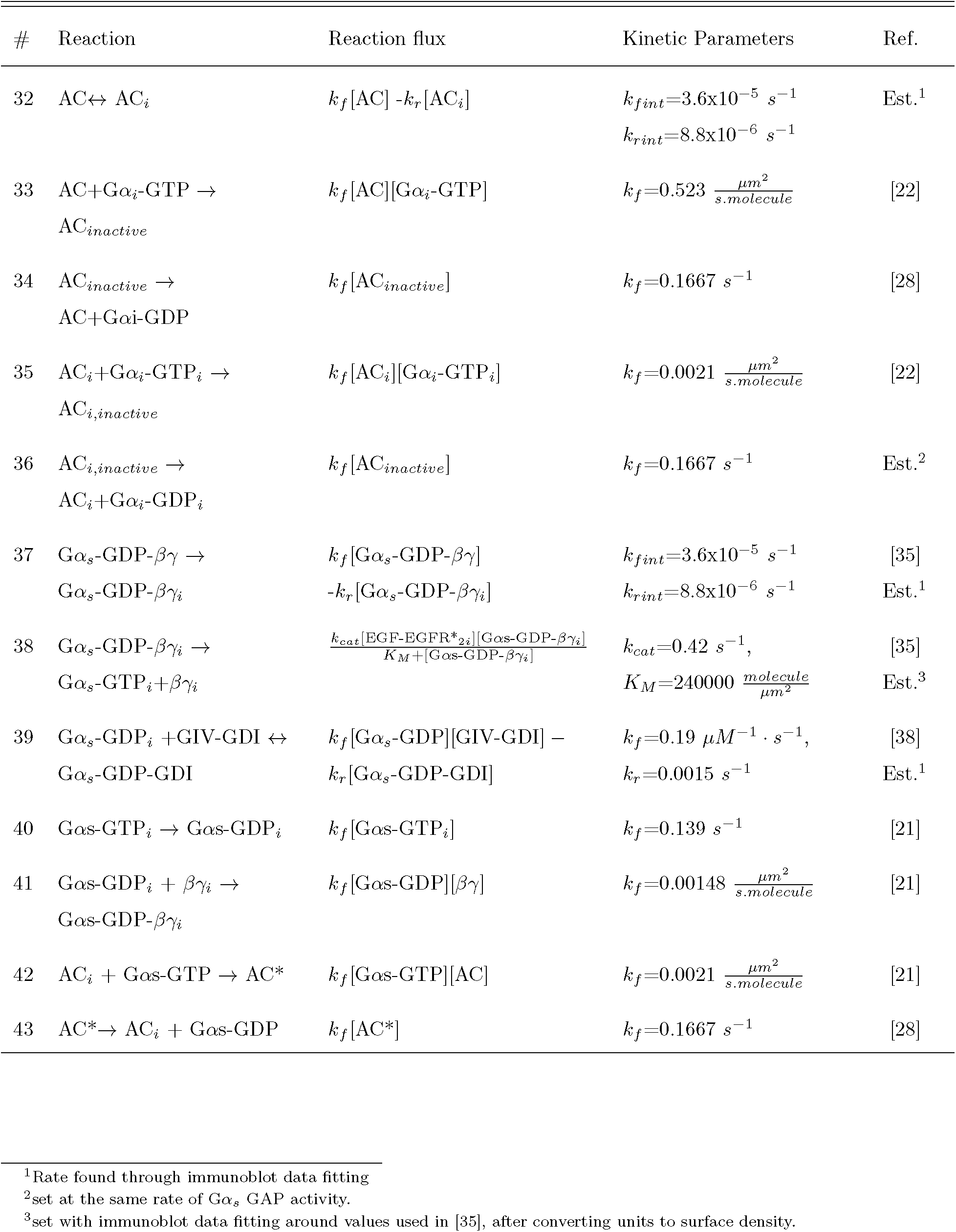
Reactions for **Module 4** (**Figure 4A**), outlining reactions for the activation and inhibition of cAMP.

**Table S6:**
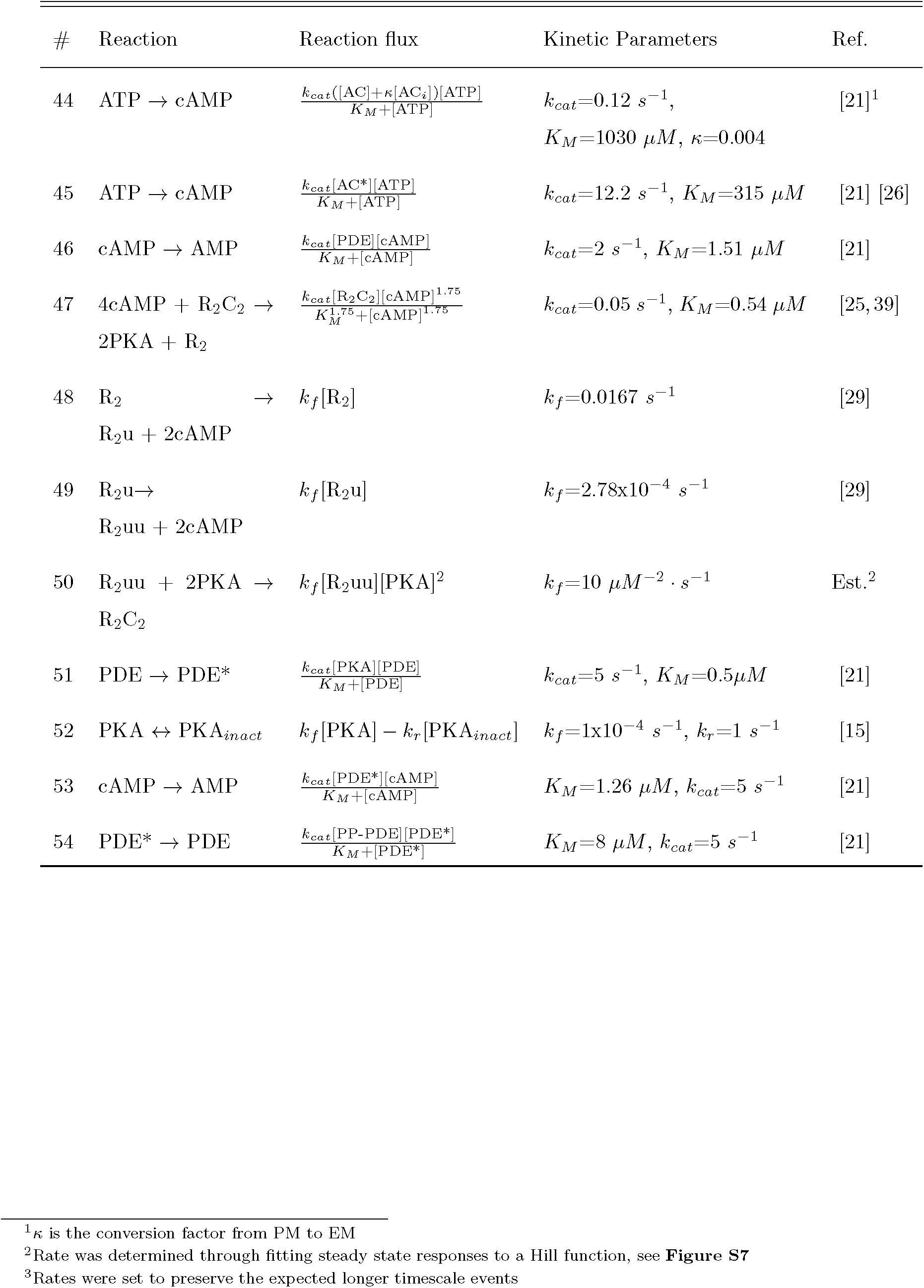
Reactions for **Module 4**(cont.)

**Table S7:** Reactions for the additional interactions modeling the effect of PKA and CDK5 phosphorylation of PDE.

| # | Reaction | Reaction flux | Kinetic Parameters | Reference |
| --- | --- | --- | --- | --- |
| 55 | $\text{PDE} \rightarrow \text{PDE}^*$ | $\frac{k_{cat}[\text{PKC}\theta^*][\text{PDE}]}{K_M + [\text{PDE}]}$ | $K_M=0.89 \mu M, k_{cat}=15 s^{-1}$ | [31] |
| 56 | $\text{PDE} \rightarrow \text{PDE}^*$ | $\frac{k_{cat}[\text{CDK}^*][\text{PDE}]}{K_M + [\text{PDE}]}$ | $K_M=0.5 \mu M, k_{cat}=1 s^{-1}$ | [30] |

**Table S8:**
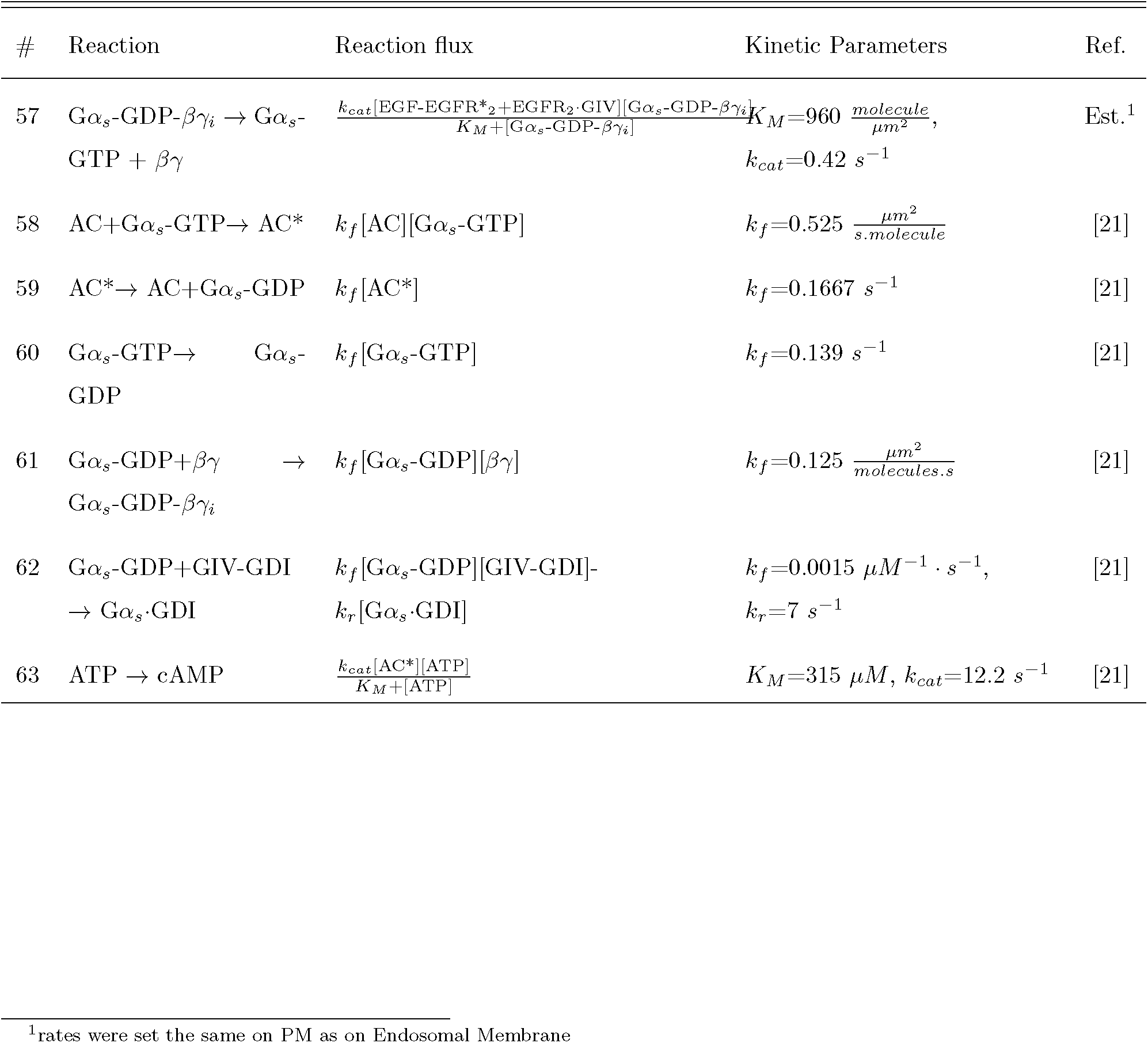
Reactions for rapid production of cAMP at the PM (blue line in **Figure 7A**) using the dynamics shown in [40].

**Table S9:** Initial conditions for components (components not listed have zero initial conditions)

| Species | Initial concentration | Compartment | Notes and References |
| --- | --- | --- | --- |
| EGFR | 240 molecule.$\mu m^{-2}$ | PM | [4, 41] |
| EGF | 0.05 $\mu M$ | EC | Experiment stimulation |
| PLC-γ | 0.1 $\mu M$ | Cytosol | Assumed |
| PKC-θ | 0.1 $\mu M$ | Cytosol | Assumed |
| ATP | 5000 $\mu M$ | Cytosol | [21] |
| cAMP | 0.0045 $\mu M$ | Cytosol | basal SS<sup>1</sup> |
| PKA | 0.0066 $\mu M$ | Cytosol | basal SS<sup>1</sup> |
| PDE | 0.345 $\mu M$ | Cytosol | basal SS<sup>1</sup> |
| R2C2 | 0.0967 $\mu M$ | Cytosol | basal SS<sup>1</sup> |
| AMP | 1000 $\mu M$ | Cytosol | [21] |
| PDEp | 0.055 $\mu M$ | Cytosol | basal SS<sup>1</sup> |
| PPPDE | 0.2 $\mu M$ | Cytosol | Assumed |
| GIV | 0.831 $\mu M$ | Cytosol | Set such that $GIV_{total} = 1 \mu M$ |
| GIV-GEF | 0.05 $\mu M$ | Cytosol | Assumed; immunoblot values |
| GIV-GDI | 0.1 $\mu M$ | Cytosol | Assumed; immunoblot values |
| CDK5 | 0.05 $\mu M$ | Cytosol | Assumed; [12] |
| R2 | 6x10<sup>-5</sup> $\mu M$ | Cytosol | basal SS<sup>1</sup> |
| R2u | 0.004 $\mu M$ | Cytosol | basal SS<sup>1</sup> |
| R2uu | 0.0025 $\mu M$ | Cytosol | basal SS<sup>1</sup> |
| p35 | 957 molecule.$\mu m^{-2}$ | PM | Set such that $p35_{total} = 1 \mu M$ |
| G$\alpha_i$-GDP | 48 molecule.$\mu m^{-2}$ | PM | Set to exp. immunoblot value. |
| AC | 30 molecule.$\mu m^{-2}$ | PM | Average of [21] and [26] |
| PIP2 | 9570 molecule.$\mu m^{-2}$ | PM | Assumed |
| EGFR·GIV·G$\alpha_i$ | 0.01 molecule.$\mu m^{-2}$ | PM | experimental immunoblot value |
| G$\alpha_s$-GDP-$\beta\gamma$ | 1500 molecule.$\mu m^{-2}$ | PM | Calculated from AC to G$\alpha_s$ ratio [27] |
| G$\alpha_s$·GIV-GDI | 75 molecule.$\mu m^{-2}$ | Endosomal membrane | experimental immunoblot value |

<sup>1</sup>All basal steady states are set by running basal AC stimulation until a stable cAMP, PKA, and PDE response were received.

**Table S10:** Sensitivity analysis of EGFR·GIV·G*α_i_* complex with respect to initial conditions. The colors indicate the sensitivity to the respective parameter; red indicates that the EGFR·GIV·G*α_i_* complex is sensitive to changes in the initial concentration of the corresponding parameter (i.e. sensitivity index greater than 1) and blue indicates that the EGFR·GIV·G*α_i_* complex is partially sensitive to changes in the initial concentration of the corresponding parameter (i.e. sensitivity index greater than 0.5) over the time course of signaling. Sensitivity is shown at 5, 15, 30, and 60 min intervals.

| Initial Parameter | 5 min | 15 min | 30 min | 60 min |
| --- | --- | --- | --- | --- |
| G$\alpha_i$-GDP | blue | red | red | red |
| GIV | white | blue | blue | red |
| PLC-γ | white | blue | blue | blue |
| Cdk5 | white | white | white | blue |
| PIP<sub>2</sub> | white | white | white | blue |
| PLCγ | white | blue | red | red |

**Table S11:** Sensitivity analysis of cAMP with respect to initial conditions. The colors indicate the sensitivity to the respective parameter; red indicates that cAMP is sensitive to changes in the initial concentration of the corresponding parameter (i.e. sensitivity index greater than 1) and blue indicates that cAMP is partially sensitive to changes in the initial concentration of the corresponding parameter (i.e. sensitivity index greater than 0.5) over the time course of signaling. Sensitivity is shown at 5, 15, 30, and 60 min intervals.

| Initial Parameter | 5 min | 15 min | 30 min | 60 min |
| --- | --- | --- | --- | --- |
| AC | red | red | red | blue |
| G$\alpha_i$-GDP | red | red | blue | white |
| GIV | white | blue | white | white |
| PDE | blue | blue | blue | blue |

<sup>0</sup>The G $\alpha_s$ ·GIV-GDI complex is not sensitive to any initial condition values and is not given an initial sensitivity table.

**Table S12:** Sensitivity analysis of EGFR·GIV·G*α_i_* complex with respect to the model kinetic parameters. The colors indicate the sensitivity to the respective parameter; red indicates that the EGFR·GIV·G*α_i_* complex is sensitive to changes in the value of the corresponding parameter (i.e. sensitivity index greater than 1) and blue indicates that the EGFR·GIV·G*α_i_* complex is partially sensitive to changes in the value of the corresponding parameter (i.e. sensitivity index greater than 0.5) over the time course of signaling. Sensitivity is shown at 5, 15, 30, and 60 min intervals. The index in the square brackets refer to the reaction number.

| Parameter[reaction] | 5 min | 15 min | 30 min | 60 min |
| --- | --- | --- | --- | --- |
| $k_{cat}$[R14] | | | | blue |
| $k_{cat}$[R22] | | blue | blue | red |
| $k_{cat}$[R24] | | blue | red | red |
| $k_{cat}$[R29] | | blue | red | red |
| $k_f$[R1] | blue | | | |
| $k_f$[R4] | | | blue | red |
| $k_f$[R15] | blue | blue | blue | red |
| $k_f$[R16] | blue | | | |
| $k_f$[R17] | blue | blue | blue | blue |
| $k_f$[R28] | | | blue | red |
| $K_m$[R14] | | | | blue |
| $K_m$[R22] | | | blue | blue |
| $K_m$[R24] | | | | blue |
| $K_m$[R29] | | blue | red | red |
| $k_r$[R14] | | blue | red | red |

**Table S13:**
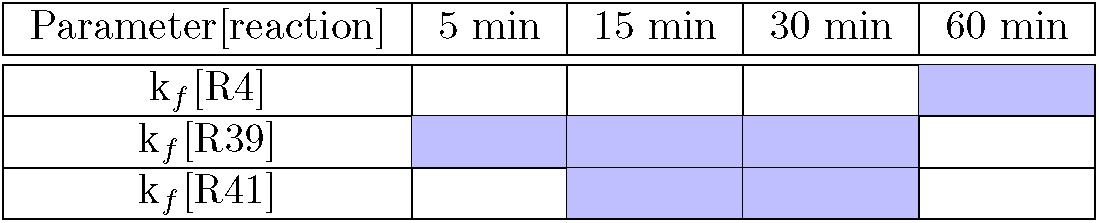
Sensitivity analysis of G*α_s_*·GIV-GDI complex with respect to the model kinetic parameters. The colors indicate the sensitivity to the respective parameter; red indicates that the G*α_s_*·GIV-GDI complex is sensitive to changes in the value of the corresponding parameter (i.e. sensitivity index greater than 1) and blue indicates that the G*α_s_*·GIV-GDI complex is partially sensitive to changes in the value of the corresponding parameter (i.e. sensitivity index greater than 0.5) over the time course of signaling. Sensitivity is shown at 5, 15, 30, and 60 min intervals. The index in the square brackets refer to the reaction number.

**Table S14:**
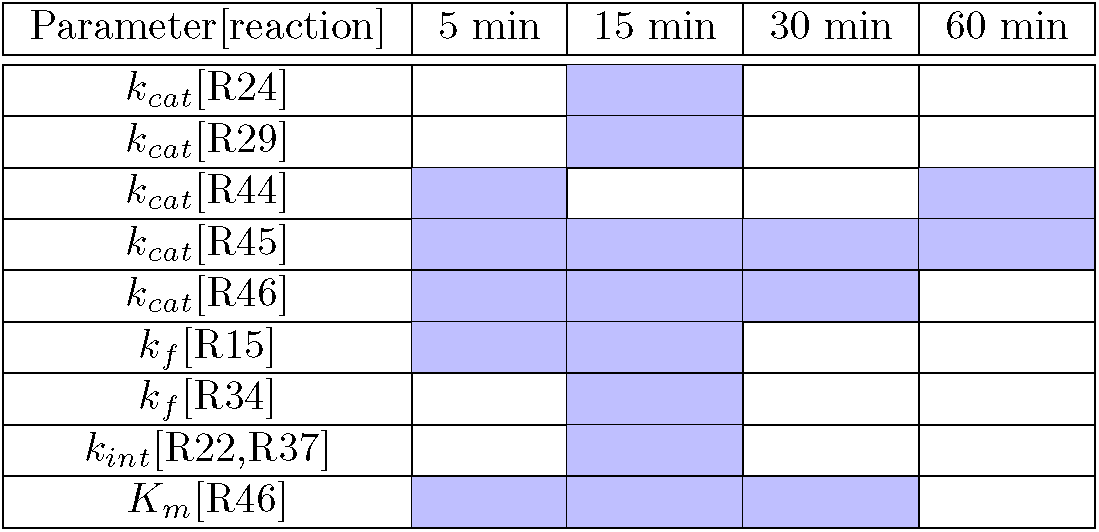
Sensitivity analysis of cAMP with respect to the model kinetic parameters. The colors indicate the sensitivity to the respective parameter; red indicates that the cAMP production is sensitive to changes in the value of the corresponding parameter (i.e. sensitivity index greater than 1) and blue indicates that cAMP production is partially sensitive to changes in the value of the corresponding parameter (i.e. sensitivity index greater than 0.5) over the time course of signaling. Sensitivity is shown at 5, 15, 30, and 60 min intervals. The index in the square brackets refer to the reaction number.

### 5 Supplementary Figures

**Figure S1:**
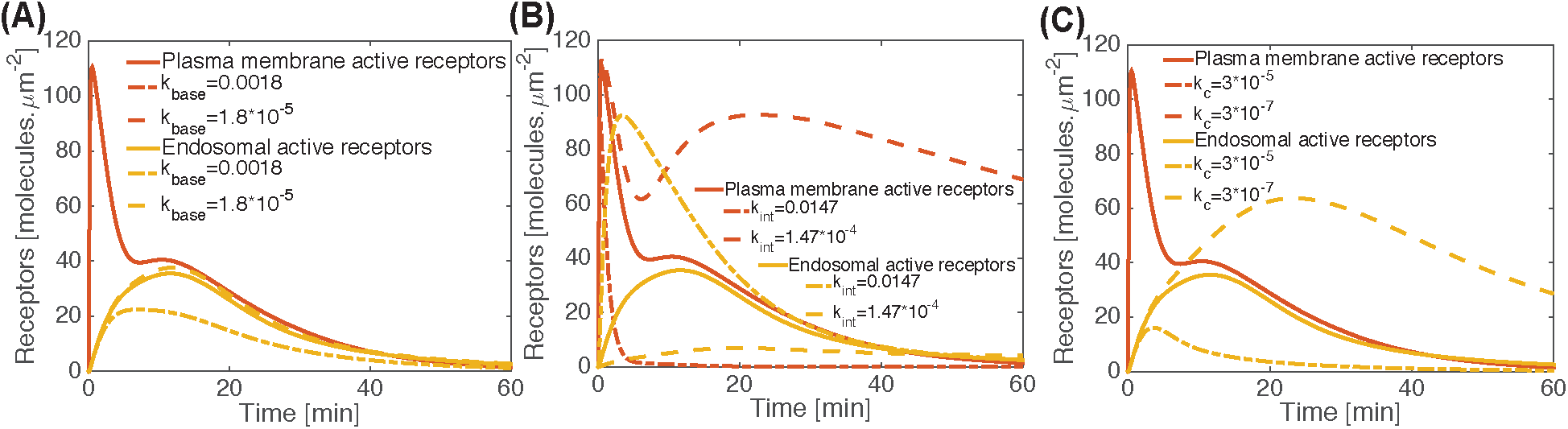
Supplementary figure for Figure 2 – Dynamics of different pools of EGFR. Simulations are shown for the dynamics of the PM (red line) and endosomal (yellow line) pools of EGFR computed over 1 h based on network Module 1. Parameter variations were conducted for values one order of magnitude above and below the control value for **(A)** basal degradation rate (reaction number 9 in **Table S2**), **(B)** receptor internalization rate (reaction number 4 in **Table S2**), and **(C)** rate of G*α_s_*-GDP dependent catalytic degradation of EGFR (reaction number 9 in **Table S2**). The solid line shows the value used in the control model, the dot dashed lines represent a ten-fold increase the value of the parameter from the control value and the dashed lines represent a ten-fold decrease in the value of the kinetic parameter from the control value. Variation of the basal degradation rate of EGFR doesn’t affect the PM receptors but proportionally affects the endosomal receptor pool (**A**); an increase in the basal degradation rate decreases the endosomally active receptors (Reaction 9, **Table S2**). On the other hand, variation in the rate of internalization of EGFR affects both the PM and endosome pool of receptors (Reaction 4, **Table S2**). An increase in the rate of internalization of EGFR leads to a rapid decrease in the PM receptor pool with a corresponding rapid increase in the endosome pool of receptors (**B**). Variation of the G*α_s_*-GDP-dependent catalytic degradation rate of EGFR (Reaction 9, **Table S2**) affects the endosomal receptor pool proportionally, with no discernible effect on the PM receptor pool (**C**).

**Figure S2:**
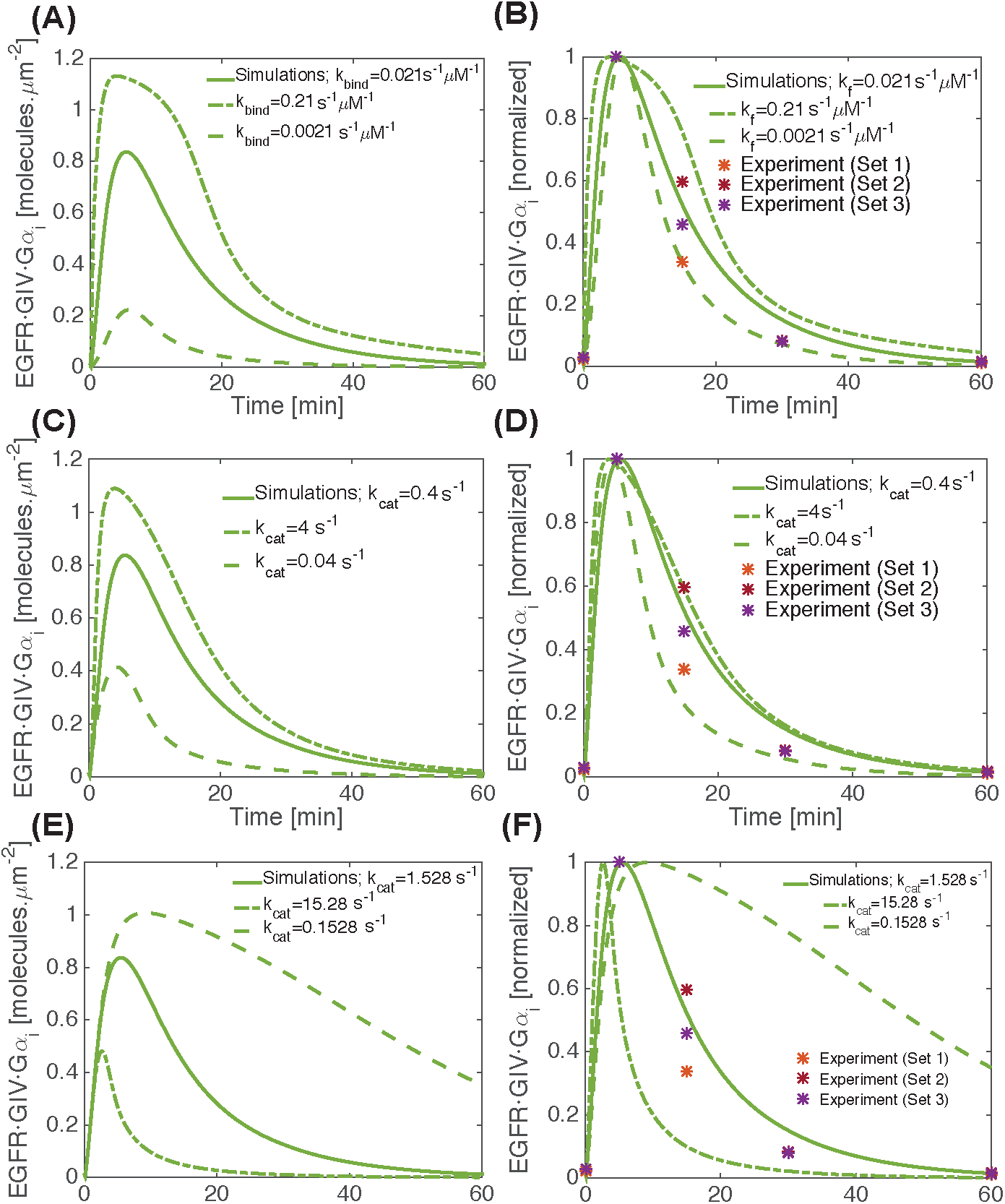
Supplementary figure for Figure 2 – Impact of model parameters on the dynamics of the formation of the EGFR·GIV·G*α_i_* complexes downstream of EGFR activation. **(A-F)** Effect of different kinetic parameters on the dynamics of the EGFR·GIV·G*α_i_* complex formation was tested by changing the parameter of interest to one order of magnitude above (dot dashed lines) and one order of magnitude below (dashed line) the control value (solid line) used in the model. **(A)** and **(B)** show the effect of binding rate constant of GIV-GEF to ligand-bound, dimerized EGFR (reaction number 15 in **Table S3**) on the dynamics of the formation of the EGFR·GIV·G*α_i_* complex. Although the density of the EGFR·GIV·G*α_i_* complex is affected by this rate constant **(A)**, the normalized complex density shows good agreement with experiment. **(C)** and **(D)** show the effect of CDK5-mediated phosphorylation of GIV to GIV-GEF on the formation of the EGFR·GIV·G*α_i_* complex (*k_cat_* reaction number 14 in **Table S3**). Even though the density of the EGFR·GIV·G*α_i_* complex is affected by the *k_cat_* **(C)**, the normalized values are in good agreement with experiment. **(E)** and **(F)** Simulations display the effect of PKC-*θ*-mediated phosphorylation at S1689 for GIV-GEF·EGFR, resulting in conversion of GIV-GEF to GIV-GDI (reaction number 29 in **Table S4**). Changing this k_cai_ changes the dynamics of the EGFR·GIV·G*α_i_* complex formation such that a decrease in this rate constant leads to a prolonged lifetime of the complex and this effect is seen both in the number density of the complex **(E)** and in the normalized data compared against experiments **(F)**.

**Figure S3:**
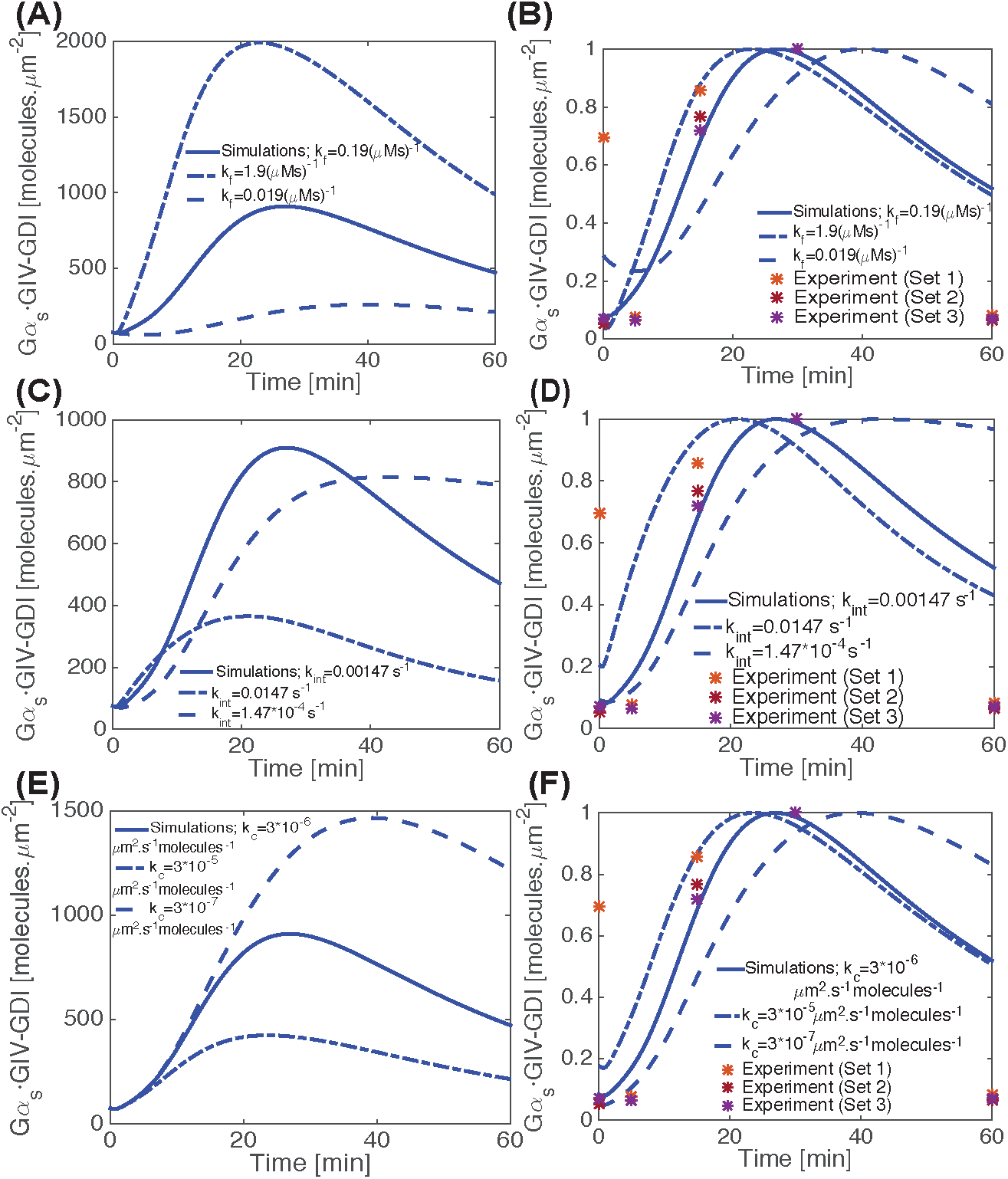
Supplementary figure for Figure 3 — Dynamics of G*α_s_*·GIV-GDI complex formation downstream of EGFR and the effect of different kinetic parameters. **(A-F)** Effect of different kinetic parameters on the dynamics of the G*α_s_*·GIV-GDI complex formation was tested by changing the parameter of interest to one order of magnitude above (dot dashed lines) and one order of magnitude below (dashed line) the control value (solid line) used in the model. (**A**-**B**) Variation of the binding rate of GIV-GDI binding rate to G*α_s_*-GDP (reaction number 39 in **Table S5**) affects both the density of the bound G*α_s_*·GIV-GDI molecules (**A**) and the temporal dynamics **(B)**. Reducing the value of this rate constant shifted the peak time of complex formation to the right, an effect that is clearly visualized when comparing the normalized data (**B**). (**C**-**D**) Varying the internalization rate of dimerized EGFR, from the PM to the endosomal compartment (reaction number 4 in **Table S2**) dramatically changes the dynamics of the G*α_s_*·GIV-GDI complex formation both in terms of the surface density (**C**) and when the normalized quantities are compared against experiment (**D**). Faster internalization rates of EGFR lowered the density of the complex (**C**) and the complexes were assembled earlier than observed in experiments (**D**). Reducing the rate of EGFR internalization, on the other hand, also lowered the density of G*α_s_*·GIV-GDI complexes, and they were assembled later. (**E**-**F**) The dynamics of the G*α_s_*·GIV-GDI complex formation are affected by the catalytic degradation rate of internalized dimerized EGFR, which is enabled by G*α_s_*-GDP [7] (reaction number 9 in **Table S2**). The effect of changing this parameter was proportional on both the membrane density of the G*α_s_*·GIV-GDI complex (**E**) and affected the peak time and dynamics of the complex formation **F**. Because the degradation of internalized EGFR requires endosomal maturation that is enhanced by G*α_s_*-GDP [7], increasing the rate of G*α_s_*-GDP dependent endosome maturation and EGFR degradation decreased the G*α_s_*-GDI density and increased the rate of the G*α_s_*·GIV-GDI complex formation, whereas decreasing the rate of the G*α_s_*-GDP-mediated EGFR degradation increased the density of the complex formation and slowed down the process.

**Figure S4:**
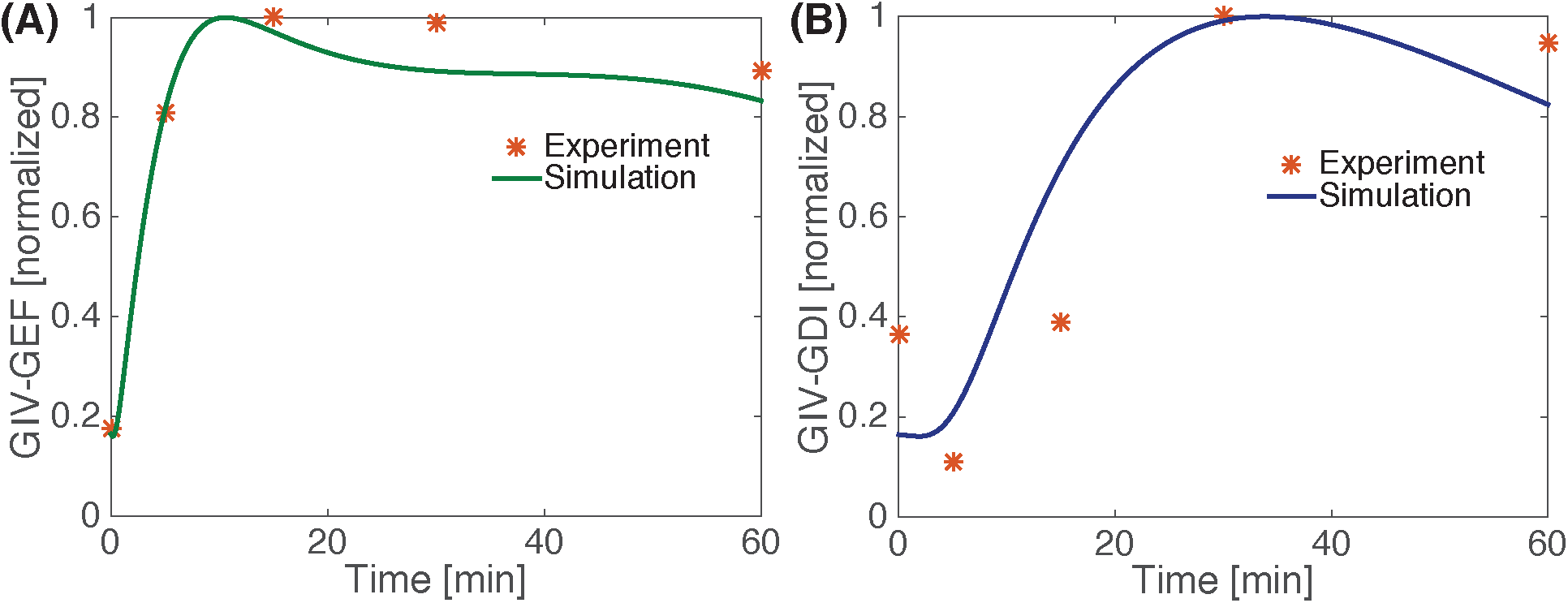
Supplementary figure for Figures 2 and 3 - Parameter estimation and validation of cytosolic GEF and GDI concentrations. (**A**) Comparison of simulation and experimental data for GIV-GEF. (**B**) Comparison of simulation and experimental data for GIV-GDI. The concentration of GIV-GEF and GIV-GDI was normalized to its peak value and compared against experimental data (*). Experimental data were obtained from Figures 1D and S1 of [2], in which protein-protein interaction assays were performed using lysates of cells responding to EGF.

**Figure S5:**
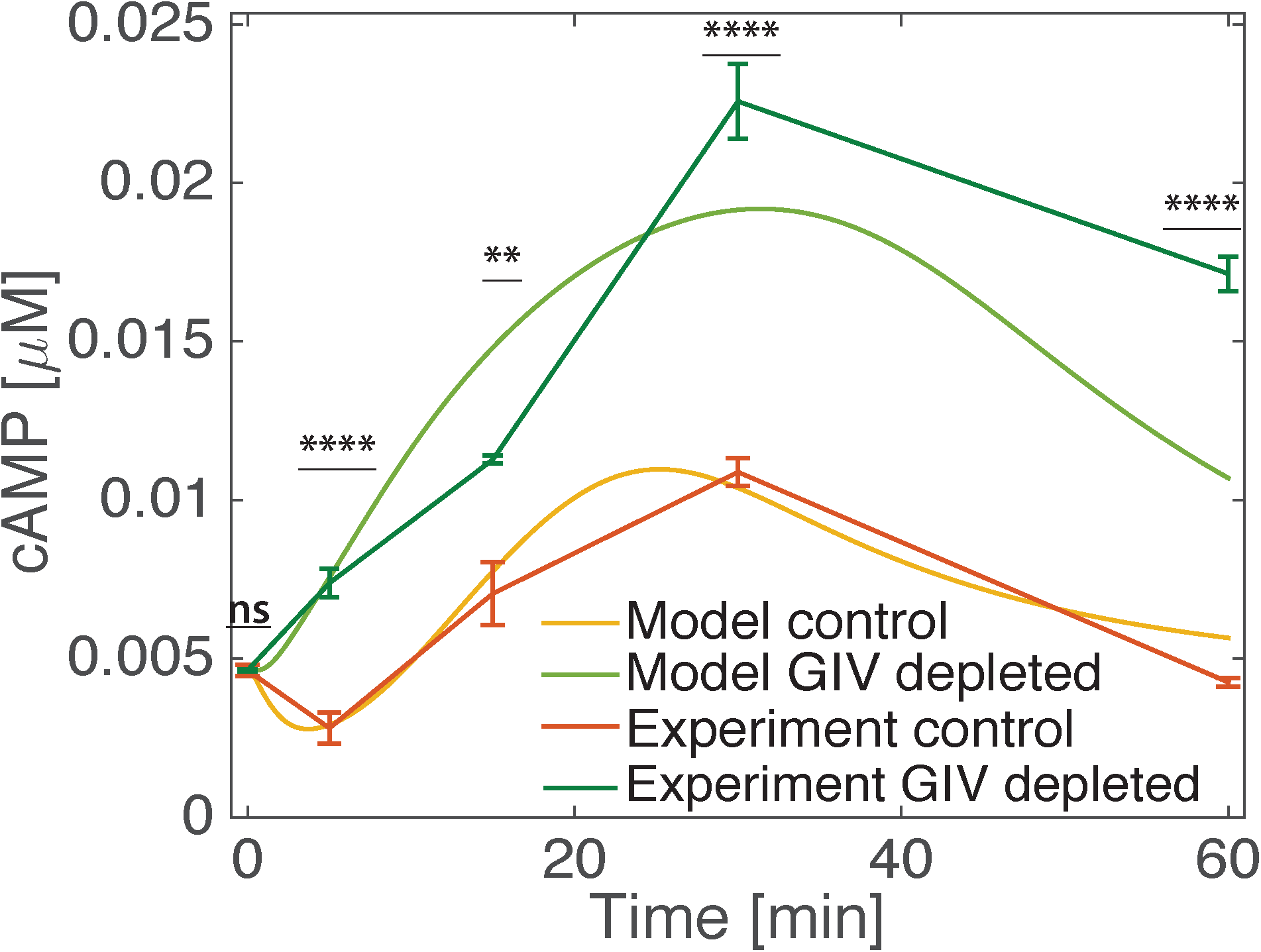
Supplementary figure for Figure 4 - Parameter estimation and validation of the cAMP concentration. Comparison of cAMP time course from simulations and experiments. Simulations of dynamics of the production of cAMP based on the network modules was performed. The concentration of cAMP was normalized to its initial value and compared experimental data in which control or GIV-depleated (shGIV) HeLA cells were serum starved (0.2% FBD, 16h) prior to stimulation with 50nM EGF for the indicated time points. cAMP produced in response to EGF was measured by radioimmunoassay (RIA) as detailed in the main text ‘Materials and method’ section. Error bars indicated mean S.D of three independent experiments. ns= not significant; ^**^p=0.01,^****^p=0.0001. cAMP was normalized against the initial value to ensure the pre-stimulation steady state concentrations of cAMP are satisfied.

**Figure S6:**
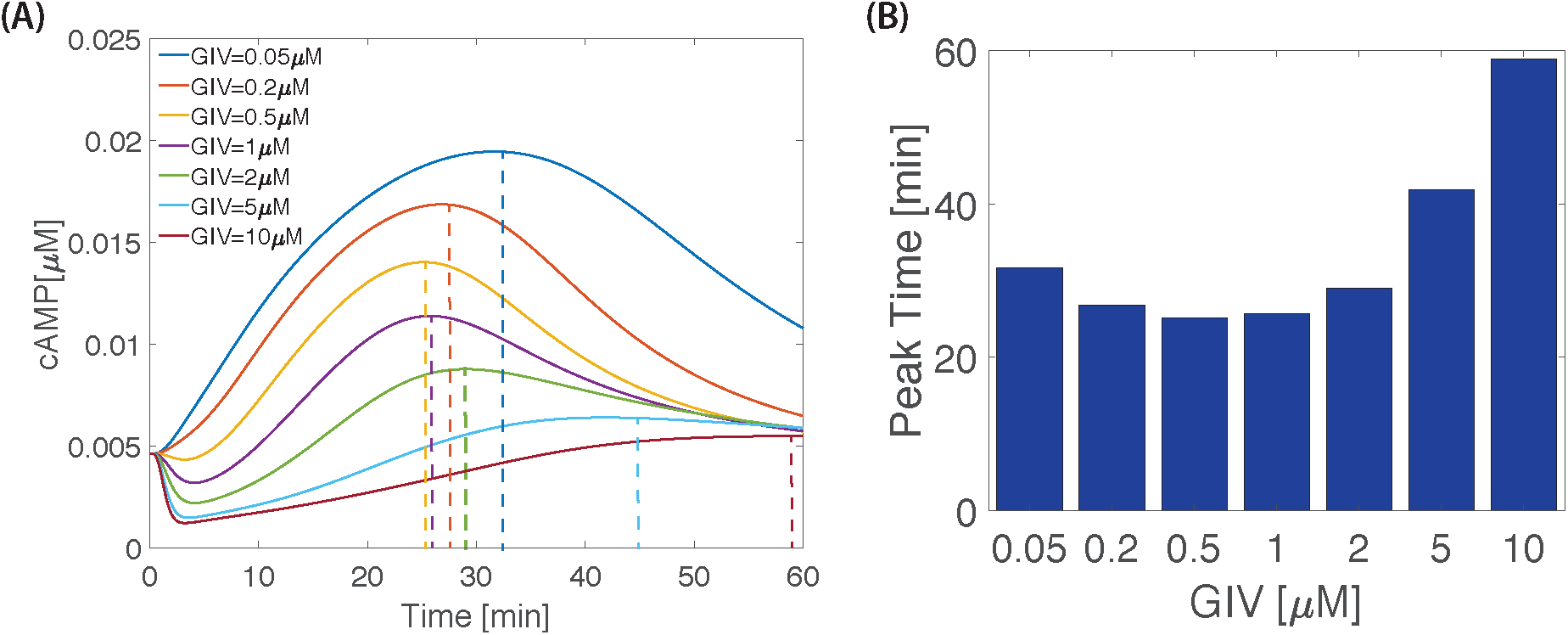
Supplementary figure for Figure 4 - GIV concentration affects peak cAMP time. Simulations were performed by varying the concentration of GIV with values ranging from 0.05 to 10 *μM*, the peak times for cAMP were then extracted (**A**). These times were then plotted on a bar graph (**B**). cAMP peak times were found to shorten and then lengthen with increasing GIV concentrations in a nonlinear manner. The initial, low GIV, peak time decrease is due to the action of G*α_s_*·GIV-GDI shortening the timescale of G*α_s_* activation. While the later, high GIV, increase is due to the cAMP the action of *Gai* inhibition becoming more prevalent over the G*α_s_* generation of cAMP.

**Figure S7:**
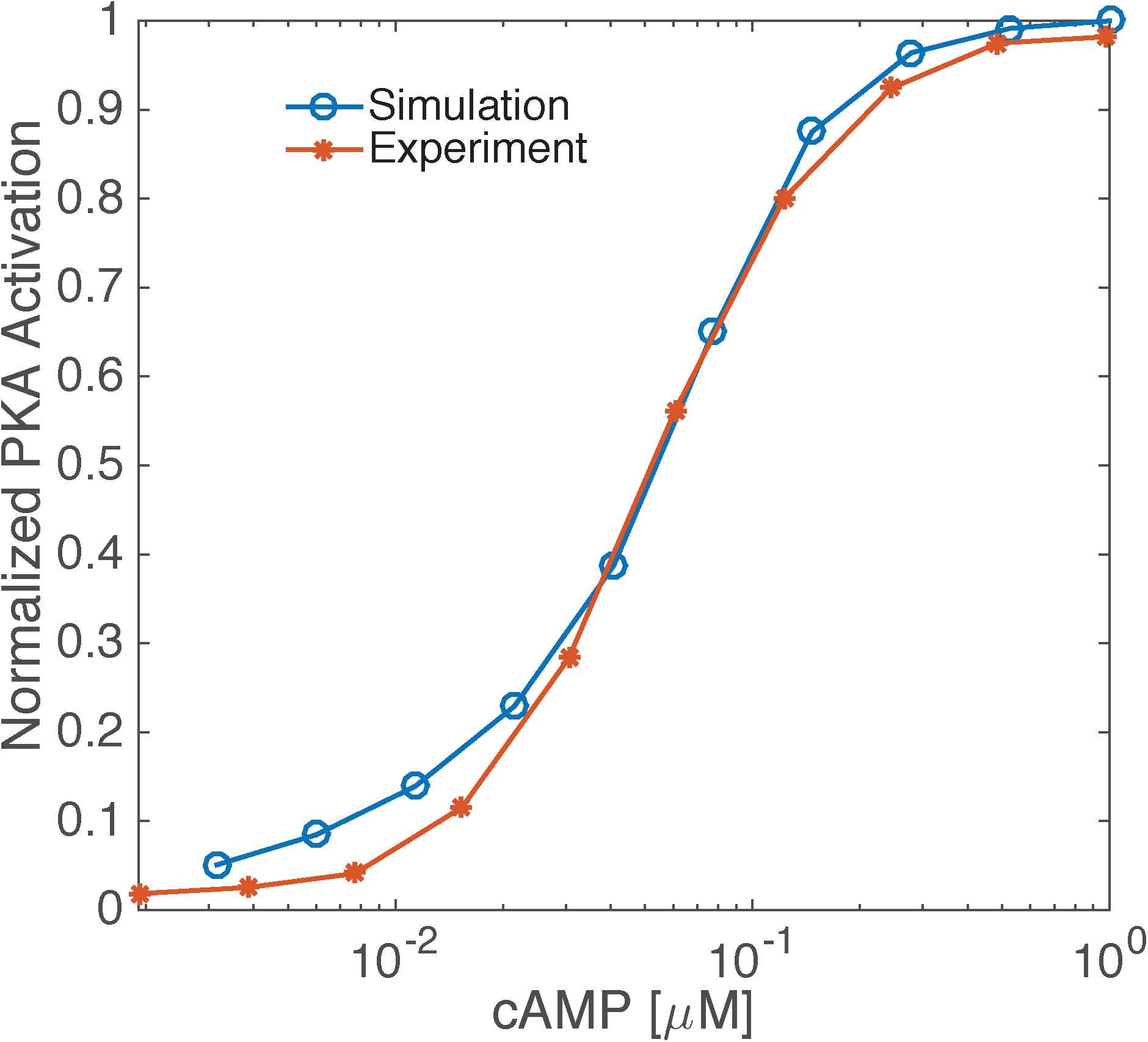
Supplementary figure for Figure 4 – Parameter estimation and validation of cAMP-PKA interactions. Dose response curve for PKA activation as a function of cAMP concentration was calculated from simulations and compared against previously published experiments. The red starred line shows the normalized PKA activation from [25] and the blue line shows the model PKA activation as a function of cAMP concentration. Experimental data from Bruystens *et al*. [25] was fit to a Hill function to obtain a Hill coefficient of n=1.73 and an EC_50_=54 nM. These values were used in our model (**Module 4, Reactions 47-50**) to obtain a good qualitative agreement between simulations and experimental data for cAMP-mediated activation of PKA.

**Figure S8:**
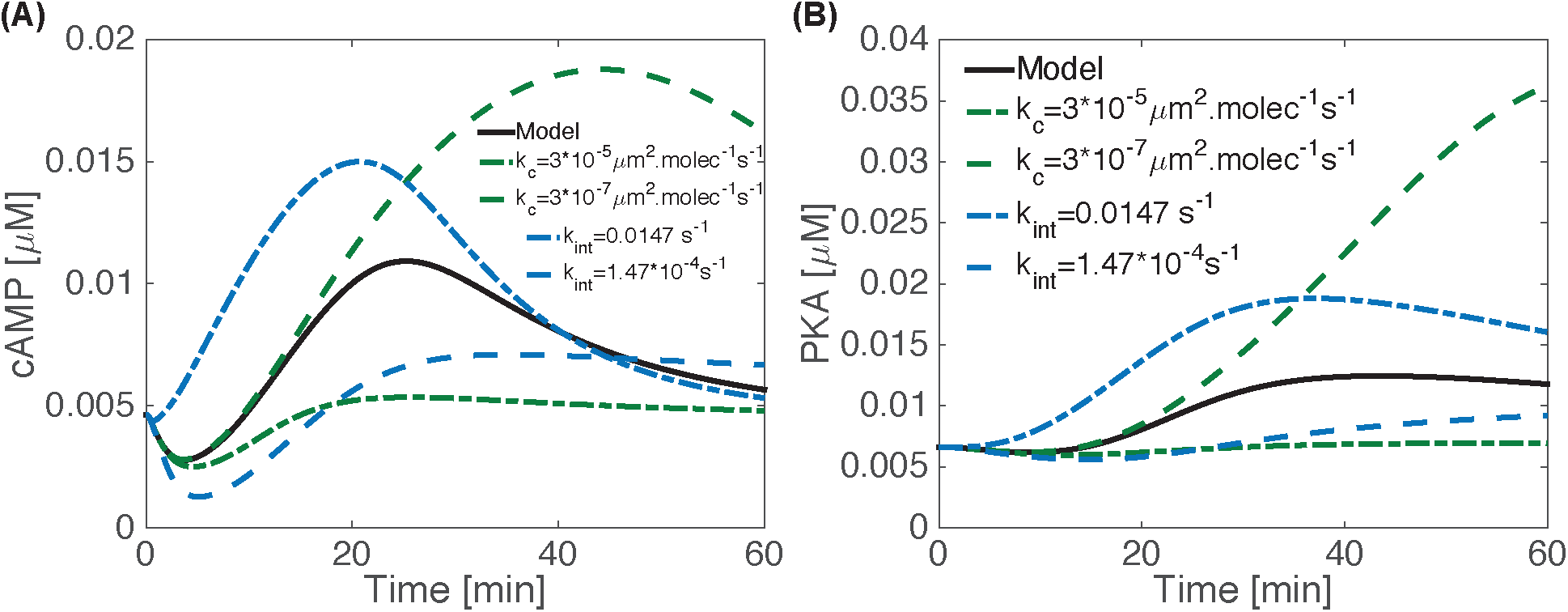
Supplementary figure for Figure 4 – Effect of EGFR dynamics on cAMP dynamics. Simulations display the impact of variation of receptor internalization and catalytic rate on cAMP **(A)** and PKA **(B)**. An increase in the internalization rate of EGFR leads to the rapid increase in cAMP and loss of the early G*α_i_*-dependent reduction in cAMP concentration (**A**). On the other hand, a decrease in the internalization rate of EGFR leads to a reduction in cAMP concentration without an appreciable increase in cAMP concentration. An increase in the EGFR degradation rate diminishes cAMP concentration because of reduced G*α_s_* activation in the endosomes and a corresponding decrease in the degradation rate leads to an increase in the cAMP concentration (**A**). Neither of these conditions affect the early reduction of cAMP due to G*α_i_*. PKA mimics the response of cAMP (**B**) by virtue of being downstream effectors of cAMP. Increasing endosomal stimulation times, by reducing catalytic degradation rate or by increasing internalization rates, resulted in an amplified PKA response.

**Figure S9:**
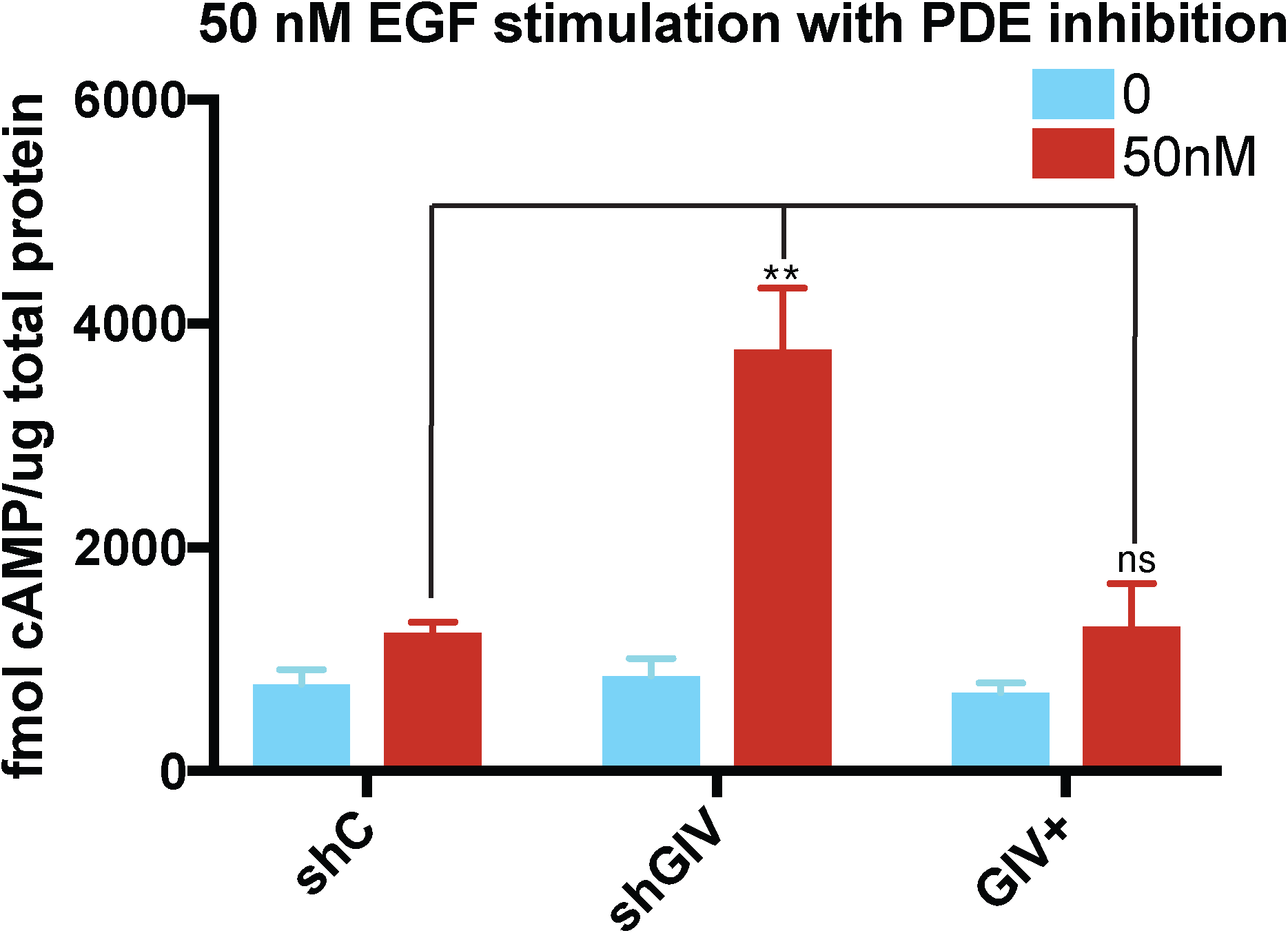
Supplementary figure for Figure 4 – GIV is required for suppressing EGF-triggered cAMP production in HeLa cells. Control (shC) or GIV-depleted (shGIV) HeLa cells or GIV-depleted cells rescued with shRNA-resistant GIV-WT (GIV+) were serum starved (0.2% FBS, 16h), treated with 200 *μ*M IBMX (20 min) prior to stimulation with 50 nM EGF for 60 min cAMP produced in response to EGF was measured by radioimmunoassay (RIA) as detailed in ‘Materials and methods’. Bar graphs compare the cAMP levels before (0, blue) and after EGF stimulation (50nM, red). Error bars indicate mean ± S.D. of three independent experiments.

**Figure S10:**
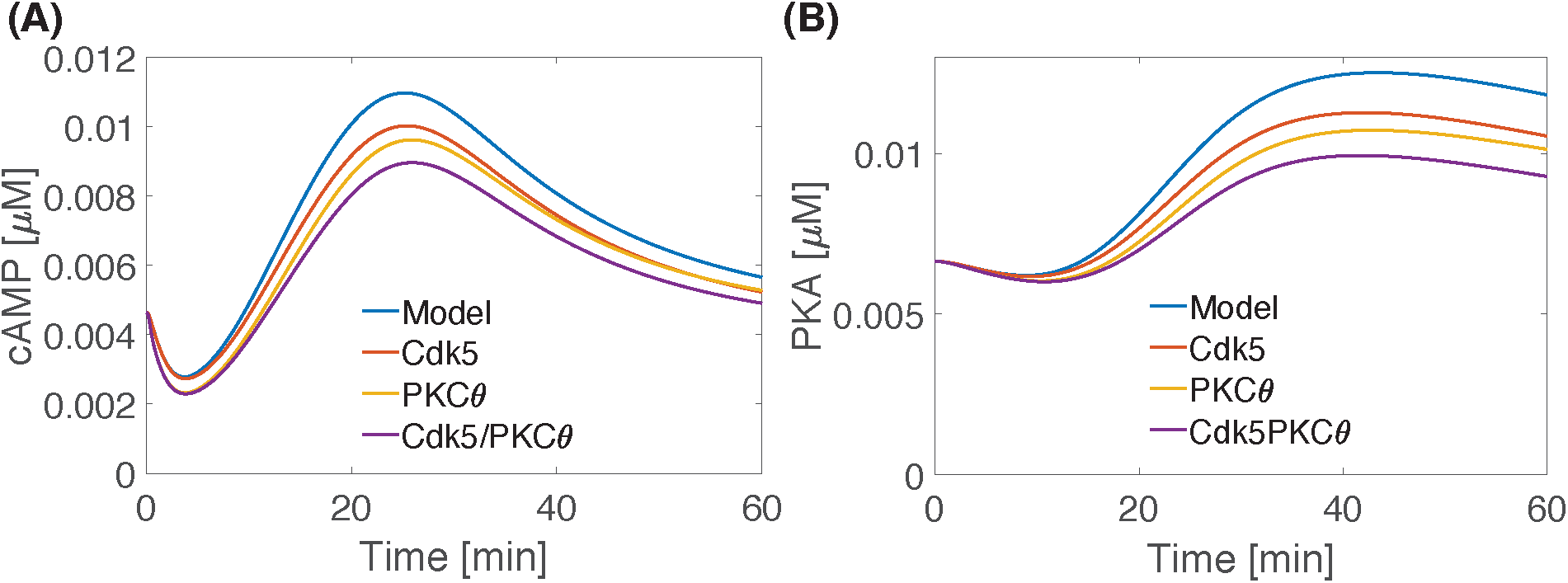
Supplementary figure for Figure 4 – Effect of PKC-*θ* and CDK5 phosphorylation on PDE phosphorylation. Simulations of the impact of phosphorylation of PDE by PKC-*θ* and CDK5 (reactions are shown in **Table S7**) on (**A**)cAMP and (**B**) PKC-*θ* are shown. The control values of cAMP and PKC-*θ* without accounting for either of the two feedforward interactions are shown in blue. Inclusion of the feedforward loops. i.e., phosphorylation of PDE by either CDK5 alone (red) or PKC-*θ* alone (yellow), or both (purple) are also displayed. When both CDK5 and PKC-*θ* feedback loops are taken into account (purple), PDE activity appears to be enhanced because cellular cAMP dynamics are significantly dampened.

**Figure S11:**
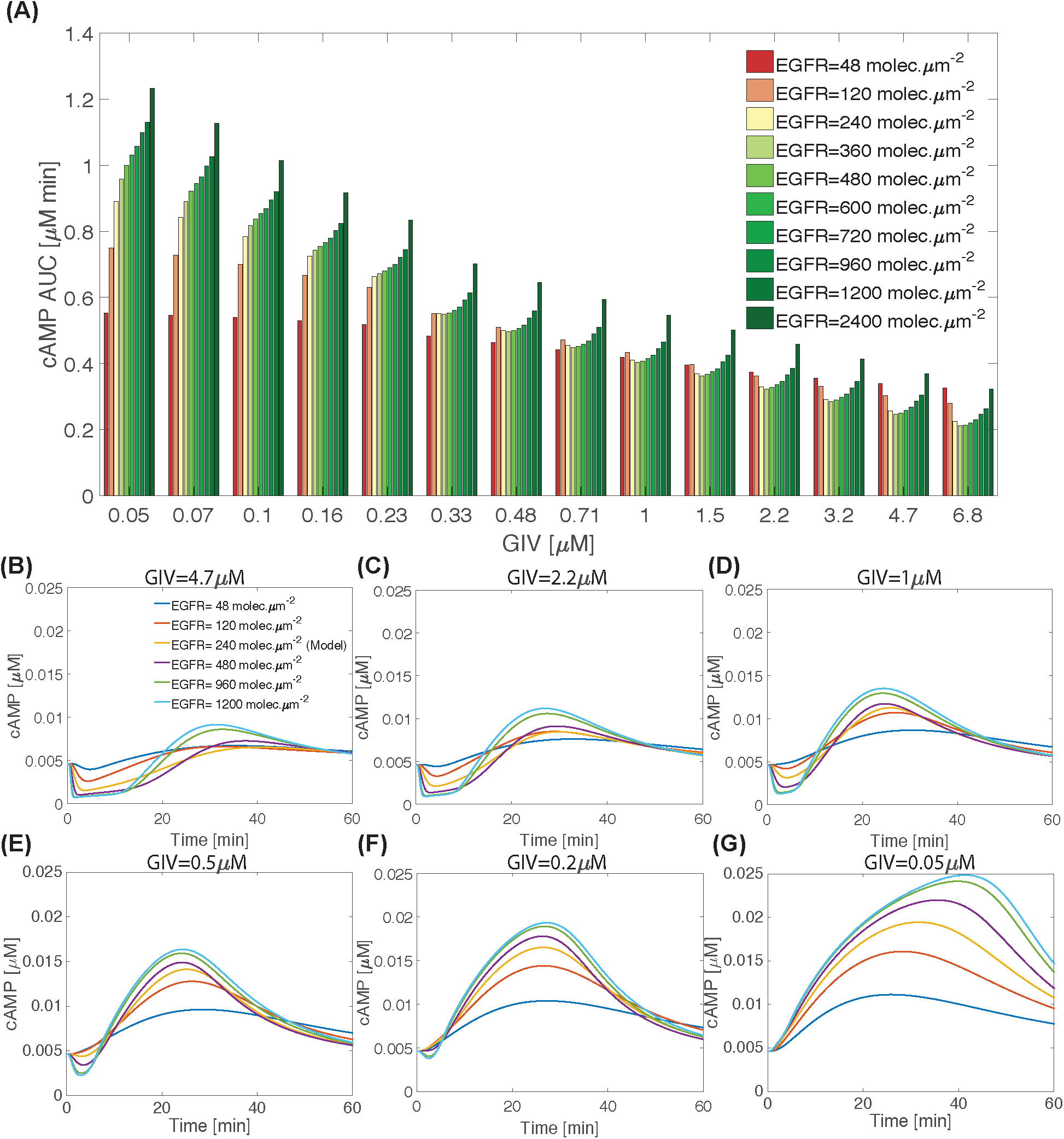
Supplementary figure for Figure 5 – Impact of varying EGFR and GIV concentrations on cellular levels of cAMP. (**A**) cAMP AUC, computed at 1 h is shown for different values of GIV and EGFR. Time-course of cAMP for various EGFR concentrations at (**B**) GIV=0.05 *μM*, (**C**) GIV=0.2 *μM*, (**D**) GIV=0.5 *μM*, (**E**) GIV=1 *μM*, (**F**) GIV=2 *μM*, (**G**) GIV=5 *μM* are shown.

**Figure S12:**
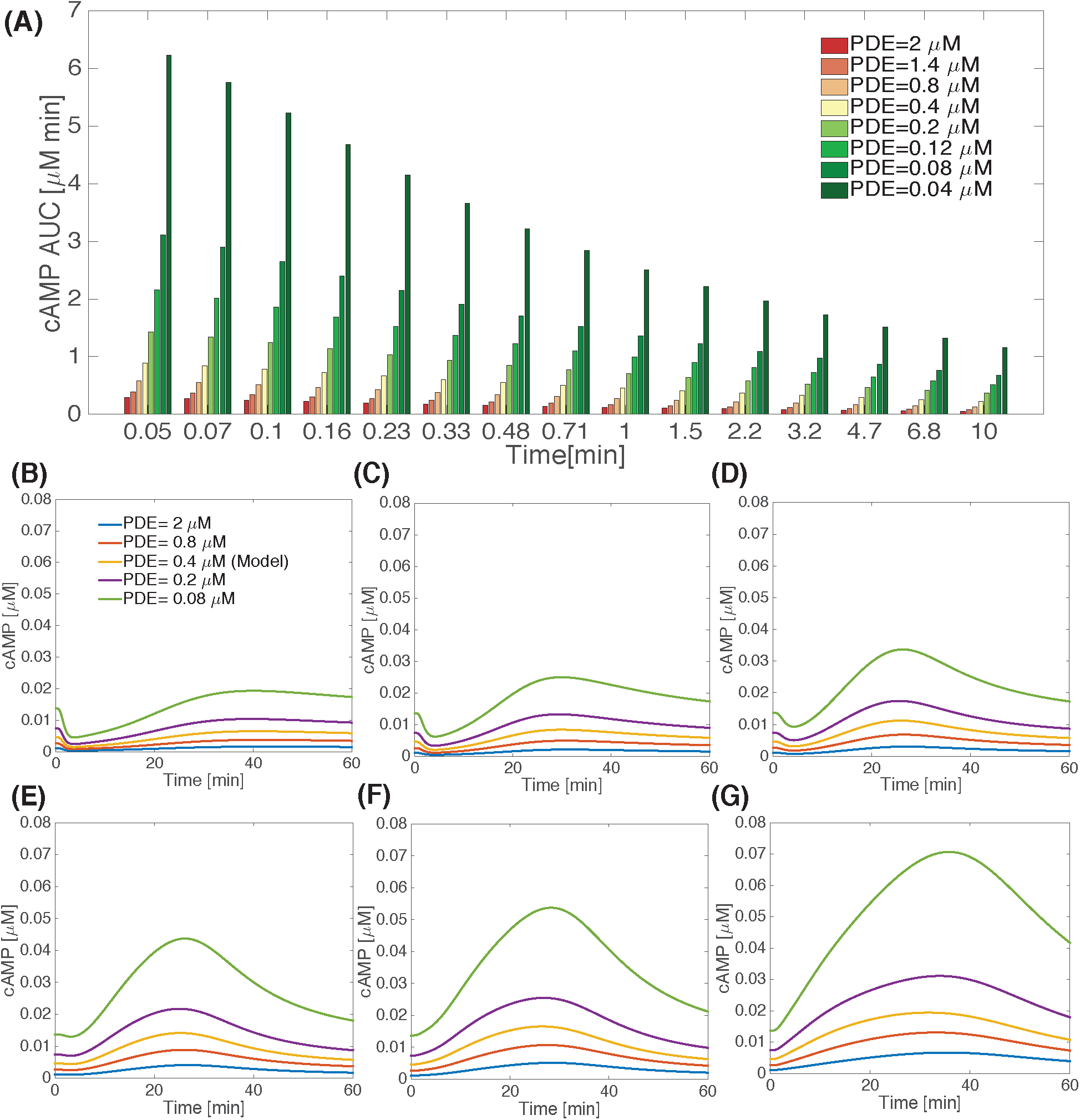
Supplementary figure for Figure 6 – Impact of varying PDE and GIV concentrations on cellular levels of cAMP. (**A**) cAMP AUC, computed at 1 h is shown for different values of GIV and PDE. Time-course of cAMP for various PDE concentrations at (**B**) GIV=0.05 *μM*, (**C**) GIV=0.2 *μM*, (**D**) GIV=0.5 *μM*, (**E**) GIV=1 *μM*, (**F**) GIV=2 *μM*, (**G**) GIV=5 *μM* are shown.

**Figure S13:**
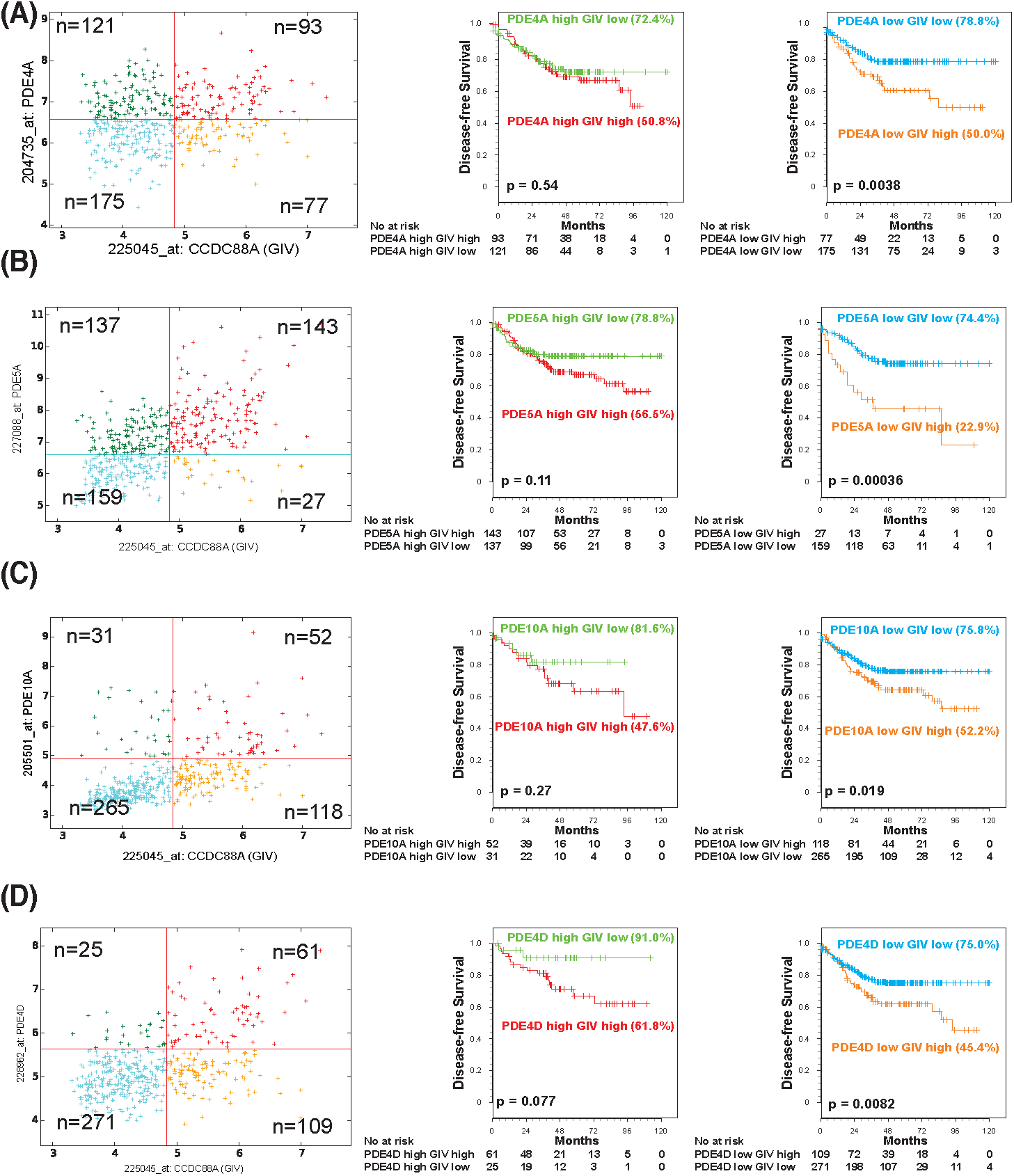
Supplementary figure for Figure 6 — The impact of levels of expression of GIV and PDE on cAMP dynamics; comparison of model predictions and clinical outcome [disease-free survival] in patients with colorectal cancers. (**A**-**D**) GIV expression status in colon cancers has an impact on disease-free survival (DFS) only when the level of expression of various PDE isoforms are low. Hegemon software was used to analyze individual arrays according to the expression levels of GIV (CCDC88A) and either PDE4A (**A**), or 5A (**B**), 10A (**C**), 4D (**D**) in a data set containing 466 patients with colon cancer (Left panels, **A**-**D**; see Methods; **E**). Survival analysis using Kaplan-Meier curves showed that among patients with high PDEs [middle panels, **A**-**D**], high vs low GIV expression did not carry any statistically significant difference in DFS (all p values > 0.05). Survival analysis among patients with low PDEs [right panels, **A**-**D**] showed that patients whose tumors had high levels of expression of GIV had a significantly shorter DFS than those with tumors expressing low levels of GIV (all p values < 0.05). See also **Figure 6** for patient survival curves for PDE5A isoform and GIV on DFS.

**Figure S14:**
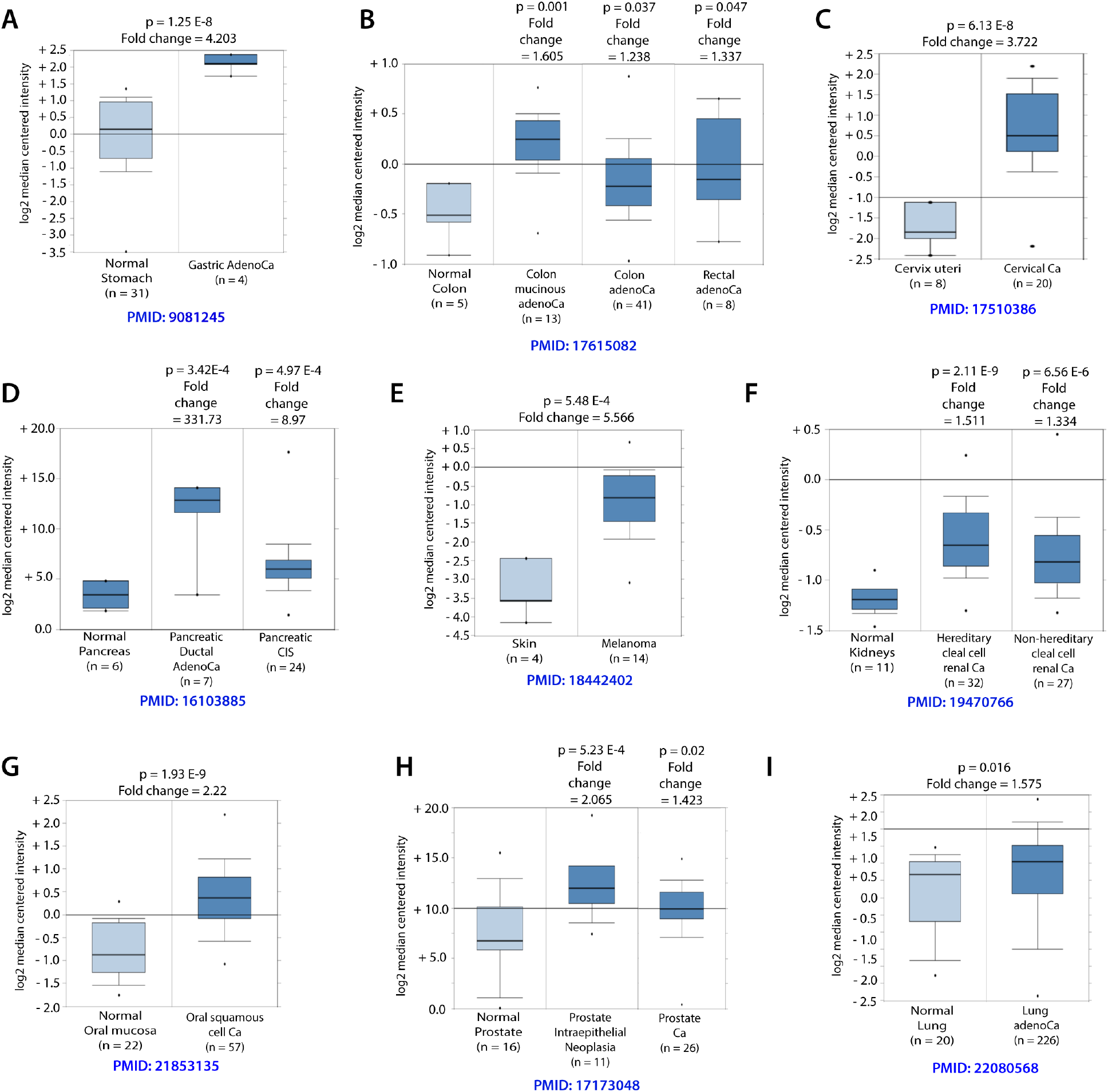
Supplementary figure for Figure 7 – GIV mRNA expression is elevated in various cancers. Expression levels of GIV [CCDC88a] mRNA in normal vs. cancers was analyzed in publicly available RNA Seq datasets using Oncomine.org. PMIDs listed under each box plot refers to the original manuscript associated with the dataset.

**Figure S15:**
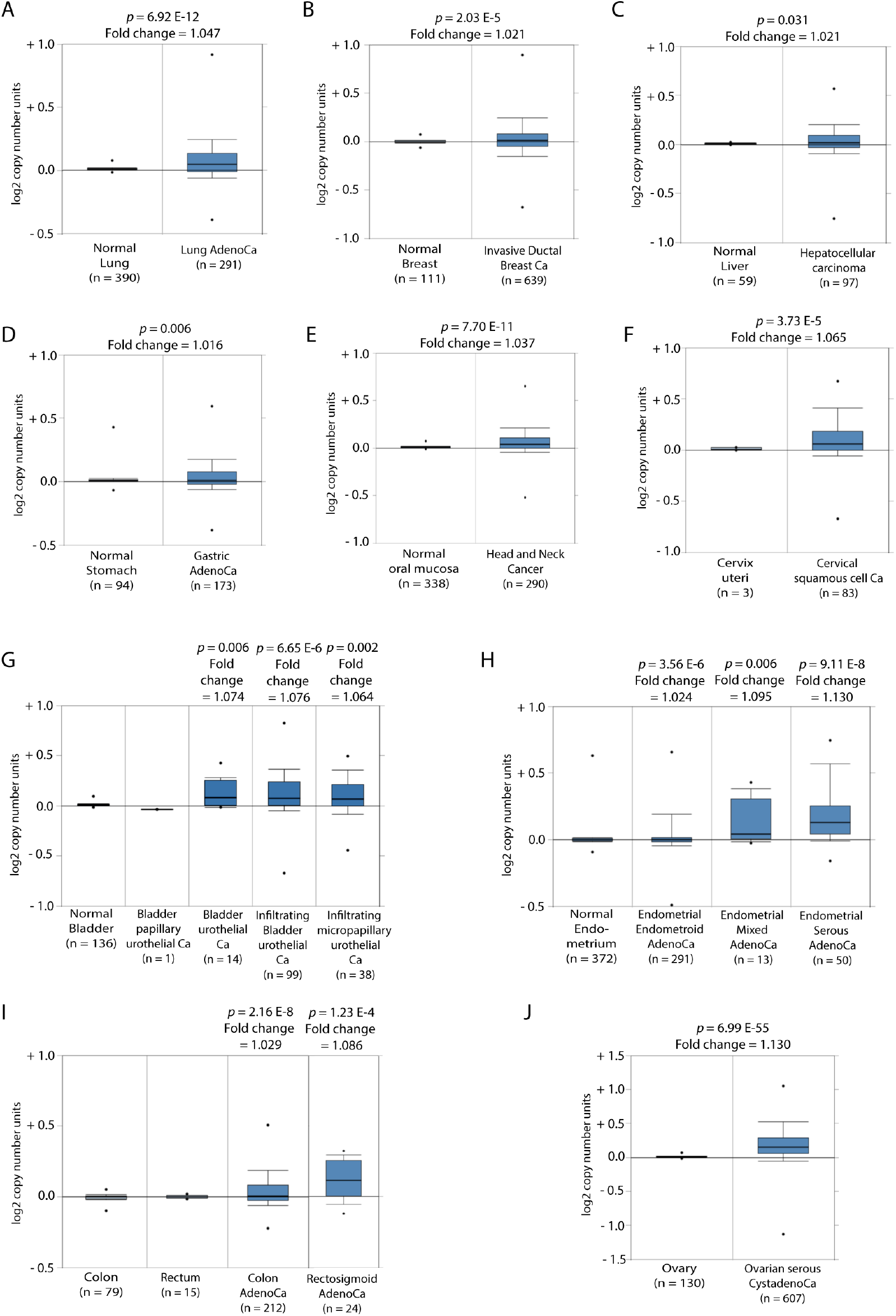
Supplementary figure for Figure 7 – Copy numbers of GIV-gene is elevated in various cancers. Copy numbers of GIV gene is elevated in various cancers. TCGA datasets were analyzed for copy number variations (CNV) in GIV gene [CCDC88a] in normal vs. cancers using Oncomine.org.

## Abbreviations used

cAMP: Cyclic adenosine monophosphate
RTK: Receptor Tyrosine Kinase
EGFR: Epidermal growth factor receptor
GEM: G protein exchange modulator
GEF: Guanine nucleotide exchange factor
GDI: Guanine nucleotide dissociation inhibitor
SH2: Src homology 2
PM: plasma membrane
DAG: Diacylglycerol
EGF: Epidermal growth factor
AUC: Area under the curve
AC: Adenylyl cyclase
PDE: Phosphodiesterase
AMP: adenosine monophosptate
ATP: adenosine triphosphate
CDK5: cyclin dependent kinase 5
PKC-*θ*: Protein kinase C *θ*
PLC-*γ*: Phospholipase C *γ*
Epac1: Exchange factor directly activated by cAMP 1
RIA: Radioimmunoassay
IBMX: 3-isobutyl-1-methylxanthene

